# Discovering host protein interactions specific for SARS-CoV-2 RNA genome

**DOI:** 10.1101/2022.07.18.499583

**Authors:** Roberto Giambruno, Elsa Zacco, Camilla Ugolini, Andrea Vandelli, Logan Mulroney, Manfredi D’Onghia, Bianca Giuliani, Elena Criscuolo, Matteo Castelli, Nicola Clementi, Massimo Clementi, Nicasio Mancini, Tiziana Bonaldi, Stefano Gustincich, Tommaso Leonardi, Gian Gaetano Tartaglia, Francesco Nicassio

**Affiliations:** Center for Genomic Science of IIT@SEMM, Fondazione Istituto Italiano di Tecnologia, 20139 Milano, Italy; Institute of Biomedical Technologies, National Research Council, 20090 Segrate, Italy; Center for Human Technologies, Istituto Italiano di Tecnologia, Genoa, Italy; Department of Biochemistry and Molecular Biology, Universitat Autònoma de Barcelona, Bellaterra, 08193 Barcelona, Spain; Universitat Pompeu Fabra (UPF), 08003 Barcelona, Spain; European Molecular Biology Laboratory, European Bioinformatics Institute, Hinxton, UK; Laboratory of Microbiology and Virology, Vita-Salute San Raffaele University; via Olgettina 58, 20132 Milan, Italy; Laboratory of Medical Microbiology and Virology, IRCCS San Raffaele Scientific Institute; via Olgettina 60, 20132 Milan, Italy; Department of Experimental Oncology, European Institute of Oncology IRCCS, 20139 Milano, Italy; Department of Oncology and Hematology-Oncology, University of Milan, Via Festa del Perdono 7, 20122 Milano, Italy; Catalan Institution for Research and Advanced Studies, ICREA, Passeig Lluís Companys 23, 08010, Barcelona, Spain

## Abstract

SARS-CoV-2, a positive single-stranded RNA virus, interacts with host cell proteins throughout its life cycle. These interactions are necessary for the host to recognize and hinder the replication of SARS-CoV-2. For the virus, to translate, transcribe and replicate its genetic material. However, many details of these interactions are still missing. We focused on the proteins binding to the highly structured 5’ and 3’ end regions of SARS-CoV-2 RNA that were predicted by the *cat*RAPID algorithm to attract numerous proteins, exploiting RNA-Protein Interaction Detection coupled with Mass Spectrometry (RaPID-MS) technology. The validated interactors, which agreed with our predictions, include pseudouridine synthase PUS7 that binds to both ends of the viral RNA. Nanopore direct-RNA sequencing confirmed that the RNA virus is heavily modified, and PUS7 consensus regions were found in both SARS-CoV-2 RNA end regions. Notably, a modified site was detected in the viral Transcription Regulatory Sequence - Leader (TRS-L) and can influence the viral RNA structure and interaction propensity. Overall, our data map host protein interactions within SARS-CoV-2 UTR regions, pinpointing to a potential role of pseudouridine synthases and post-transcriptional modifications in the viral life cycle. These findings contribute to understanding virus-host dynamics and may guide the development of targeted therapies.

## INTRODUCTION

The outbreak of severe acute respiratory syndrome coronavirus 2 (SARS-CoV-2) in the human population has dramatically affected life expectancy worldwide. SARS-CoV-2 infection causes the coronavirus disease 2019 (COVID-19), a potentially fatal condition especially for the elderly population and for individuals with underlying health issues. To date, vaccination is the most effective strategy to significantly reduce the incidence of SARS-CoV-2 infection and the onset of severe COVID-19. However, new and fast-spreading SARS-CoV-2 variants might be able to escape the antibody response of vaccinated people (1).

SARS-CoV-2 is a positive-sense single-stranded RNA virus of the *Coronoviridae* family (2, 3). Its genome is composed of ∼30.000 nucleotides and contains 14 open reading frames (ORFs) encoding 16 non-structural proteins (nsp 1 to 16), 12 accessory proteins (ORF3a, ORF3b, ORF3c, ORF3d, ORF6, ORF7a, ORF7b, ORF8, ORF9b, ORF9c, ORF9d and ORF10) and 4 structural proteins: spike (S), envelope (E), membrane (M) and nucleocapsid (N) (4–6). Structural and accessory proteins are encoded in subgenomic RNAs (sgRNAs) that bear a common 5′ leader sequence fused to different body sequences upstream to the encoded ORF (7). During its life cycle, SARS-CoV-2 RNAs interact with several host proteins specifically required for the translation of viral proteins, the transcription of the subgenomic RNAs, the replication of the genome and the generation of new viral particles (8). At the same time, host proteins recognize the presence of the SARS-CoV-2 RNA in the cytoplasm, activating the innate immune response through the interferon signalling pathways (9–11). Although the interaction between viral RNAs and host proteins has been recognized as a key molecular process for virulence, a complete and clear understanding of these interactions and their underlying mechanisms has not yet been achieved. Hence, our understanding on how viral RNA exploits cellular mechanisms for its own advantage is still limited.

Several laboratories have independently mapped the SARS-CoV-2 RNA–host protein interactome in mammalian cell lines that are permissive to SARS-CoV-2 infection (12–15). In these studies, the interactome was analysed using an RNA-centric approach to specifically purify the viral RNA at 8h-24h post-infection, a time frame in which the virus is actively replicating and produces viral proteins that hijack the host innate immune response (16, 17). Many host RNA-binding proteins (RBPs) have been reported to interact with the SARS-CoV-2 RNA. These RBPs are involved in multiple biological processes such as RNA splicing, RNA metabolism, nonsense-mediated decay, translation and viral processing. However, the interactions between host proteins and viral RNAs at earlier stages of the viral life cycle and the RNA regions specifically bound by the host proteins remain largely undetected. With this work, we aim to fill these gaps by expressing defined SARS-CoV-2 RNA fragments in human living cells and analysing those interactions with the host proteins that might occur in the absence of viral proteins. In particular, we focused on the highly structured regions containing the 5′ and the 3′ ends of SARS-CoV-2, which we previously predicted to be interaction hot spots for the host proteins, as observed also for other coronaviruses (18).

To thoroughly investigate the interactome of these key SARS-CoV-2 RNA regions, we used an approach based on the combination of the *in house* algorithm *cat*RAPID and a well-established proximity-ligation approach (19), RNA–Protein Interaction Detection-Mass Spectrometry (RaPID-MS) (20). In addition to the host proteins already reported, we unveiled other RBPs which may have key functions in the early phases of viral infection by interacting with SARS-CoV-2 genomic RNA. Interestingly, we found also the enzyme pseudouridine synthase 7 (PUS7), an RNA modifier which catalyses the pseudouridylation of RNA transcripts, highlighting a cellular mechanism in charge of introducing post-transcriptional modification on viral RNA. Nanopore direct RNA sequencing of SARS-CoV-2 RNA from infected mammalian cells confirmed the presence of many RNA modifications, including pseudouridylation sites on the viral RNAs. In particular, we found PUS7 consensus sequences heavily modified in the SARS-CoV-2 subgenomic RNAs, with one present in the stem loop 2 of the SARS-CoV-2 5′UTR, within the Transcription Regulatory Sequence - Leader (TRS-L). Biochemical and biophysical assays support a role for this unique modification by influencing both RNA structure and interaction propensity. In conclusion, we propose that the interaction and the activity of cellular pseudouridine synthases may represent a new mechanism able to influence SARS-CoV-2 life cycle.

## MATERIALS AND METHODS

### Preliminary predictions of protein-RNA interactions

The c*at*RAPID algorithm (21, 22) was used to identify the binding propensity of the 10 SARS-CoV-2 500 bp fragments against a library of 2064 human RBPs. For each fragment, only interactions with associated propensity score > 85th percentile of the fragment score distribution were retained and subsequently, the average score was calculated.

### Reagents and plasmids

The following plasmids were purchased from Addgene and used for the RaPID-MS assay: BoxB-plasmid (Addgene #107253); BoxB-EDEN15 plasmid (#107252); BASU RaPID plasmid (#107250). The following plasmids were a gift from Paul Khavari (Addgene plasmid #107253; #107252; #107250. http://n2t.net/addgene: 50917; 107252; 107250. RRID: Addgene_50917; Addgene 107252; Addgene 107250) (Ramanathan et al., 2018). SARS-CoV-2 RNA fragments, derived from the SARS-CoV-2 RNA sequence MN908947.3, were synthesized by GeneArt (ThermoFisher Scientific) according to the sequence reported in **Supplementary Table S1**, flanked by Esp3I restriction sites. The same strategy was adopted for the Scramble control sequence: 3′-CGTCTCCGCTTTCGACGACAATTTATAAAGACAGCGGTCGAGGGAAGATTTACGAGTTGAATCGAGATGCGCTGATTCGACGCAGTGTCGCGTTGTGGTGAGGTAAATTGATAGGTGTATTTTGCGAGATACAGTGATGAACACTTCATTAACAACATGATTTATACGACGATTACTAGAATTATGAAAAATGAGTCATCTACAAGCGCGTTTTTACATTGCCGTGGTTAATCGTAAGGATAGCACAGTTAACAGCGGACCCCGGCGGACTCGGCCCTATCTGAACGAATTGAGCTCCGTTCGAAATATCTAGTGAATGACCCTCCCCACGTGCCTTGATAAGCCGTGGTATTTCGTATCATACAAGTTCCAGAAGGATGGTTCAACATAGTAGGGTACCGACTGGATAGAACAAACTACTCATGTTTTCGCCGGGGGACGAACGGTAAGCTCCGCTGGGTTGACTTCTTGACCAAAGTATTTGGGTATCCAAACAGTGCCGTTAACAGCCAAGCTAGAGACG-5′.

The RNA fragments were then cloned into the pLEX BoxB-plasmid using the Esp3I sites, as described in literature (20).

pDONR207 SARS-CoV-2 NSP1 was a gift from Fritz Roth (Addgene plasmid # 141255; http://n2t.net/addgene:141255; RRID: Addgene_141255). NSP1 coding sequence was cloned into the mammalian expression pCDNA5-FRT/TO-2xSTREP-3xHA vector, gently provided by G.Superti Furga, through the Gateway cloning system (Invitrogen).

The human coding sequence of PUS7 (variant 1, C-ter tag) cloned into the pDONOR221 vector was purchased from DNASU (HsCD00867933). The sequence was mutagenized to obtain the N-ter tag canonical variant 2 through the QuikChange II XL Site-Directed Mutagenesis Kit (Agilent, 200521) using the following DNA primers: 5’-GGGAAAAAGGCTTTGGCAAATCCAAGAAAACATTCTTGGCC-3’ and 5’-GGCCAAGAATGTTTTCTTGGATTTGCCAAAGCCTTTTTCCC-3’. The same strategy was used to generate the catalytic inactive mutant PUS7 D294A, using the following primers: 5’-ATTCTCCTACATGGGAACCAAAGCTAAAAGGGCTATAACAGTTC -3’ and 5’ - GAACTGTTATAGCCCTTTTAGCTTTGGTTCCCATGTAGGAGAAT-3’.

The resultant cDNA was sequence verified and through the Gateway cloning system cloned into the pDEST-cDNA5-FRT/TO-3*Flag-APEX2 N-term, which was a gift from Benjamin Blencowe (Addgene plasmid # 182925; http://n2t.net/addgene: 182925; RRID: Addgene_182925).

The following antibodies were used in this study: Streptavidin-HRP (Cat. #3999) from Cell Signaling Technology. Anti-Vinculin (Clone H Vin 1 0, 2 Ml; Cat. #V9131) from Merck. Anti-HA-11 epitope tag, clone 16B12 from Biolegend (Cat #901501). Anti-Flag-M2 (Merck, F3165).

### Cell lines

HEK293T cells were grown in DMEM with Glucose and L-Glutamine (Lonza, BE-12-604Q) supplemented with 10% fetal bovine serum (FBS) Tetracycline free (Euroclone, ECS01822) and 100 U/ml Penicillin and Streptomycin. CaCo-2 cell lines were cultivated in Minimum Essential Medium Eagle (MEM) (Merck, M4655) complemented with 20% FBS Tetracycline free, 2 mM L-Glutamine (Lonza, BE17605E), 1 mM Sodium Pyruvate (Lonza, BE13115E), 0.1 mM not essential amino acids (Lonza, BE13114E) and 100 U/ml Penicillin and Streptomycin (Euroclone, ECB3001D).

### Virus isolation and cell infection

SARS-CoV-2 virus was isolated from a mildly symptomatic SARS-CoV-2 infected patient, as described in Ugolini et al. (6). CaCo-2 cells were infected at 80% confluency into a 25 cm^2^ tissue culture flask with SARS-CoV-2 at 0.1 Multiplicity of infection (MOI). After 1 h adsorption at 37 °C, cells were washed with PBS, and further cultured at 37 °C for 48 h with 4% FBS. After a PBS wash, enzymatic dissociation was performed for 4–6 min at 37 °C in 1 ml TrypLE (Invitrogen), then cell pellets were washed with ice-cold PBS and lysed with 1 ml of TRIzol (Invitrogen). The samples were stored at -80 °C for subsequent RNA extraction.

### RNA extraction and nanopore direct RNA sequencing

RNA was extracted from CaCo-2 cells infected with SARS-CoV-2 using the RNeasy Mini kit (Qiagen) including an in-column DnaseI treatment step, following the manufacturer protocol. The isolated RNA was then processed by direct-RNA sequencing protocol as described in Ugolini et al. (6).

### RaPID assay

The RaPID protocol was performed as reported (20) and slightly adapted as described below. HEK293T cells were transfected with plasmid vectors expressing λN-HA-BASU and one of the BoxB-RNA fragments using Lipofectamine^TM^ 3000 transfection reagent (L3000001; ThermoFisher Scientific), according to vendor’s instructions. After 48h from transfection, medium was changed and replaced with standard growth medium complemented with 200 µM biotin (Merck, B4639-1G) for 1 hour. Cells were harvested, washed once with PBS 1x and lysed with RIPA buffer (10mM Tris-HCl pH 8.0, 1%Triton-X100, 150mM NaCl, 0.1% SDS, 0.1% NaDeoxycholate, 1mM EDTA) supplemented with 1mM 1, 4-Dithiothreitol (DTT), cOmplete^™^ Protease Inhibitor Cocktail (11697498001; Merck) and PhosSTOP^™^ (4906845001; Merck). Cell lysates were incubated for 15 minutes on ice and then centrifuged at 15000 g for 15 minutes. The supernatants containing the protein extracts were transferred into fresh 1.5 ml tubes and protein concentration was measured by Bio-Rad Protein Assay Kit using BSA as protein standard (5000002; Bio-Rad). From each sample, 3 mg of protein extract was taken and brought to the same volume (600uL) with RIPA buffer. 5% of input material was taken for further analysis and 150 μL of pre-washed Streptavidin Mag Sepharose^®^ (GE28-9857-99; Merck) were added to the remaining material. Then, samples were rocked over night at 4 °C. The following day, beads were separated from the unbound fractions and 5% of each fraction was collected in fresh tubes. Beads containing the biotinylated proteins were washed 3 times with 1 mL of Wash Buffer 1 (1% SDS supplemented with 1mM DTT, protease and phosphatase cocktail inhibitors); 1 time with Wash Buffer 2 (0.1% Na-DOC, 1% Triton X-100, 0.5M NaCl, 50mM HEPES pH 7.5, 1µM EDTA supplemented with 1mM DTT, protease and phosphatase cocktail inhibitors) and 1 time with Wash Buffer 3 (0.5% Na-DOC, 150mM NaCl, 0.5% NP-40, 10mM Tris-HCl, 1µM EDTA supplemented with 1mM DTT, protease and phosphatase cocktail inhibitors). All the washes were performed by rocking samples for 5 minutes at 4 °C. Finally, proteins were eluted with Laemmli buffer containing 100 mM DTT, boiled for 5 minutes at 95 °C and processed by *In-gel* digestion. The resultant peptide mixtures were analysed by Nano-LC-MS/MS analysis (see **Supplementary methods**).

### Data analysis of MS data

Proteins were identified and quantified using MaxQuant software v.1.6.0.16. using the Andromeda search engine (23, 24). In MaxQuant, the false discovery rate (FDR) of all peptide identifications was set to a maximum of 1%. The main search was performed with a mass tolerance of 6 ppm. Enzyme specificity was set to Trypsin/P. A maximum of 3 missed cleavages was permitted and the minimum peptide length was fixed at 7 amino acids. Carbamidomethylation of cysteines was set as a fixed modification. The 2021_01 version of the human UniProt reference proteome (UP000005640) was used for peptide identification. Proteins were profiled by quantitative label-free analysis (25), activating the label-free software MaxLFQ (23) and analysed using Perseus software (26), plotting the LFQ values in a volcano plot graph, where the proteins enriched with each SARS-CoV-2 RNA fragments were compared to the Scramble RNA control. The p-Value was calculated by Perseus using a two-tailed *t*-Test. Missing values were replaced by the minimum detection value of the matrix. Proteins found as significantly enriched also with the EDEN15 RNA were removed from the final network. The interaction network generated with the statistically significant proteins was visualized using Cytoscape 3.8.1 (27).

### Computational evaluation of experimental interactions

The *cat*RAPID algorithm was used to assess the predictive accuracy on the experimental RAPID dataset. Here, for each SARS-CoV-2 fragment-human protein pair, a fragmentation procedure was performed to identify the binding regions. The propensity score is calculated taking into consideration the maximum and the minimum score of all the sub-fragments computed by the algorithm (21) and computing their difference (28). The median LFQ value for each protein-fragment pair was then normalized taking into account protein abundance in HEK293 cell lines from PAXdb (29) and protein length from UniProt database (30), using the following equation:

Normalized LFQ value = LFQ – 0.6*log_2_(abundance)-0.0001*log_2_(length).

This approach was introduced to reduce the experimental bias in LFQ scores with high protein abundance and length.

### RNA folding analysis

Predictions of RNA folding were performed using the RNAfold algorithm hosted in ViennaRNA Web Services (31) using the default parameters and displaying the centroid secondary structures.

### Gene Ontology (GO) Analysis

The GO analysis was performed on the 73 proteins specifically associated with the 10 SARS-CoV-2 RNA fragments and identified by RaPID-MS using the GOTERM biological process (BP) and molecular function (MF) present in the DAVID 6.8 Bioinformatics Resources (https://david-d.ncifcrf.gov/home.jsp) (32).

### RNA secondary structure investigation

Circular dichroism analyses were performed on the RNA oligonucleotide “SL2” of sequence 5’ -AACCAACUUUCGAUCUCUUGUAGAUCUGUUCU -3’ synthesized by Eurofins, and on the pseudouridylated SL2 sequence 5’ - AACCAACUUUCGAUCUCUUG(psi)UAGAUCUGUUCU -3’, synthesized by Tebu-bio. The two versions of SL2 were prepared in a water stock at 100 µM and diluted to a concentration of 20 µM in a buffer containing 20 mM potassium phosphate at pH 7.2 and 150 mM KF. The RNA was then incubated for 1 hour at 30 °C to allow correct folding. Data acquisition of near-UV CD spectra was performed in one-millimetre path-length quartz cuvettes on a JASCO-1100 spectropolarimeter supplied with a constant N_2_ flush at 3.0 L/min. The experiment was performed in triplicates.

### Nanopore DRS and Nanocompore analysis

The nine DRS datasets were compared to the IVT dataset, using the NRCeq assembly and the methods in (6). The dataset of Vero E6 cells, Calu-3 and CaCo-2 sample 1 and 2 are derived from (6) (ENA ID: PRJEB48830); CaCo-2 sample 3 and 4 are deposited with this manuscript (ENA ID: PRJEB53497). Reads were grouped by cell type and their fastq files concatenated.

Reads were resquiggled using F5C (v0.6) (33). IVT reads were separately mapped to each reference sgRNA using minimap2 (v2.17-r974-dirty) (34) with the following parameters: -ax map-ont -p 0 -N 10.

Nanocompore (v1.0.4) (35) was used to detect RNA modifications by comparing the IVT reads to a set of reads from each cell line for every reference sgRNA (**Suppl. Table 5, 6, 7**) with the subsequent parameters:

--fasta <NRCEQ_ASSEMBLY_FASTA_FILE> --overwrite --downsample_high_coverage 5000 -- allow_warnings --pvalue_thr 0.01 --min_coverage 30 --logit --nthreads 3 --bed <NRCEQ_ASSEMBLY_BED_FILE>.

### Merging analysis

Individual significant k-mers from the different cell lines may be offset by one or two positions, therefore the significant k-mers were transformed into larger nucleotide ranges of nine nucleotides, that include the five nucleotides of the significant k-mer, plus the two neighbouring nucleotides present at both its extremities. Any of these nine nucleotides may be a post-transcriptional modified nucleotide detected by Nanocompore. In addition, we minimized ambiguous mapping by counting significant k-mers found on any isoform of VME1 and NCAP as a single isoform according to their genomic coordinates. Sites were compared to the “high-confidence sites” defined by Fleming et al. (36) using the single-nucleotide genomic position. Sites were manually inspected by plotting the distribution of signal intensity and dwell time for each position using functions implemented in Nanocompore (35).

### Comparison with RaPID fragments

Both significant k-mers and sites were compared with the RaPID fragments. Genomic coordinates were used for all comparisons with RaPID fragments (**Suppl. Table 1**). In the case of k-mers, we defined a RaPID fragment match when the first nucleotide of the k-mer overlapped with any RaPID fragment, while for sites, we searched for every nucleotide of the site that could overlap any fragment. Genomic regions overlapping two fragments were identified with both fragment names. K-mers were assigned to each fragment based on the genomic position of their first nucleotide.

### Genomic RNAs analysis

All infected datasets were mapped to the viral genome reference with minimap2 with the subsequent parameters:

-k 8 -w 1 -ax splice -g 30000 -G 30000 -A1 -B2 -O2, 24 -E1, 0 -C0 -z 400, 200 --no-end-flt -F 40000 -N 32 --splice-flank=no --max-chain-skip=40 -un -p 0.7.

All the positive strand primary alignments, with no deletions, ranging from at least 45 and 29850 in genomic coordinates were extracted from all the datasets. These reads were compared to the IVT reads, pre-aligned to the viral reference genome, through Nanocompore with the subsequent parameters:

--downsample_high_coverage 2100 --allow_warnings --min_coverage 10 --logit.

The downsampling parameter was set to 2100 as it was obtained from the multiplication of the number of gRNA reads (140) for the number of IVT fragments (15), as downsampling is performed over all the dataset. K-mers were evaluated nucleotide per nucleotide, to assess their overlap with IVT junctions, RaPID fragments, Fleming et al. sites, sgRNAs sites from the previous analysis and Single Nucleotide Polymorphisms (SNPs). The list of SNPs collects data from the literature (see **Supplementary Table 5**). Only k-mers not overlapping any SNP or IVT junction were considered. K-mers were considered significant if having an absolute LOR >=0.5 and a GMM p-value <=0.01. We then investigated if, despite the low coverage noticed for gRNAs, a comparison between the sgRNAs-specific sites and the gRNAs-specific ones could be performed. Therefore, to make the comparison between the gRNAs and the sgRNAs datasets fair, we pulled together all the data from each cell line analysed and selected sgRNAs uridine-containing significant k-mers present in at least one canonical transcript model (see **Supplementary Table 10**). To determine if the common sites were randomly selected or if such a number was to be statistically expected, we performed a hypergeometric test on common sites located after the TRS-B spike junction sequence, both for gRNAs and sgRNAs. The significance of the test (p-value = 1.44e-15, see github directory) suggests that, if we increased the coverage of the gRNAs dataset, we might retrieve the same sites found for the sgRNAs in the region starting from the TRS-B spike junction sequence to the end of the viral genome. The test could not be performed on the region starting at the genomic zero coordinate to the TRS-B spike junction sequence, as, of course, sgRNAs do not cover this portion of the genome.

## RESULTS

### Prediction of SARS-CoV-2 regions with the highest propensity to interact with host proteins

We previously reported that RNA regions with high structural content tend to tightly interact with a large number of proteins (37, 38), which suggests that structured regions of SARS-CoV-2 RNA genome might play a crucial role in its regulation (18). In line with this finding, we and others have observed that the first and last 1.5 kb of the viral genome recruit specific factors required for viral translation and replication, and that they act as targets for the intracellular host defence response (18, 39). Despite previous attempts, a comprehensive and reproducible map of RNA:host protein interactions is missing (40). We decided to accurately map the regions with the highest propensity to interact with host proteins and, accordingly, divided the initial as well as terminal 1.5 kb regions of the viral genome into 500 nucleotide-long RNA fragments. Each fragment overlaps the following one by 250 nucleotides. A total of five fragments for the 5′ region (numbered 1 to 5) and five for the 3′ region (numbered 6 to 10) was identified (**Supplementary Table 1** and **Fig. 1A**). Through the *cat*RAPID algorithm (21), which estimates the binding affinity of protein-RNA pairs (41, 42), we predicted the interactions of each fragment with human RBPs (over 2000 entries; **Supplementary Table 2**). In this analysis we did not consider the cell-specificity pattern of expression of the RBPs, as the calculations serve to provide an estimate of the protein binding ability of RNA fragments. We identified fragments 1 and 2 and fragments 9 and 10 as those harbouring the highest scores (**Fig. 1B**). Considering that the 5′ and 3′ UTRs (fragments 1 and 10) have the strongest scores, we reasoned that these regions might also be the most functionally relevant (43). Altogether, these results provide important indications on the location of structural hot spots potentially relevant for the functional interactions between the virus and the host.

**Figure 1.**
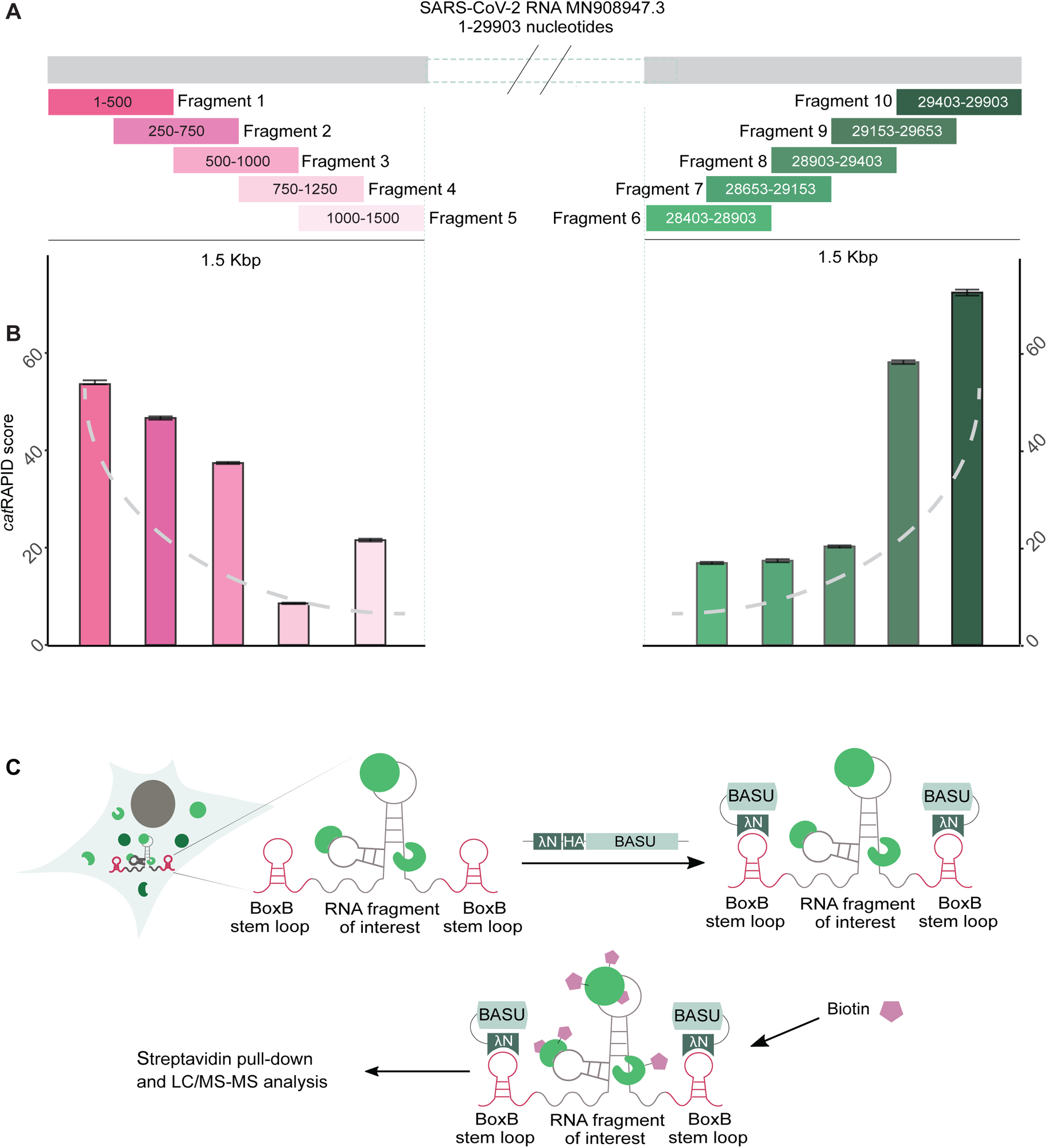
Experimental approach to investigate SARS-CoV-2 interactome. **A)** Schematic representation of the SARS-CoV-2 RNA fragments selected to be studied with the RaPID-MS strategy. The scheme shows the fragments names, positions within the SARS-CoV-2 genome and degree of overlap between them. **B)** *cat*RAPID predictions of the 10 selected RNA fragments with the catalogue of human RNA-binding proteins. Fragments belonging to the first and last 1.5 Kbp of SARS-CoV-2 genome are coloured in different shades of pink and green, respectively. Error bars for each fragment correspond to the average value ± the standard error (SE). The grey dashed line indicates the trend of the *cat*RAPID score in the chart. **C)** Description of the technique RaPID-MS, in which the RNA fragment of interest is expressed in cells flanked by BoxB stem loops. BoxB is specifically recognized by the co-transfected λN peptide fused to the biotin ligase BASU. Upon biotin addition to the growth medium, BASU biotinylates the host protein interactors attracted by the RNA of interest.

### Identification of the human interactome for the 5′ and 3′ ends of SARS-CoV-2 genome

To analyse the RNA interaction with the host proteins experimentally, we exploited the RaPID-MS approach (20) using the 10 above-described fragments generated by *cat*RAPID (**Fig. 1A and Supplementary Fig. 1**). The fragments were cloned within BoxB sequences (**Materials and Methods**), used as tags for the RaPID assay, and were expressed into HEK293T cells as host cell line (**Fig. 1C**). As a negative control, we generated a scrambled sequence of 500 nt of similar GC content compared to the other fragments (named ‘Scramble’ henceforth). The BoxB-tagged EDEN15 RNA was selected as positive control (20). We verified that: i) each plasmid expressing an RNA fragment was co-expressed with a plasmid coding for the biotin protein ligase λN-HA-BASU by FACS analysis (**Supplementary Fig. 2)**; ii) λN-HA-BASU was expressed correctly and able to biotinylate protein substrates in the presence of exogenous biotin by Western Blot (WB) analysis (**Supplementary Fig. 3A**). The resulting biotinylated proteins were purified under denaturing conditions using streptavidin beads, as confirmed by WB analysis (**Supplementary Fig. 3B**) and analysed by liquid chromatography tandem mass spectrometry (LC-MS/MS). In total, 3 independent biological replicates of RaPID-MS were performed using the 10 SARS-CoV-2 fragments, as well as the Scramble and the EDEN15 control RNAs. We identified a total of 1296 proteins interacting with our selected fragments, a resource reported in **Supplementary Table 3**. A comparison between our dataset and the ones generated by previous studies indicates an overlap of about 40% (**Supplementary Table 3)**, demonstrating good agreement with literature (44, 45).

As we are interested in finding host proteins that specifically bind to SARS-CoV-2 regions rather than other RNAs, we compared the list of interactors obtained with each individual fragment with those obtained with our negative control Scramble (**Supplementary Fig. 4A and 4B, and Supplementary Table 3**). We removed RBPs that were found as interacting to the same extent of Scramble (**Supplementary Table 3)**, including PTBP1, G3BP1 and SYNCRIP, which were previously reported to specifically interact with the SARS-CoV-2 RNA (12–15). We were able to verify the specificity of our approach, as we identified CELF1 as the interactor of our positive control EDEN15 (**Supplementary Fig. 4B**), as expected (46). In total, we isolated 73 proteins significantly and specifically interacting with SARS-CoV-2 RNA fragments, hereafter called ‘RaPID-MS dataset’ (**Fig. 2**). In the RaPID-MS dataset, we have 10 proteins, previously reported as interactors of SARS-CoV-2 RNA, namely FAM120A, HAT1, LSG1, SHMT1, SYNE2 and the ribosomal proteins RPL14, RPL18A, RPL24, RPL35, RPS6. The remaining proteins were identified for the first time (**Supplementary Fig. 5A and 5B)**. The highest number of interacting proteins were in fragments 1, 4, 7 and 10. Conversely, fragment 2 displayed only two specific interactions (**Fig. 2A**).

**Figure 2.**
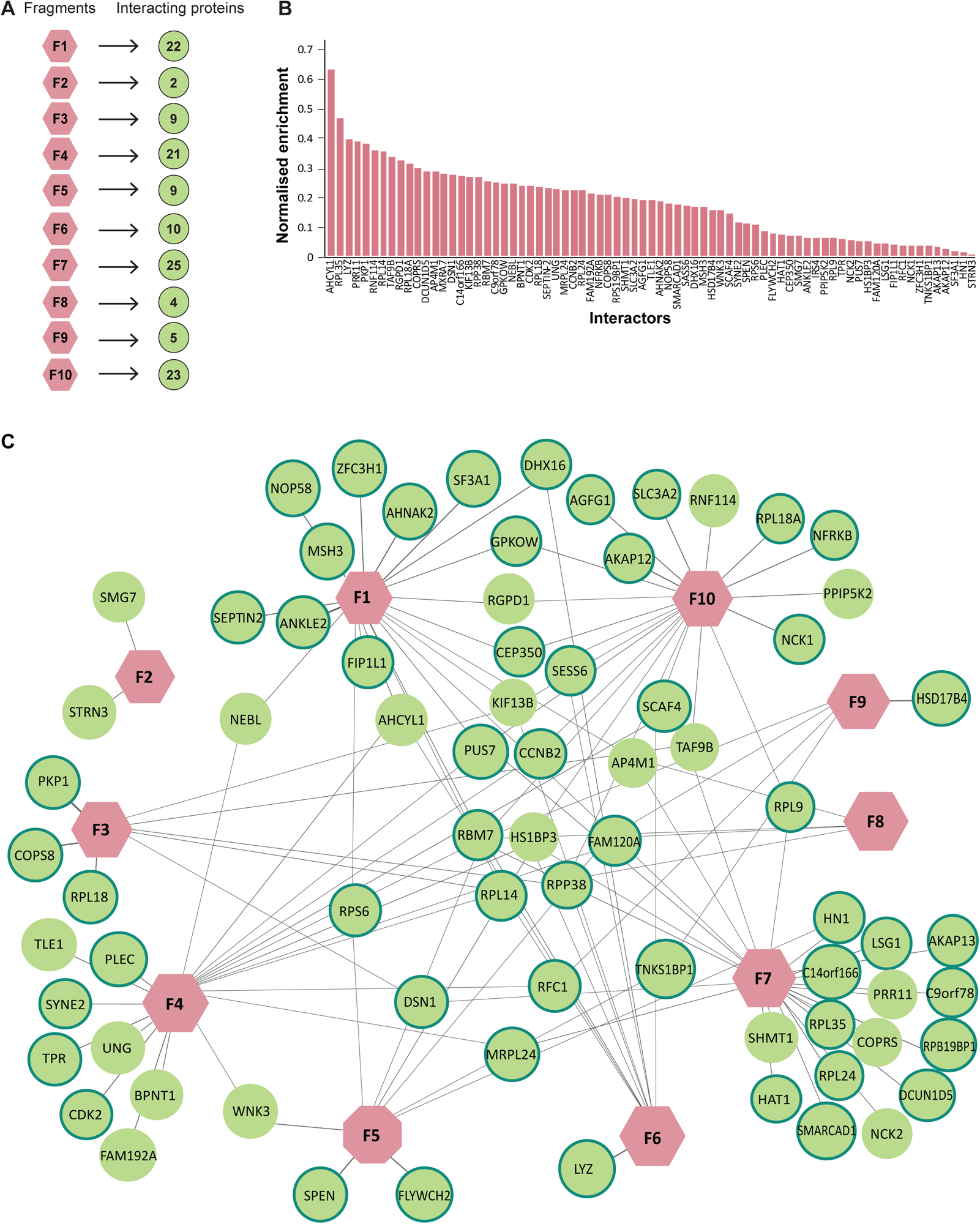
Defining SARS-CoV-2 interaction network with the human proteome. **A)** Number of specific interactors identified for each SARS-CoV-2 RNA fragment (labelled with the letter “F” follow by a number). **B)** Enrichment distribution of the specific interactors for all considered fragments compared to the control “Scramble”. Data are normalized for protein length and abundance, as specified in the Materials and Methods section. **C)** The network of proteins identified by RaPID-MS. Only proteins significantly enriched over the control “Scramble” are displayed. Data are derived from the analysis of 3 independent biological replicates. In pink are displayed the RNA fragments, while in green the retrieved interactors. Proteins circled in green are RNA-binding proteins, according to the RBPome database (https://rbpbase.shiny.embl.de/).

**Figure 3.**
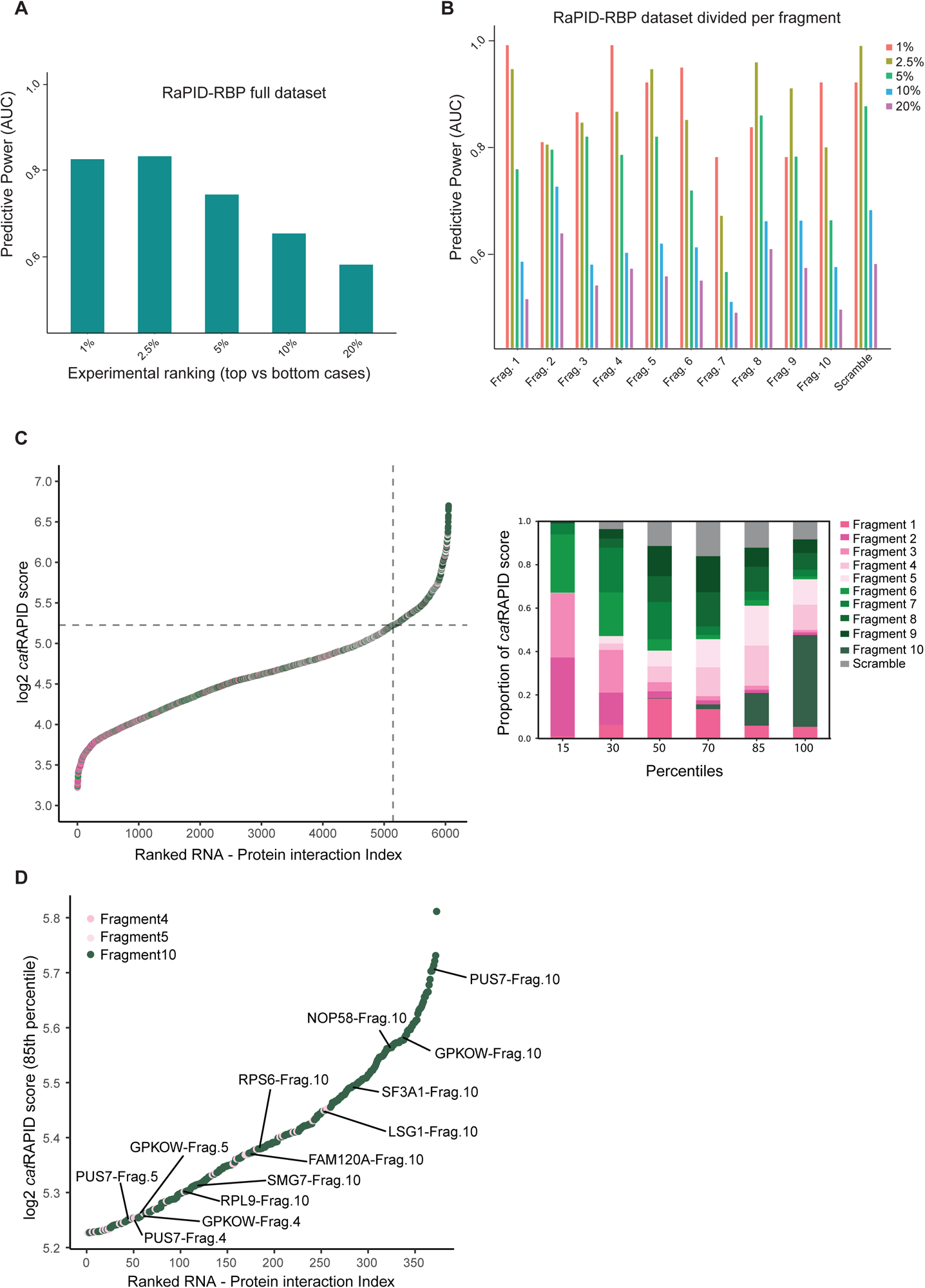
*cat*RAPID performances on the RaPID-RBP datasets. **A)** *cat*RAPID performances on the RaPID-RBP dataset. The predictive power of the method is calculated for different percentages of the dataset. *cat*RAPID performance is calculated on LFQ experimental value, normalized by taking into account the abundance of the proteins in PAXdb and protein length in Uniprot. **B)** *cat*RAPID performance on the RaPID-RBP dataset, focusing on the single RNA fragments. For each fragment, the AUC at the different percentage of the dataset is shown. The performances are evaluated as in **A** and the fragments are ordered according to the respective genomic position. **C) Left.** Scatter chart of the analyzed RBPs ranked according to their relative log_2_ *cat*RAPID score. The dashed lines indicate the 85th percentile. **Right.** Boxplot representation of the whole distribution divided in the indicated percentiles. **D)** Scatter chart of the analysed RBPs ranked according to their relative log2 *cat*RAPID score, displaying only proteins present in the 85th percentile of the distribution.

**Figure 4.**
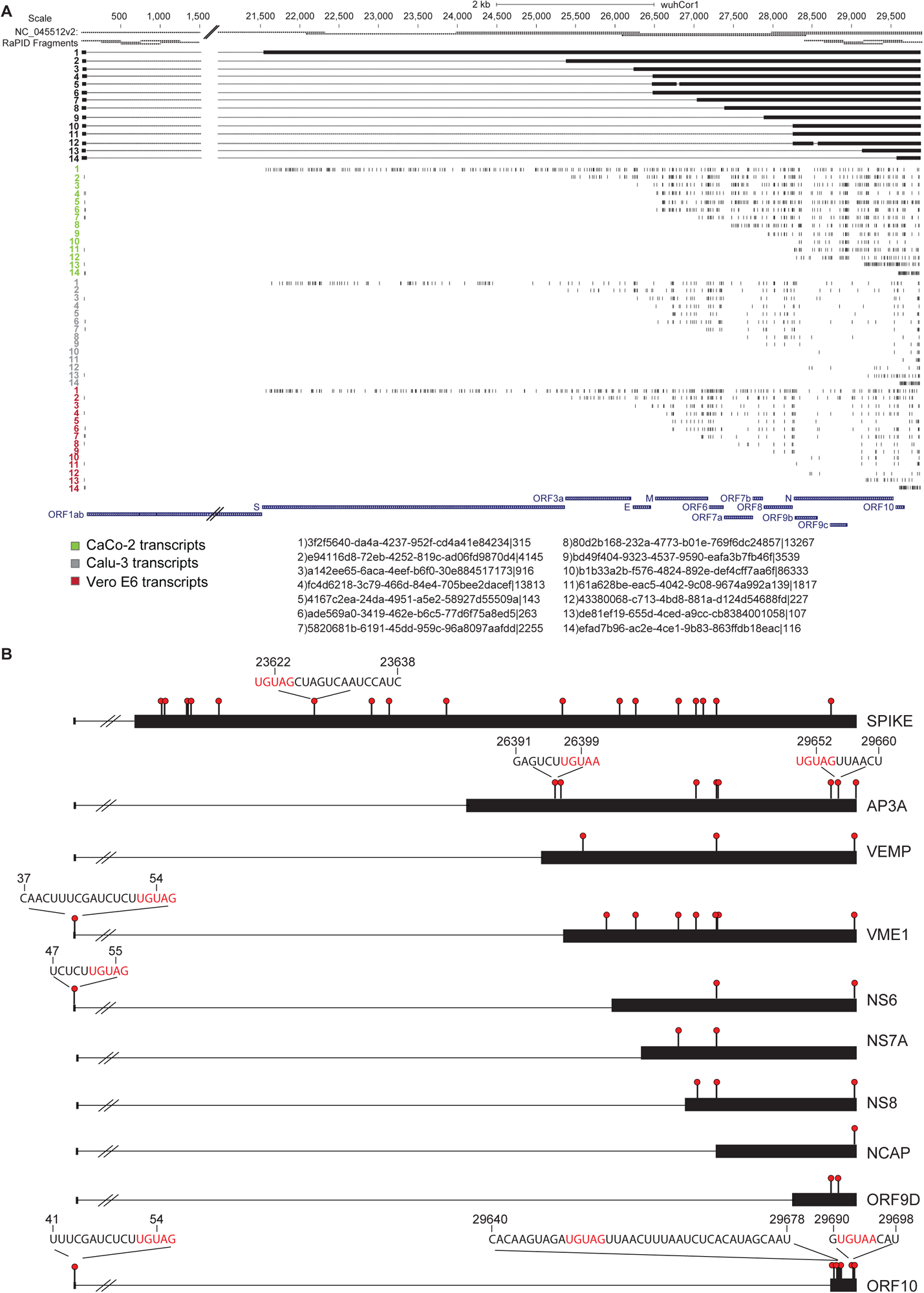
Nanocompore analysis identified significant k-mers in the analysed cell lines. **A)** UCSC Genome Browser (http://genome.ucsc.edu) annotation of the modified k-mers containing a uridine found in SARS-CoV-2 infected CaCo-2, Calu-3 and Vero E6 cells and distributed over 14 different reference sgRNAs. From the top to the bottom, IVT and RaPID fragments, SARS-CoV-2 NRCeq assembly and RefSeq SARS-CoV-2 ORFs are present as a reference. **B)** Graphical representation of nucleotide ranges shared between at least two cell lines (red circles). Here, sites have been grouped per ORFs encoded by each reference sgRNA of the assembly and the sequence of those carrying an UGUAR motif is displayed.

**Figure 5.**
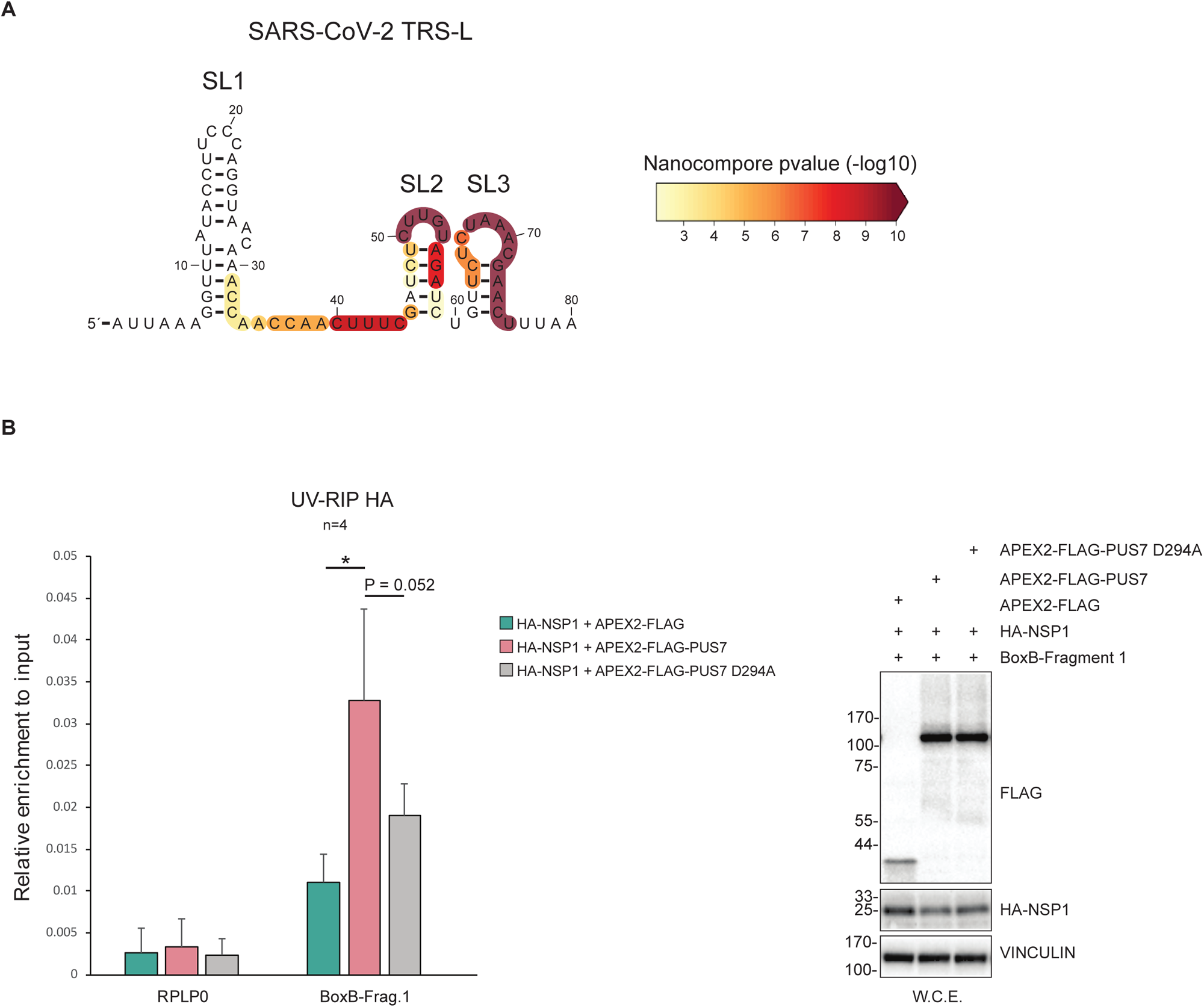
PUS7 modifies the SL2 of SARS-CoV-2 TRS-L favouring the binding of NSP1. **A)** The secondary structure of SARS-CoV-2 TRS-L with the Nanocompore p-value (NS6 sgRNA vs IVT, GMM-logit test) overlaid as a colour scale. The colour represents the lowest p-value among those of the 5 overlapping k-mers. Only k-mers with a *p*-value ≤ 0.01 are coloured according to the relative scale. **B) Left.** UV-RIP of HA-NSP1 co-expressed in HEK293 cells together with BoxB-Frag.1 expressing vector and either APEX2-FLAG, APEX2-FLAG-PUS7 or APEX2-FLAG-PUS7 D294A. The RNA bound to HA-NSP1 was eluted, retrotranscribed into cDNA and analysed by RT-qPCR analysis using specific primers for RPLP0 and BoxB-Frag.1. The data shown are the average of 4 biological replicates (n =4) and are expressed as relative enrichment over each respective input. Statistical analysis was performed using non-parametric, 2-tailed *t*-Test. *= Values with a P value < 0.05. **Right.** Representative image of the four WB analysis conducted on an aliquot of the UV-RIP input material (10%), showing the expression levels of the exogenous protein transiently co-expressed in HEK293 cells.

To balance the possible bias that could be introduced by the RaPID technique, we considered protein length and abundance to compute the median enrichment value of each significant interactor (**Fig. 2B**). As for other studies of SARS-CoV-2 RNA interactome (40), we retrieved ribosomal proteins (N=7) among the most highly enriched interactors (**Fig. 2B**), explained by the fact that some of the fragments contain open reading frames (ORFs). We expected the RaPID-MS dataset to be enriched for direct RNA-protein interactions. Indeed, 73% (53/73) interactors are annotated as RBPs according to the RBPome database (https://rbpbase.shiny.embl.de/) (green circles in **Fig. 2C**), while the remaining are members of the protein-complexes biotinylated in proximity of the target. We noted that while some proteins (N=26) were shared among 2 or more fragments, most interactions (N=47) are specific for just one RNA fragment (**Fig. 2C**). In some cases, the proteins interact with the overlapping regions included within the fragments, reinforcing our conclusions. Examples are WNK3 and AP4M1, which were found in association with the neighbouring fragments 4 and 5 and fragments 7 and 8, respectively. Interestingly, three proteins (CEP350, RGPD1 and GPKOW) were exclusively interacting with the two terminal fragments (1 and 10), suggesting they might bind 5′ and 3′ ends either independently or by participating in viral RNA circularization, as recently described (47). Gene set enrichment analysis (GSEA) for gene ontology (GO) processes highlighted that the retrieved interactors belonged to cellular pathways involved in viral transcription, nonsense mediated mRNA decay (NMD), rRNA processing, translational initiation, mRNA splicing and mRNA export from the nucleus (**Supplementary Fig. 5C**).

### Correlation between computationally predicted and experimentally validated RNA-host protein interactors

To properly compare predicted and experimental interactions, the median enrichment value of each experimentally identified interactor was normalized by protein length and abundance (**Supplementary Table 4**). We restricted the computational prediction analysis to human RBPs reported in the *cat*RAPID library (21), here named RaPID-RBP (**Supplementary Table 4**), and assessed to what extent the predicted binding propensities are in agreement with the experimental results obtained in the RAPID-MS dataset. In particular, we verified whether the strongest positive (*i.e.*, interacting) and strongest negative (*i.e.*, non-interacting) predicted protein-RNA pairs could be identified in the list of experimentally validated interactions. We evaluated the performance of our prediction using the Area Under the ROC Curve (AUC). Overall, *cat*RAPID reached an AUC of over 0.80. The AUC increased from 0.58 to 0.83 when applying the predictions to the experimental scores from the top 20% (*i.e.,* the 20% strongest positives vs the 20% strongest negatives) to the top 2.5% (*i.e.,* the 2.5% strongest positives vs the 2.5% strongest negatives) (**Fig. 3A** and **Supplementary Table 4**). When considering each fragment separately, the prediction performances increased even further for the terminal regions of the viral genome (fragments 1 and 10) and for some internal overlapping fragments (fragments 4, 5, 6), reaching an AUC of 0.95 for the top 1% ranked experimental cases (**Fig. 3B and Supplementary Table 4**). Overall, this analysis suggests that predicted and the experimental data are well correlated.

Prompted by this observation, we exploited the *cat*RAPID score to produce a rank of the RBP interactome of SARS-CoV-2 RNA identified by RAPID-MS. The log_2_ *cat*RAPID score for all the possible viral RNA-host protein interactions was between 3.22 and 6.70, with a median value of 4.62 (**Fig. 3C and Supplementary Table 4**). The *cat*RAPID predicted RNA-protein interactions of each viral RNA fragment were differently distributed within these intervals (**Supplementary Fig. 6A-B**). In particular, proteins above a log_2_ *cat*RAPID score of 5.22, corresponding to the 85^th^ percentile and the graphical inflection point of the distribution, are significantly enriched for interactions with fragment 10, 4 and 5 (**Fig. 3C and Supplementary Table 4**). These three fragments together displayed 373 predicted interactions that were not found with the RNA control Scramble, including 13 interactions involving 9 RBPs significantly associated with the SARS-CoV-2 RNA fragments by RaPID-MS (**Fig. 3D**). Among them, i) FAM120A, LSG1 and RPS6 that were already reported as host proteins reproducibly interacting with SARS-CoV-2 RNA (40); ii) the splicing factors GPKOW, SF3A1, NOP58 and the NDM factor SMG7 that were computationally predicted to specifically associate to the genomic viral RNA (48, 49); iii) the RNA modifier PUS7, that catalyses the isomerization of uridine into pseudouridine in cellular tRNAs and mRNAs, reported here for the first time. In the RaPID-MS dataset, PUS7 specifically associates with fragments 1, 4, 7 and 10, all harbouring a PUS7 consensus sequence. Importantly, pseudouridine has been found as the most abundant modification in SARS-CoV-2 RNA (10) and another member of the pseudouridine synthase family, PUS1, has been reported to interact with SARS-CoV-2 RNA (14). These observations strongly suggest that, in human cells, the formation of pseudouridine residues on SARS-CoV-2 genome could be catalysed by the activity of cellular RNA-independent pseudouridine synthases, such as PUS1 and PUS7.

### SARS-CoV-2 subgenomic RNAs bear multiple putative pseudouridylated sites

To directly investigate occurrence of RNA modifications in SARS-CoV-2, including pseudouridine sites, we employed Nanopore direct RNA sequencing (DRS) on cell lines infected with SARS-CoV-2 and analysed the data with the Nanocompore software package (**Materials and Methods**). Nanocompore searches for RNA post-transcriptional modifications by comparing the ionic current features generated by Nanopore DRS from two different experimental conditions: a test sample and a reference devoid of the modification of interest (or with a reduced number of them) (35). Due to the physics of nanopore sequencing, Nanocompore compares the ionic current features of five nucleotides (named as k-mer) at a time. To identify putative pseudouridine sites, we filtered the results by selecting k-mers that contained at least one uridine and identified as significant by Nanocompore (p-value ≤ 0.01, absolute value of the log odds ratio (LOR) ≥ 0.5). Initially, we used this technology to search for pseudouridine sites in the full-length genomic RNA (gRNA). We compared gRNA reads from CaCo2, Calu-3 and Vero E6 infected with SARS-CoV-2 with a baseline reference, which consists of unmodified SARS-CoV-2 RNA obtained by in vitro transcription (IVT) (50). Nanocompore identified 63 significant k-mers (p-value ≤ 0.01, abs (LOR) ≥ 0.5), of which 58 having a uridine at the centre of the identified sequence or in the first or second neighbor nucleotide (**Fig. 4, Supplementary Fig. 7A-B and Supplementary Table 5**). Among them, 6 k-mers were located within RaPID fragments 3-4, 5, 7-8, 8-9 and 9-10 (**Supplementary Table 5** and **Supplementary Fig. 7B**). However, in the analysed samples we noticed a coverage of only 140 reads spanning the whole 30 kb-long genome. Therefore, we extended our analysis to the subgenomic RNAs (sgRNAs) that were more abundant, as previously quantified (6). We analysed all the 14 canonical sgRNAs of SARS-CoV-2 (**Fig. 4A** and **Supplementary Table 6-8**) used by the virus to translate the structural and accessory proteins required to produce new virions. In the attempt of characterizing sgRNAs, we have previously employed Nanopore ReCappable Sequencing (NRCeq), a new technique that can identify capped full-length RNAs (6). Thus, we compared viral sgRNA from infected Vero E6, CaCo-2 and Calu-3 cells to the unmodified SARS-CoV-2 IVT RNA, used as reference. Sequenced reads were aligned to the most up-to-date viral reference transcriptome (6) and the ionic current features from the RNA reads were realigned to each transcriptomic position of the reference using Nanopolish (51). Then, we used Nanocompore to identify RNA post-transcriptional modifications as marked by differences in the electrical signal between the viral sgRNA reads and the IVT RNA reads (p-value ≤ 0.01, abs (LOR) ≥ 0.5). In total, we identified 1164 (CaCo-2), 430 (Calu-3) and 627 (Vero E6) significant uridine-containing k-mers, distributed among the 14 canonical SARS-CoV-2 reference sgRNAs (**Fig. 4A, Supplementary Fig. 8** and **Tables 6-8**).

To focus on sites consistently modified across samples, we considered only significant modified uridines that were identified in at least two out of three SARS-CoV-2 infected cell lines (see **Materials and Methods**). Overall, we obtained 457 candidate modified regions across the 10 different SARS-CoV-2 sgRNAs encoding for canonical ORFs (**Fig. 4B)**. Two of these regions were recently reported as a pseudouridylated sites in SARS-CoV-2 RNA (36) (**Supplementary Table 9**). In addition, from a supplementary analysis comparing the significant uridine-modified k-mers found in the SARS-CoV-2 sgRNAs and gRNAs, we retrieved an overlap of 17 sites (**Supplementary Fig. 7C** and **Supplementary Table 5**).

To shortlist the sites that have a higher likelihood of being pseudouridines (ψ) modified by PUS7, we selected those harbouring the RNA consensus sequence of PUS7. We found that 53 sites had the more generic PUS7 consensus sequence UNUAR (red lollipops of **Fig. 4B)**, while eight had the more restrictive UGUAR motif (red lollipops with displayed sequence of **Fig. 4B)**. Six sites are located within RaPID fragments 1, 9 and 10, of which 1 and 10 were bound by PUS7 in our interactomic data (**Supplementary Table 9** and **Fig. 2C**). The modified sites containing the UGUAR consensus were manually inspected to confirm that the distributions of ionic current intensities and dwell times were different between the IVT and SARS-CoV-2 reads in a window of nine k-mers centred on the central U of the UGUAR motif (**Supplementary Fig. 9)**. Interestingly, we observed that three of these sites were present within the Transcription Regulatory Sequence - Leader (TRS-L) at the 5′ UTR of the sgRNAs encoding for ORF10, NS6, and VME1 (**Fig. 4B** and **Supplementary Table 9**).

### PUS7-dependent pseudouridylation of TRS-L could modulate the binding of NSP1 to the viral 5’UTR

In total, Nanocompore analysis of the sgRNAs identified multiple putative post-transcriptionally modified sites in the TRS-L of SARS-CoV-2 (**Fig. 5A**). Conversely, the UTR of the gRNA resulted devoid of any modification (**Supplementary Table 5**). One of the modified sites is within the m6A consensus motif DRACH, which has recently been reported to be methylated by METTL3 and to regulate the translation rate of SARS-CoV-2 proteins (52).

Another modified site is, instead, present in the PUS7 consensus sequence UGUAR, thus pointing towards a putative functional role of PUS7-dependent pseudouridylation of the SARS-CoV-2 UTR. We sought to understand the impact that the presence of a pseudouridine on the TRS-L might have. The modified site is present at the highly accessible uridine 54 (U54) within the stem loop 2 (SL2) (**Fig. 5A**), the most conserved region of the 5′UTR of Coronaviruses (53). We obtained the chemically synthesized SL2 RNA sequence either unmodified or carrying a pseudouridylated U54. Structural investigation performed via circular dichroism revealed that the presence of ψ slightly increases the propensity of SL2 to form more stable secondary structures (**Supplementary Fig. 10A**), likely affecting the conformation of the SL2 pentaloop sequence (54). Since RNA secondary structure is one of the drivers of the interactions with proteins, we reasoned that the presence of a pseudouridylated SL2 might favour its interaction with its main binder, the viral protein NSP1. NSP1 binds to the SARS-CoV-2 leader sequence to protect and promote the translation of viral RNA transcripts (8, 55, 56). By means of biolayer interferometry, we estimated the impact of this pseudouridylation on the dissociation constant (K_d_) with a recombinant NSP1. We observed that NSP1 is able to bind the *in vitro*-synthesized SL2 RNA sequence, with or without pseudouridylation, with a K_d_ in the high nanomolar range. However, the presence of ψ increased 2-fold the binding affinity, improving it from 300 nM to 170 nM (**Supplementary Fig. 10B**). The same trend of enhanced binding was assessed in cells through UV-RIP analysis performed with STREP-HA-tagged NSP1. We found that NSP1 is able to bind to the BoxB-RNA fragment 1 carrying the SARS-CoV-2 5′UTR (**Fig. 5B**). This RNA-protein interaction is enhanced by the ectopic expression of PUS7, while no differences were observed by the expression of a catalytic defective version of PUS7 (D294A) (57) (**Fig. 5B**). These results hint at a possible role of PUS7-mediated pseudouridylation of SL2, a region included in the 5′UTR that is typically bound by NSP1 to favour the translation of viral sgRNAs.

## DISCUSSION

Host-virus interactions encrypt for the multiple processes of the viral life cycle, including, but not limited to, translation, transcription, and replication (40, 44, 45). Unveiling the interactions between host proteins and both SARS-CoV-2 genomic and subgenomic RNAs holds the potential for the development of therapeutic strategies to inhibit the replication of this virus in human cells. Here, we propose an integrated approach that synergizes experimental results with computational predictions to construct a comprehensive portrait of the human proteins interacting with the SARS-CoV-2 RNA, enabling a better understanding of the molecular landscape of the virus-host interactome.

We exploited the state-of-the-art proximity ligation technology RaPID-MS (20), a technique that enables the identification of the interactions between a known RNA sequence of interest and the proteins of the selected cell line. This method provides two critical advantages over other methodologies: i) it circumvents any crosslinking step, reducing the number of proteins non-directly associated with SARS-CoV-2 RNA; ii) it allows the identification of viral RNA-host protein interactions established during the early phases of infection. This critical period is marked by a minimal expression of viral proteins and displays the dependency of the virus on the interactions with the host proteins.

Existing studies have sought to determine the network of the SARS-CoV-2 RNA interactome with human proteins, yielding to a global view at various time points post-infection (12–15). However, a significant divergence is observed across these studies as a plausible consequence of different host cells and infection times selected; as well as of technical variability, such as the type of crosslinking agents employed to enrich for RNA-protein interactions (40). In addition, these discrepancies underscore the complexity of the host-viral interface: inside infected cells, host proteins may engage simultaneously with the viral genomic RNA, subgenomic viral RNAs, or both, adding layers of complexity to these dynamics. By using our predictive model, *cat*RAPID, we determined that the host proteins most frequently reported across different studies are, indeed, predicted to have higher interaction propensity towards SARS-CoV-2 RNA, compared to the non-interacting RBPs (44). This observation not only supports the strength of our predictions in identifying strong-affinity interactions but also forms the cornerstone of the strategy presented in this paper.

Our focus was primarily directed towards the most structured regions of SARS-CoV-2 RNAs, which are predicted to be highly contacted by human proteins (18).

We identified several proteins (e.g., FAM120A, HAT1, LSG1, RPL14, RPL18A, RPL24, RPL35, RPS6 SHMT1, and SYNE2) that were previously uncovered by studies conducted on SARS-CoV-2 infected cells (12–15). However, we noticed that some RBPs, reported to interact with SARS-CoV-2 RNA, were promiscuously interacting with our designed RNA fragments. Thus, these interactions likely contribute to response mechanisms triggered by the accumulation of viral RNA in the cell. The accumulation of viral RNA-host RBP interactions may necessitate the increase in local protein concentration offered by phase separation for the host proteins to sequester the viral RNA within liquid condensates (18, 44, 58). In fact, different RBPs we identified as promiscuous binders, such as G3BP1 and G3BP2 and CAPRIN1, PUM1, and PUM2, are implicated in the formation and composition of stress granules (58–60).

We also identified previously undisclosed SARS-CoV-2 RNA interactors, such as CEP350, GPKOW, and RGPD1, which, in our system, interact specifically with viral fragments containing the 5′ and 3′ UTR regions. These proteins appear to be key players in the genome cyclization process that potentially modulates SARS-CoV-2 RNA discontinuous transcription, as corroborated by recent reports (5, 47).

We were also able to link the interaction network of SARS-CoV-2 RNA with post-transcriptional modifications of the viral genomic and subgenomic RNAs. Indeed, SARS-CoV-2 RNA is heavily post-transcriptionally modified (61), although the role of each RNA modification has not been fully described. Modifications such as N6-methyladenosine (m6A), catalysed by METTL3, have been associated with the modulation of the viral RNA sensor RIG-I, which effectively circumvents the host innate immune response (10, 62). Similarly, pseudouridine, which is documented to be the most abundant modification identified in SARS-CoV-2 RNA (10), has been hypothesized to aid viruses to hijack the host immune sensors and evade the innate immune response (63). The writer enzyme responsible for this modification is still unknown, although PUS1, a member of the pseudouridine synthase family, was found to be associated with SARS-CoV-2 RNA (14).

Through our RaPID-MS analysis, we found PUS7, another member of the pseudouridine synthase family, binding to the RNA fragments present at both the 5′ and 3′ end regions of the viral genome. The retrieval of the PUS7 consensus sequence, UGUAR, within these fragments, paired with their differential electrical signal distribution compared to the unmodified *in vitro* transcribed RNA in nanopore DRS analysis, suggests PUS7 as the putative pseudouridylation agent for SARS-CoV-2 RNA.

Interestingly, we found three subgenomic RNAs with a modified UGUAR site, as identified by Nanocompore, within the stem-loop 2 (SL2) of the SARS-CoV-2 TRS-L. This phenomenon is likely to occur in all SARS-CoV-2 subgenomic RNAs, but it requires further improvements of the coverage at the 5′ end to be confidently scored by the nanopore DRS approach.

An important consequence of pseudouridylation is the potential effect on the RNA secondary structure and on its propensity to interact with both host and viral proteins. Specifically, we found evidence suggesting that PUS7-dependent pseudouridylation of the SARS-CoV-2 SL2 enhances the binding of the viral protein NSP1 to the viral 5′ UTR. An observation that fits the current model of NSP1-mediated regulation of the host translational machinery (8, 64, 65). Indeed, the non-structural protein NSP1 is a well-known factor in manipulating the host translational machinery (64, 66). By binding to the 40S ribosomal subunit, NSP1 obstructs the mRNA entry site, effectively hindering translation of host mRNAs (64, 66). However, SARS-CoV-2 subgenomic RNAs can elude NSP1-mediated translation inhibition thanks to the interaction of their 5′ UTRs with the N-terminal domain of NSP1, effectively dislodging the C-terminus from the ribosome entry channel (56). It is plausible that PUS7 acts as a pro-viral factor, aiding the pseudouridylation of the 5′ UTR of subgenomic RNAs to boost their translation rate. This hypothesis aligns with the recent proposal of NSP1 as a central element regulating RNA translation according to the affinity of each RNA towards NSP1 (65).

Both the sequence of SL2 and the NSP1-mediated regulation of translation are conserved in the Sarbecovirus subgenus of betacoronaviruses, including SARS-CoV-1 and SARS-CoV-2 (67, 68). This suggests that these viruses employ a shared strategy to prioritize the translation of their RNA transcripts over host mRNAs. Thus, a comprehensive understanding of the molecular mechanisms regulating the pseudouridylation of SARS-CoV-2 RNA could prove pivotal in devising innovative and targeted strategies to specifically impair the replication of sarbecoviruses in infected mammalian cells. In the future, the impairment of PUS7 activity could be exploited to interfere with SARS-CoV-2 replication. A PUS7 inhibitor, C17, has been recently described (69). Alternatively, RNA-molecules specifically targeting the SL2 sequence can be designed to eventually inhibit its pseudouridylation with a more specific and effective approach.

Overall, our study has unveiled that even a single occurrence of pseudouridylation can potentially alter the critical interplay between the virus and host machinery, instrumental for the biology of the virus. It is conceivable that these effects are exponentially amplified when considering the potential cumulative impact of additional pseudouridylation sites spread across the viral RNA. Moreover, other types of post-transcriptional modifications to the viral RNA, such as methylation or acetylation, could yield similar or even more drastic effects on the virus-host dynamics. As such, the interplay between the virus and its host can be viewed as a complex and refined landscape, with each post-transcriptional modification representing a potential point of intervention.

This captivating field of study has the potential for ground-breaking discoveries that could profoundly reshape our understanding of viral biology. The advent of innovative technologies, such as nanopore DRS and related methods, is opening new frontiers for exploring this terrain. These technologies offer the potential to sequence RNA molecules in real-time, providing unprecedented insights into the dynamics of RNA modifications. The capacity to probe the RNA molecules at the single-nucleotide level will enable us to map comprehensively these modifications across entire genomes. This could unlock new strategies to impede viral replication, either by disrupting the modification process itself or by altering its downstream effects. We foresee that this extensive understanding of RNA modification in SARS-CoV-2 and other viruses could modernise and improve our approach to antiviral therapies, transforming the ways to fight viral diseases.

## CODE AVAILABILITY

All the analyses are available at the Github directory: https://github.com/camillaugolini-iit/SARS-pseudoU.git.

## DATA AVAILABILITY

The mass spectrometry proteomics data have been deposited to the ProteomeXchange Consortium via the PRIDE (70) partner repository with the dataset identifier PXD034941’. Reviewer account details: **Username:**; **Password:** 1Me1DL7H.

The RNA sequencing datasets generated in this study have been deposited at the European Nucleotide Archive (ENA). Data is available under the ID: PRJEB53497. Reviewers can temporarily visualize the UCSC Genome Browser tracks at the following webpage: https://genome.ucsc.edu/s/camilla.ugolini/SARS%2DCoV%2D2%20putative%20pseudo.

## ACKNOWLEDGMENT

We thank Lorenzo di Tucci e Marco Rabozzi for developing an accelerated version of Nanopolish for preliminary analysis. The research leading to these results has been supported by the European Research Council (RIBOMYLOME_309545 and ASTRA_855923) and the H2020 projects (IASIS_727658 and INFORE_825080) to G.G.T; the Italian Association for Cancer Research (AIRC) - project IG 2019 (ID. 22851) to F.N. A grant from National Center for Gene Therapy and Drugs based on RNA Technology (CN00000041, EPNRRCN3) supported by European Union – NextGenerationEU PNRR MUR - M4C2 to F.N. E.Z. received funding from MINDED fellowship of the European Union’s Horizon 2020 research and innovation program under the Marie Skłodowska-Curie grant agreement No. 754490.

