## Supplementary figures and images for "Discovering host protein interactions specific for SARS-CoV-2 RNA genome"

### Supplementary figure 1

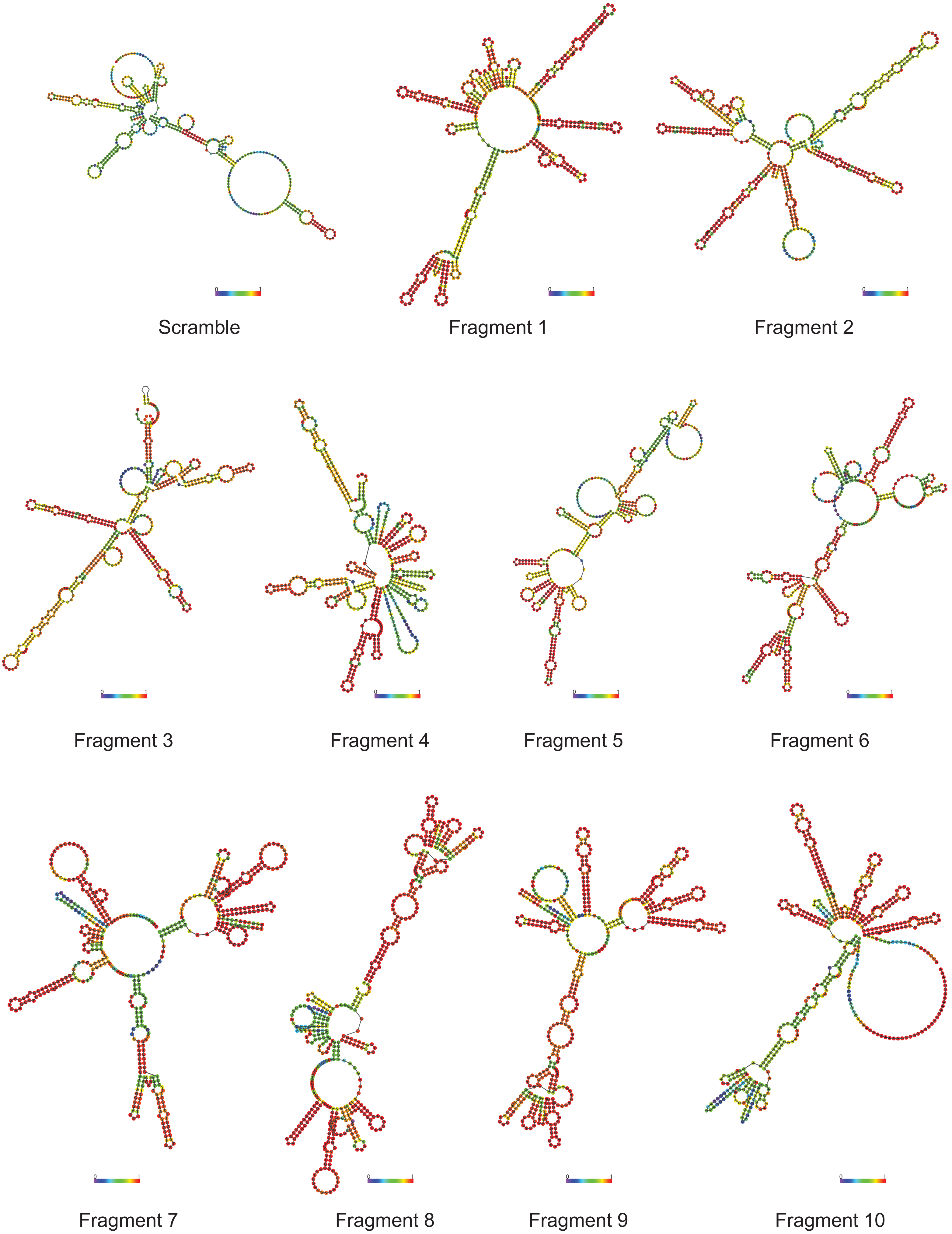


**Supplementary Figure 1.**

### Supplementary figure 2

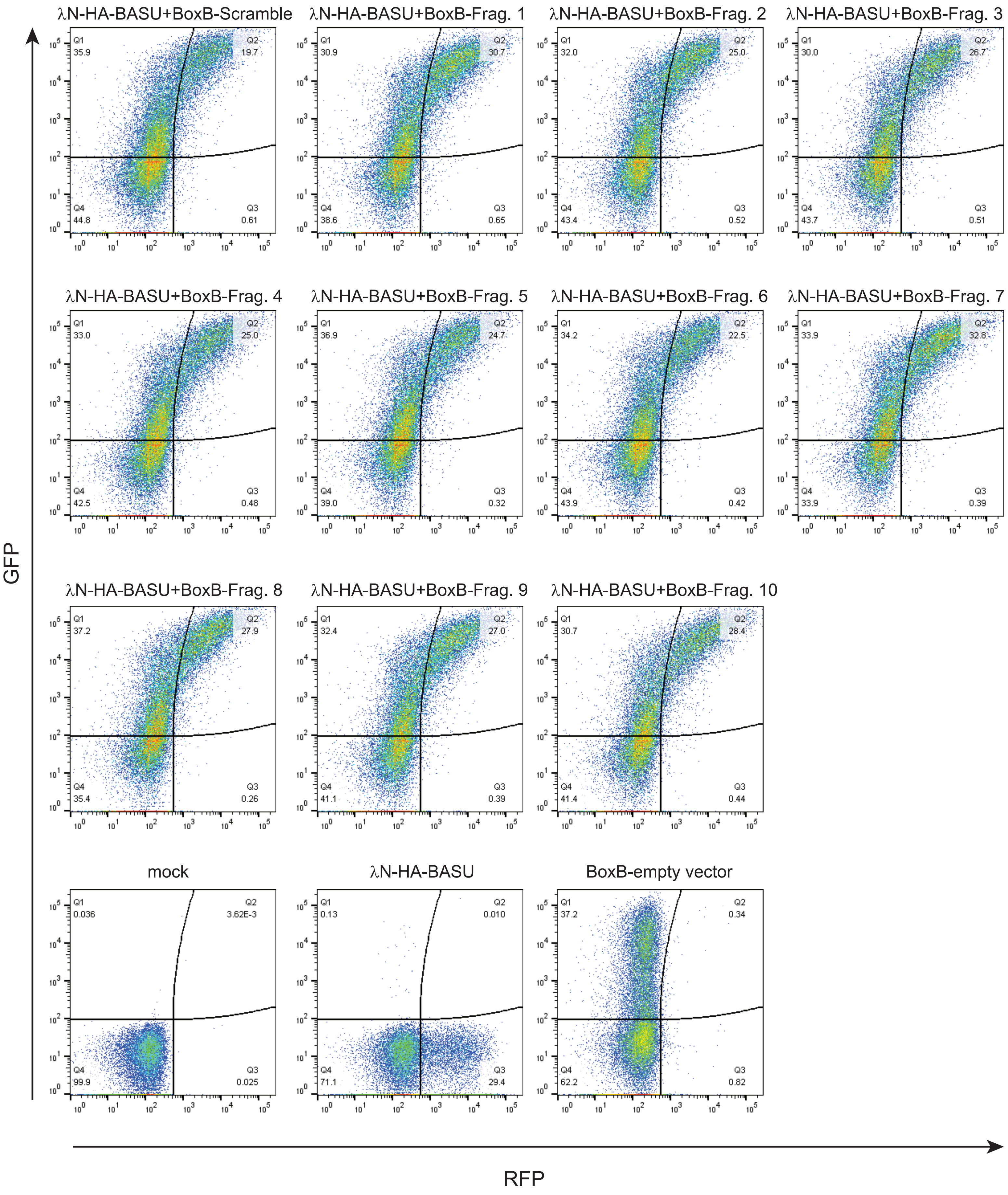


**Supplementary Figure 2.**

### Supplementary figure 3

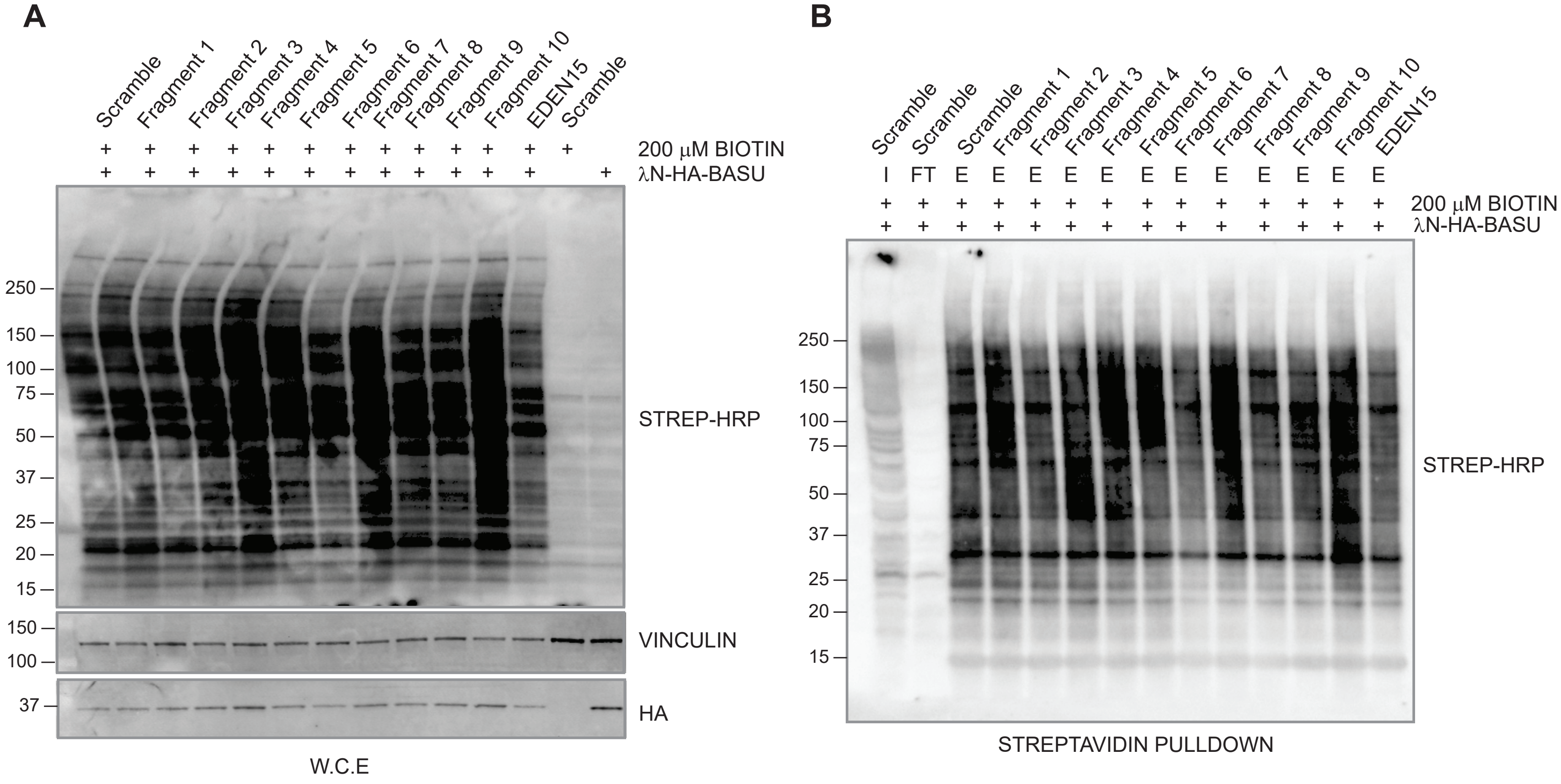


**Supplementary Figure 3.**

### Supplementary figure 4

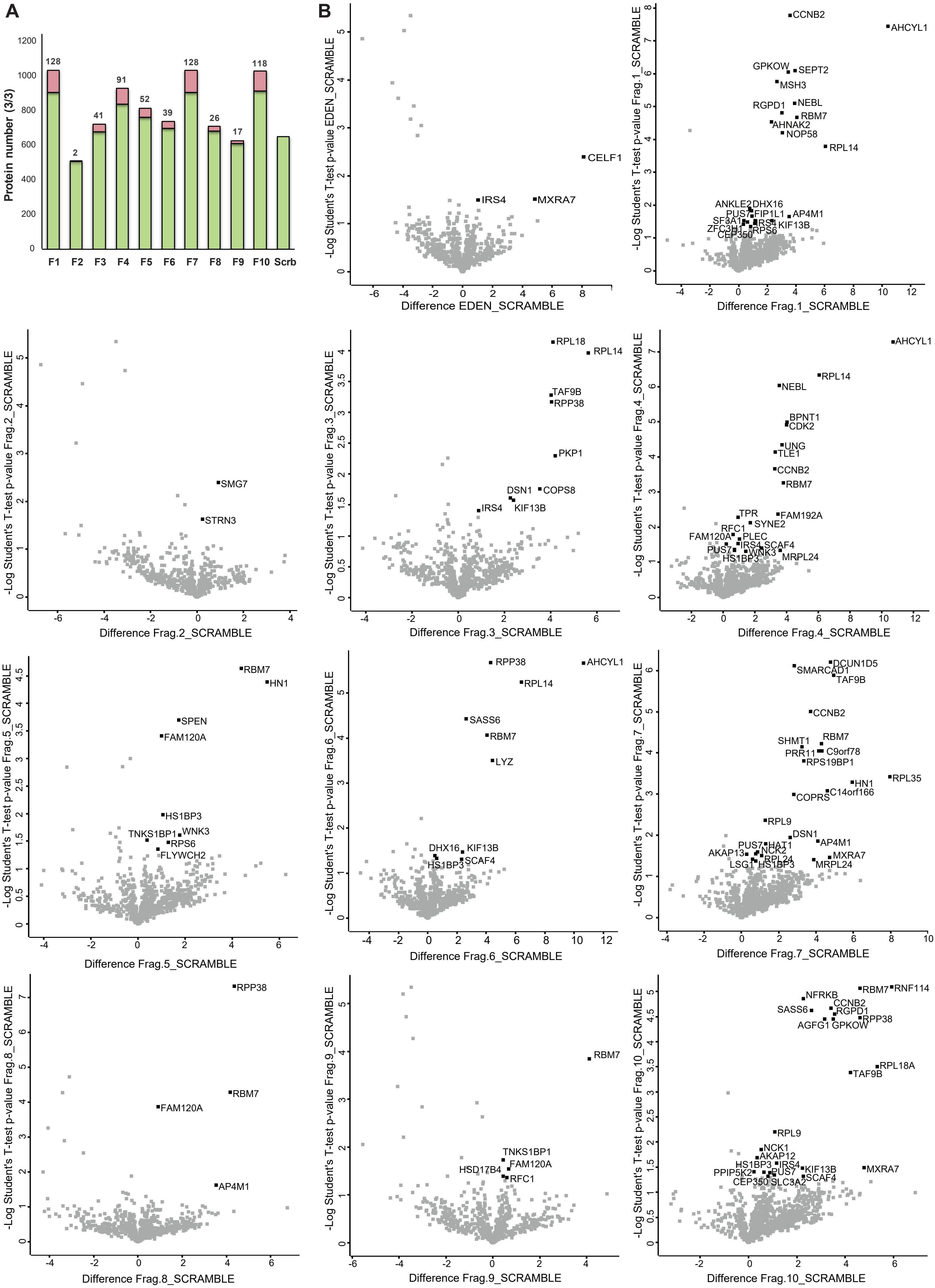


**Supplementary Figure 4.**

### Supplementary figure 6

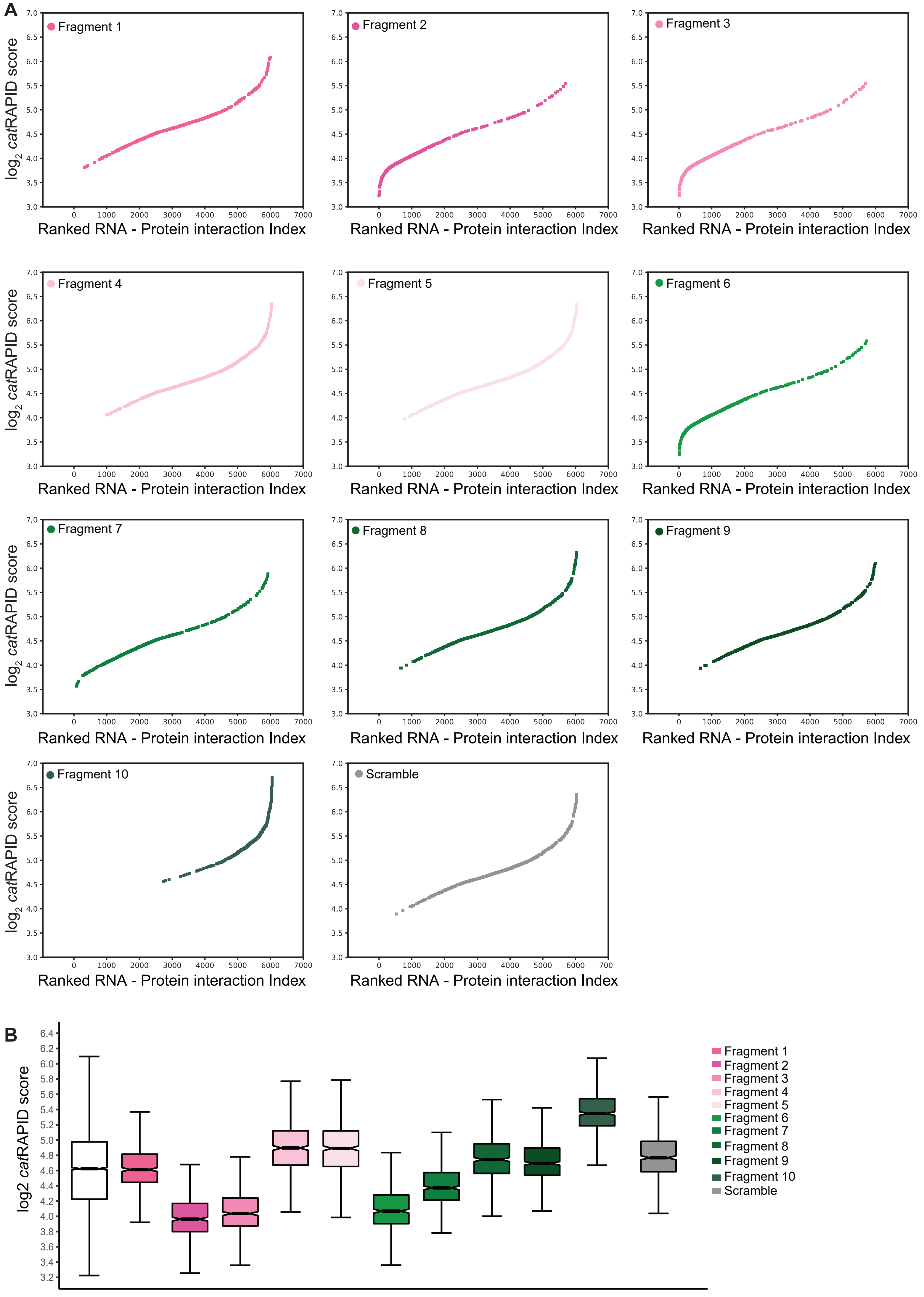


**Supplementary Figure 6.**

### Supplementary figure 7

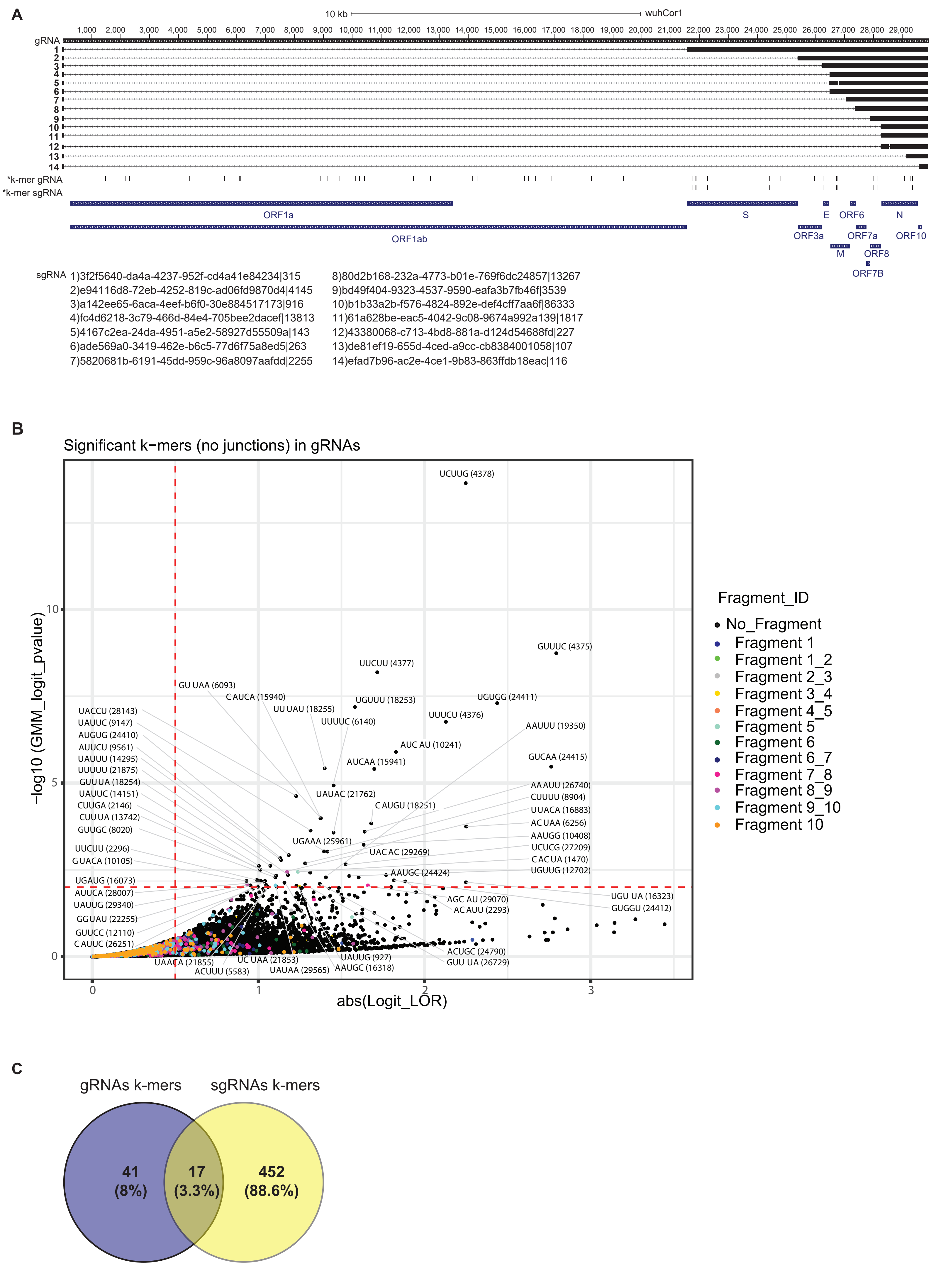


**Supplementary Figure 7**

### Supplementary figure 9

Site 23622-23638 , SPIKE, id: 3f2f5640-da4a-4237-952f-cd4a41e84234|315

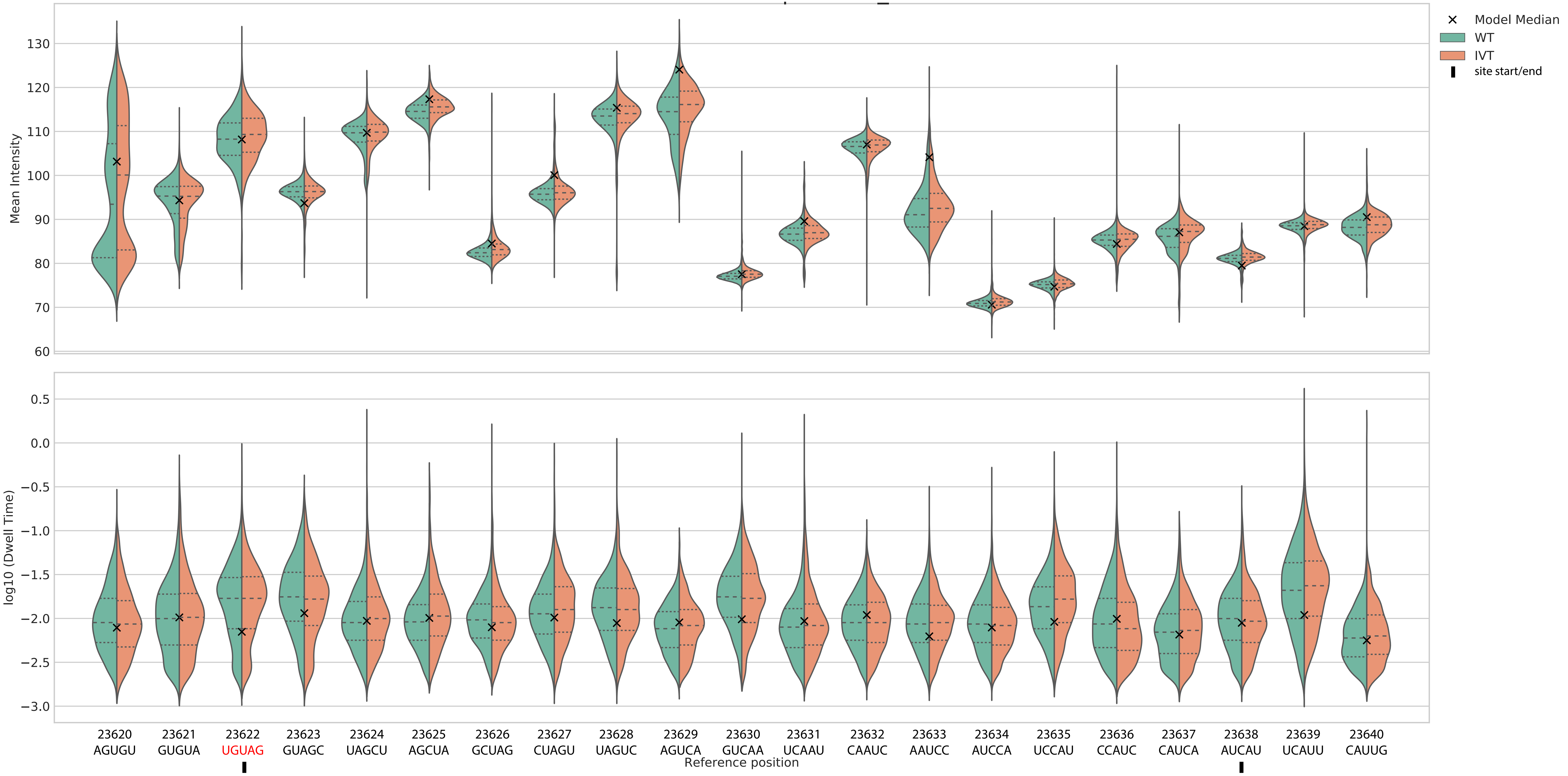

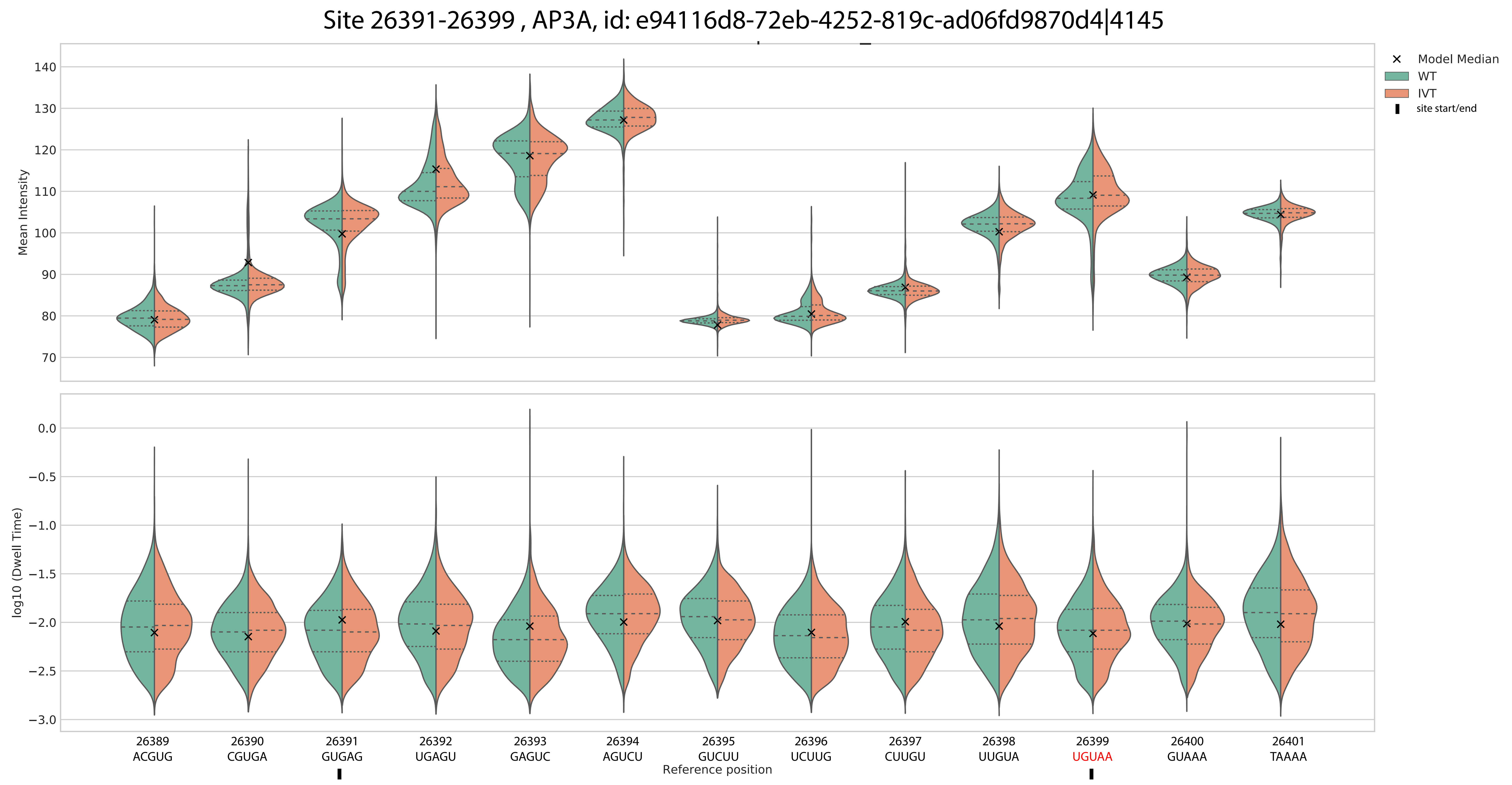

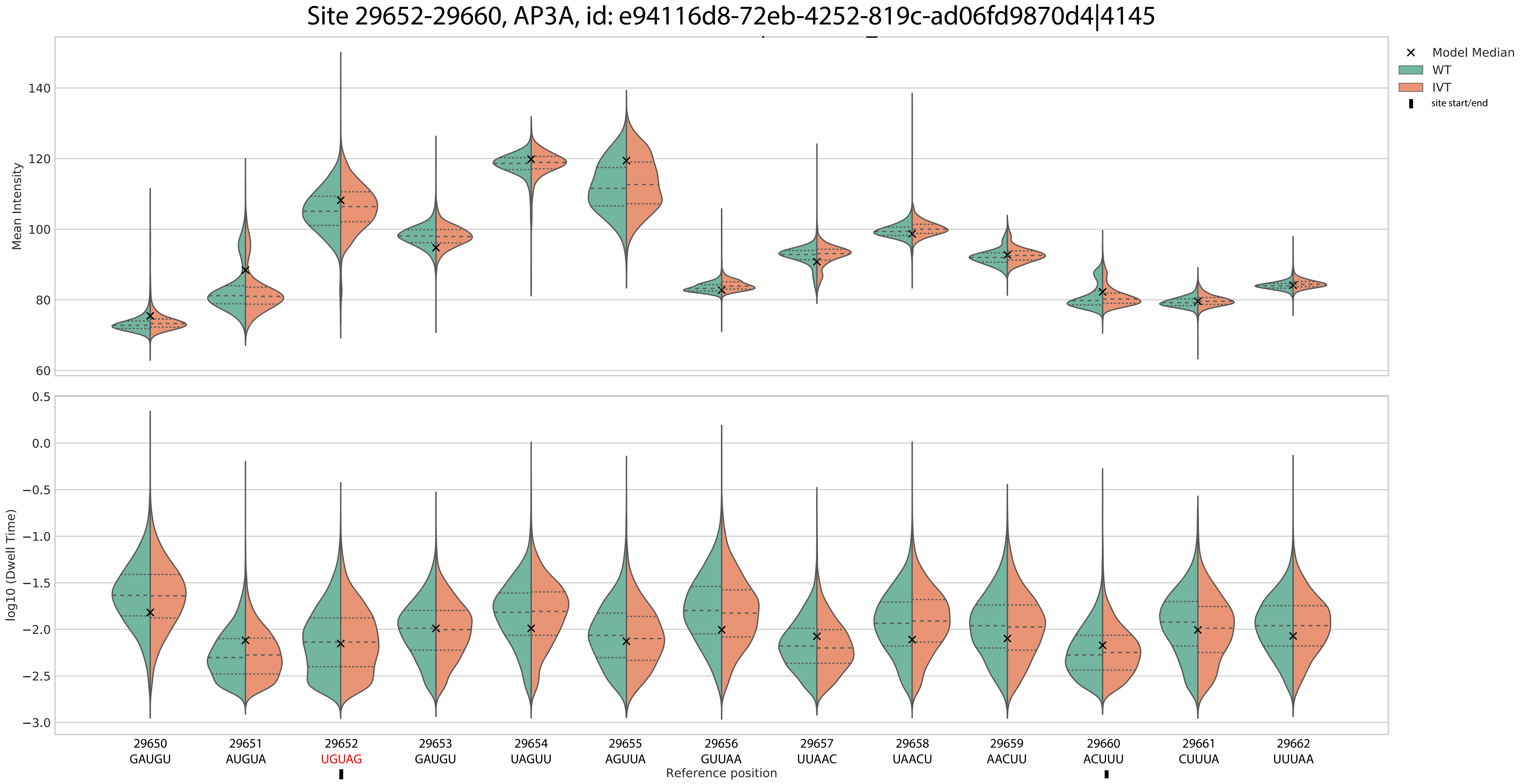

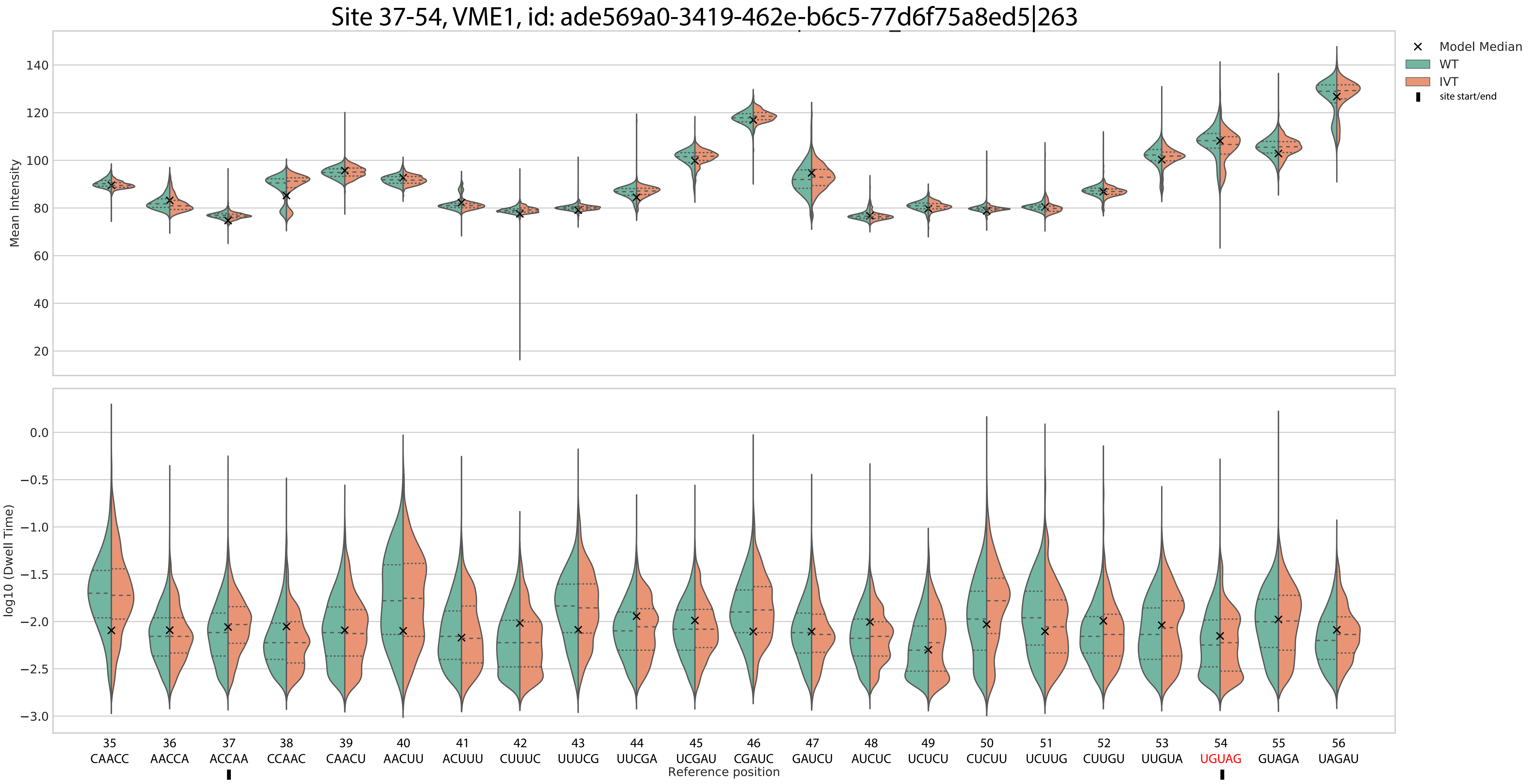

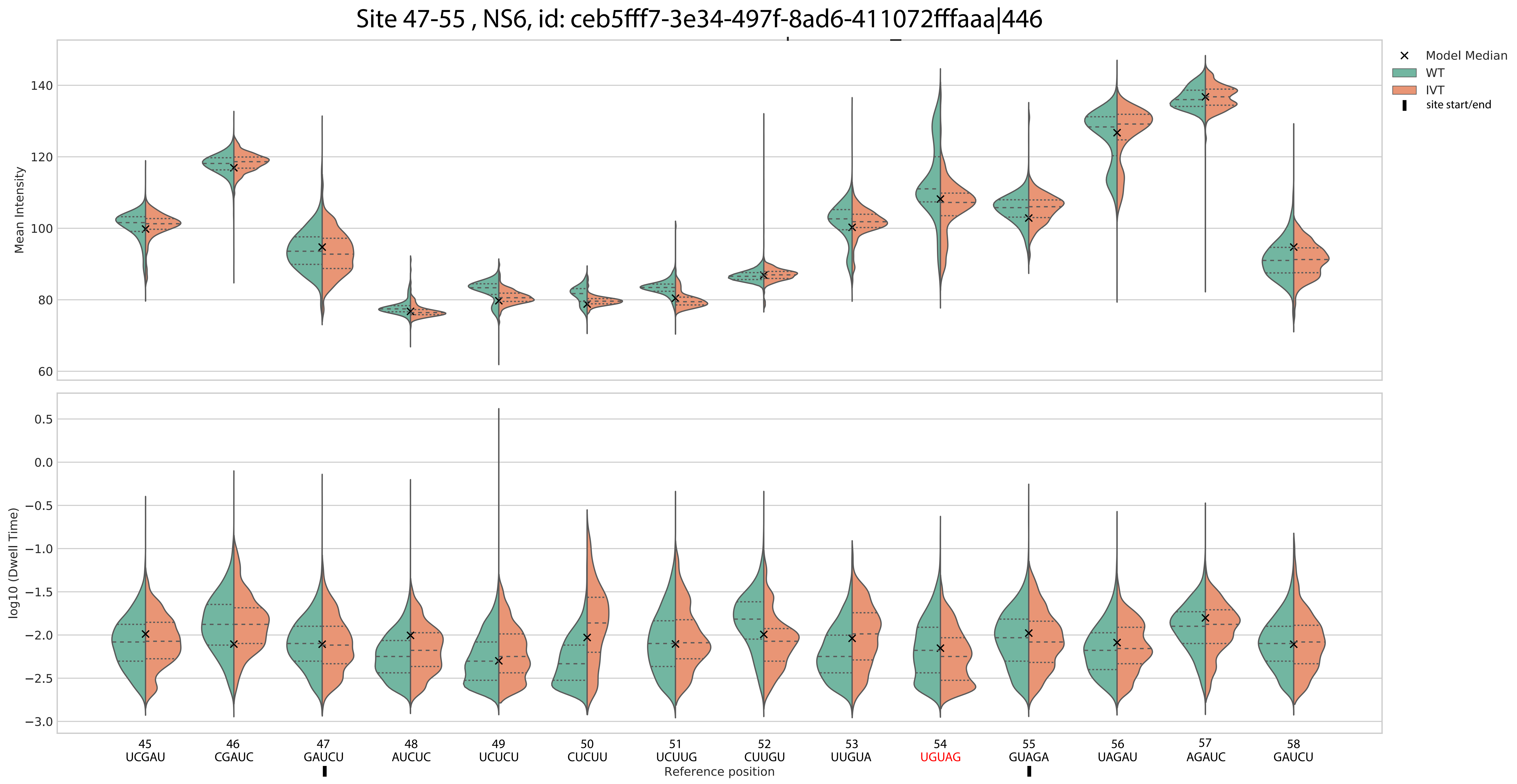

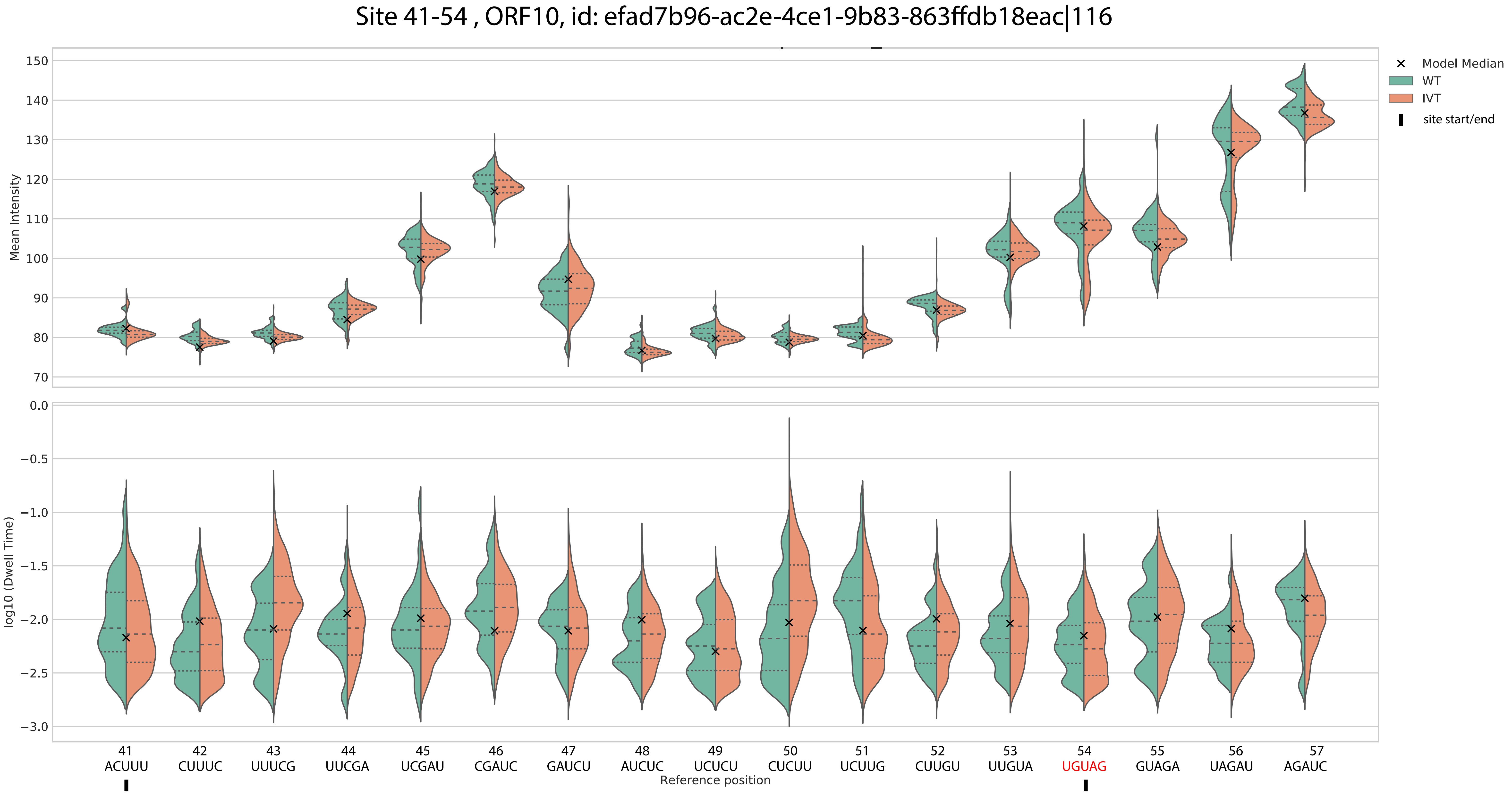

Site 29640-29678, ORF10, id: efad7b96-ac2e-4ce1-9b83-863ffdb18eac|116

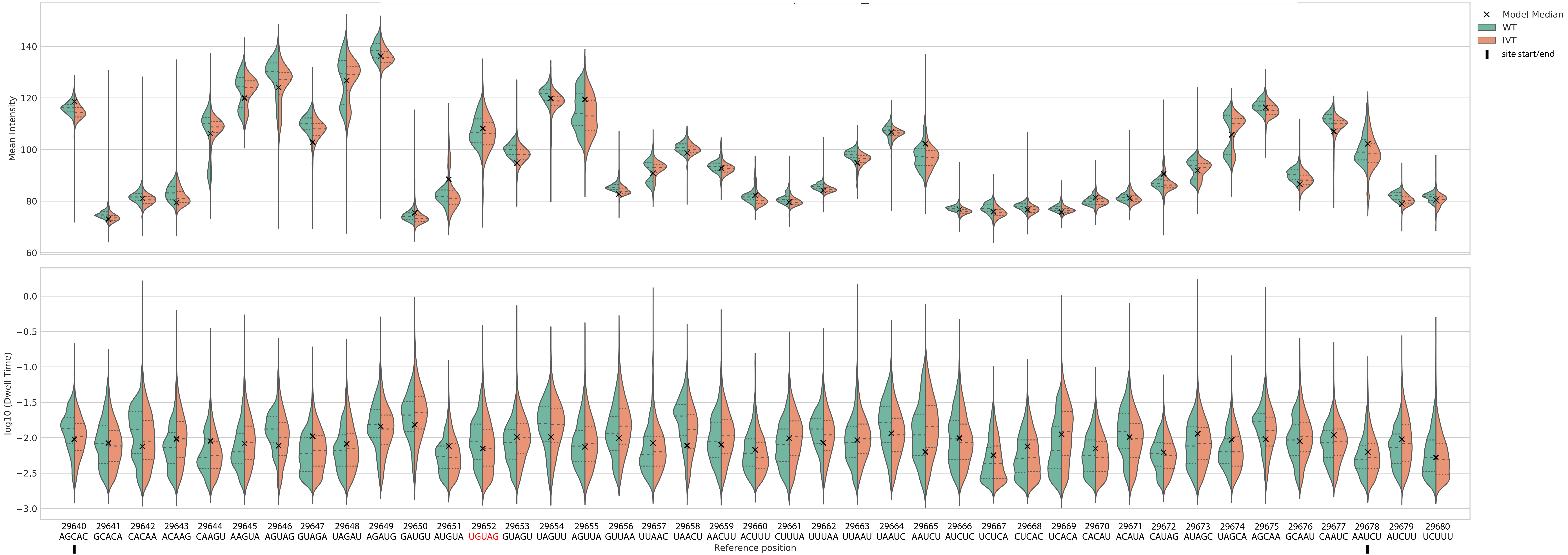

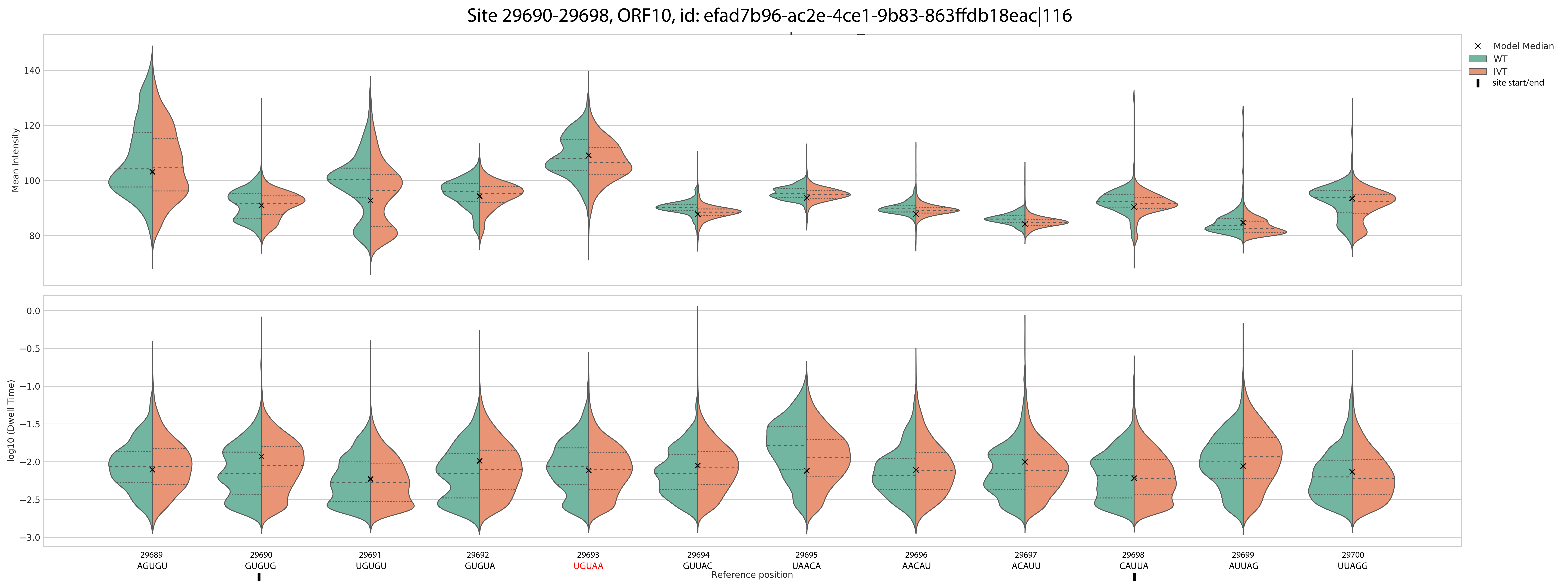

### Supplementary figure 10

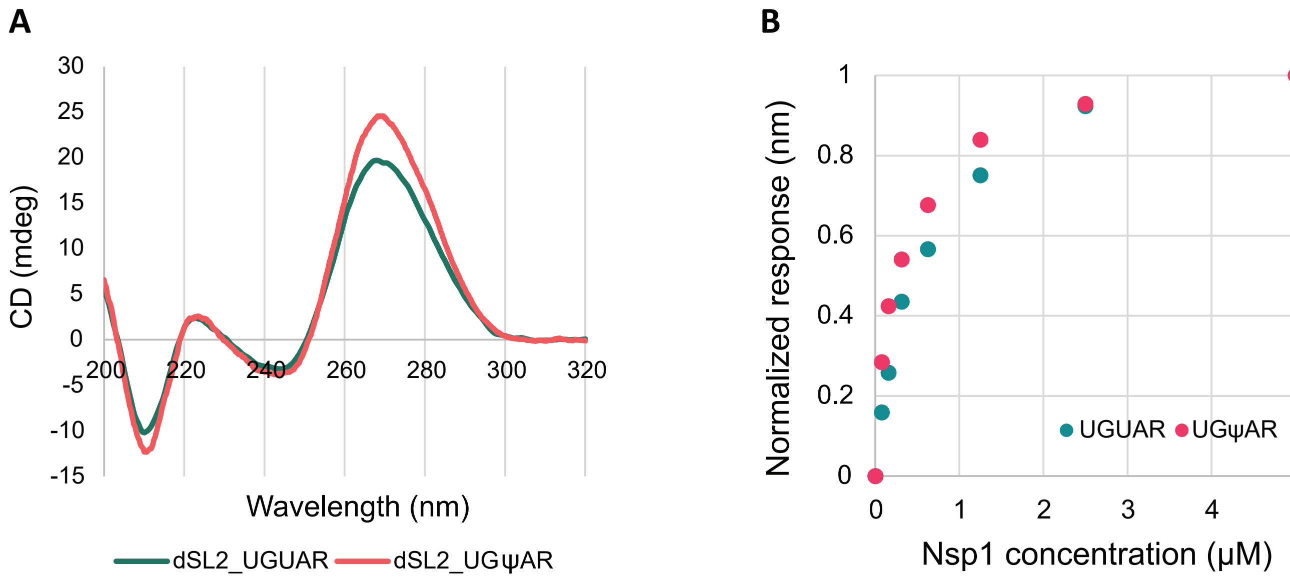


**Supplementary Figure 10**
