## Supplementary figure 5 for "Discovering host protein interactions specific for SARS-CoV-2 RNA genome"

A

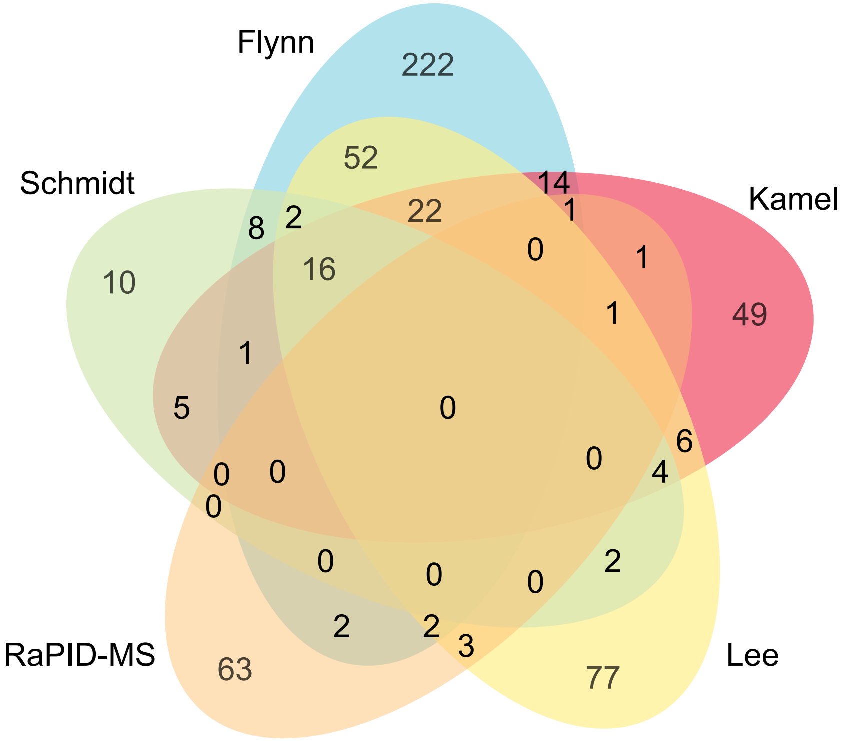


B

Common elements found in RaPID-MS and published datasets

**FAM120A**
**HAT1**
**LSG1**
**RPL14**
**RPL18A**
**RPL24**
**RPL35**
**RPS6**
**SHMT1**
**SYNE2**

C

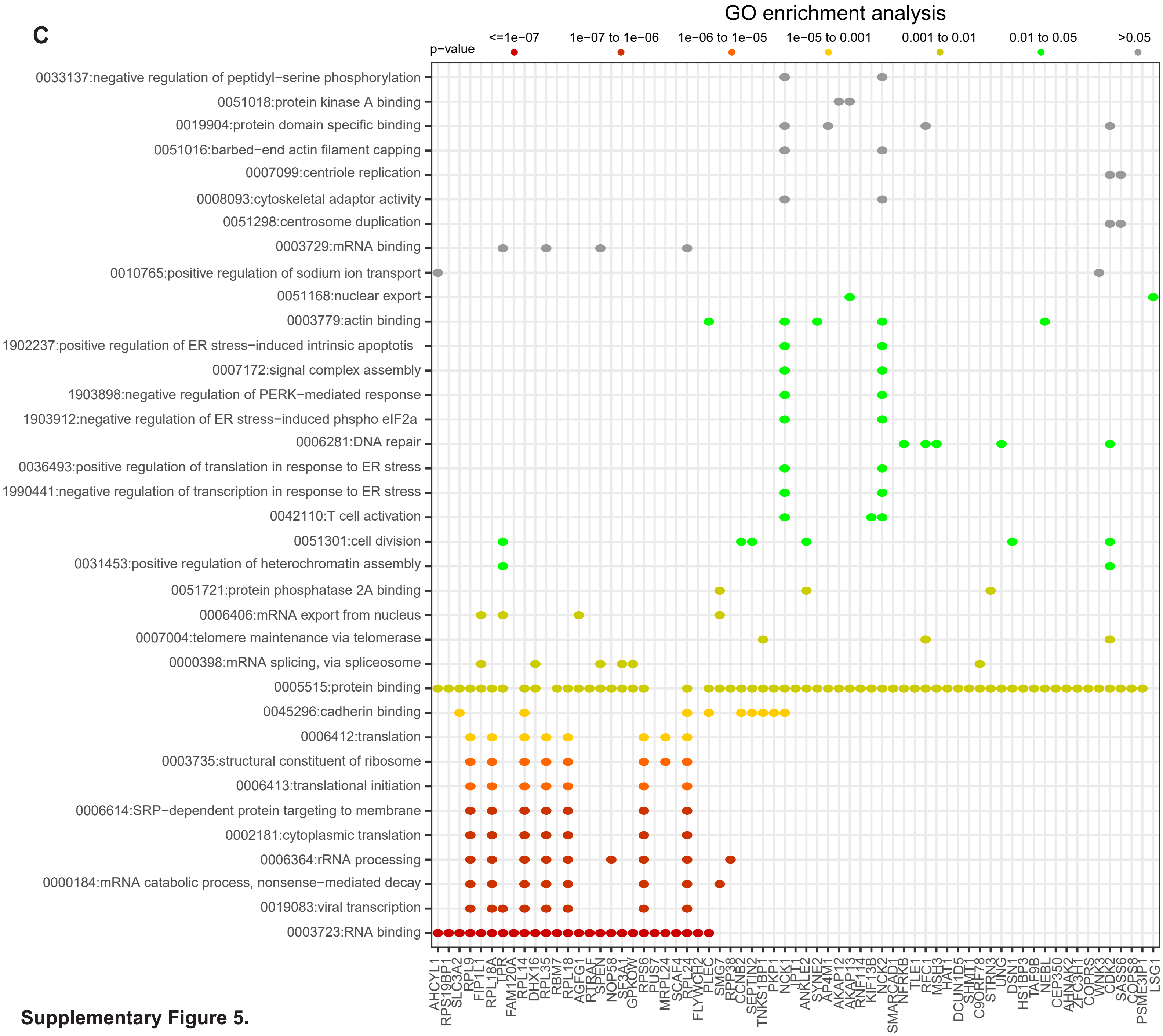


**Supplementary Figure 5.**
