## Supplementary figure 8 for "Discovering host protein interactions specific for SARS-CoV-2 RNA genome"

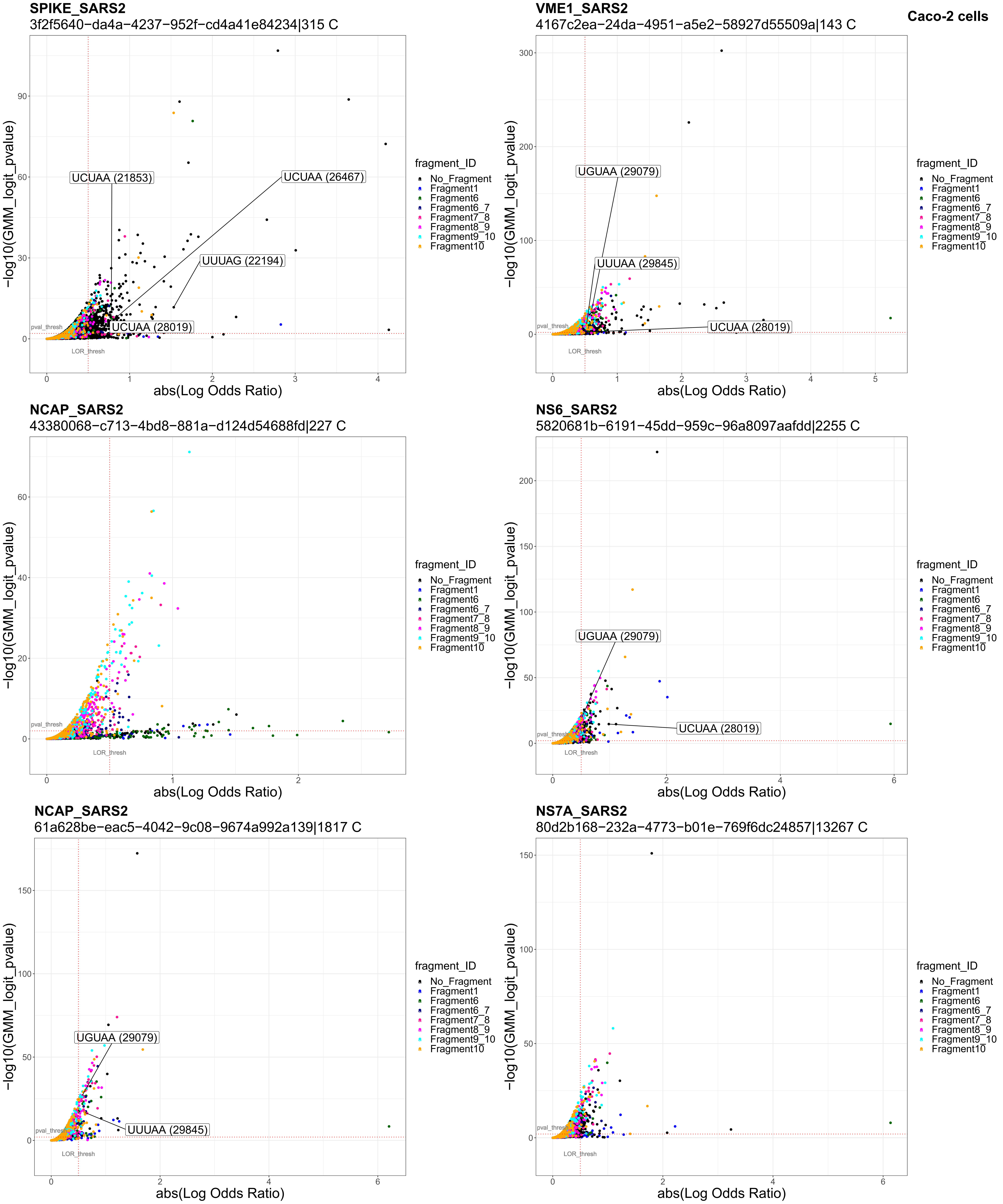
Caco-2 cells
SPIKE_SARS2
3f2f5640−da4a−4237−952f−cd4a41e84234|315 C
UCUAA (21853)
UCUAA (26467)
UUUAG (22194)
UCUAA (28019)
pval_thresh
LOR_thresh
−log10(GMM_logit_pvalue)
abs(Log Odds Ratio)
VME1_SARS2
4167c2ea−24da−4951−a5e2−58927d55509a|143 C
UGUAA (29079)
UUUAA (29845)
UCUAA (28019)
NCAP_SARS2
43380068−c713−4bd8−881a−d124d54688fd|227 C
NS6_SARS2
5820681b−6191−45dd−959c−96a8097aafdd|2255 C
UGUAA (29079)
UCUAA (28019)
NCAP_SARS2
61a628be−eac5−4042−9c08−9674a992a139|1817 C
UGUAA (29079)
UUUAA (29845)
NS7A_SARS2
80d2b168−232a−4773−b01e−769f6dc24857|13267 C
fragment_ID
No_Fragment
Fragment1
Fragment6
Fragment6_7
Fragment7_8
Fragment8_9
Fragment9_10
Fragment10

**VEMP_SARS2**
a142ee65−6aca−4eef−b6f0−30e884517173|916 C

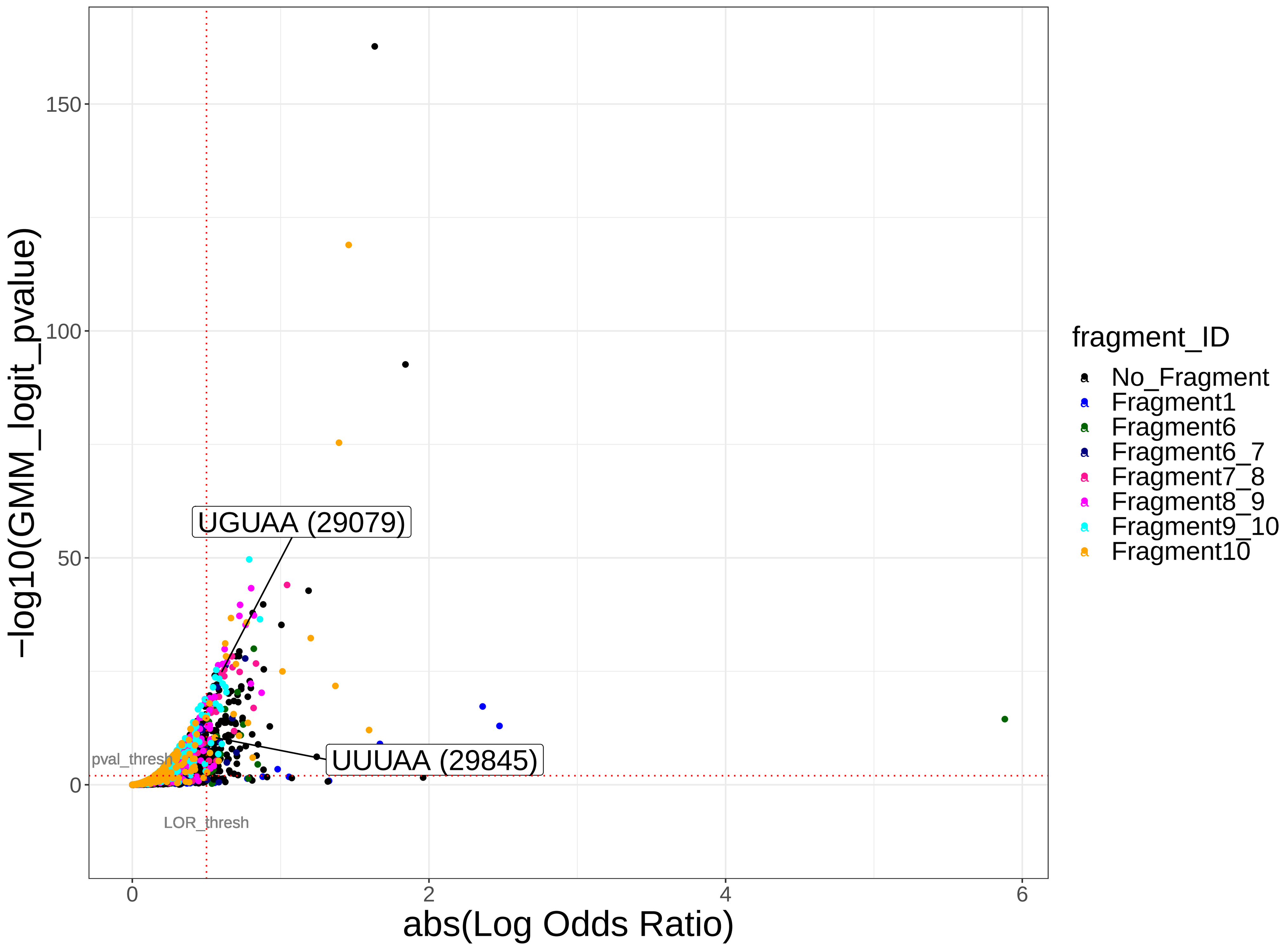


**VME1_SARS2**
ade569a0−3419−462e−b6c5−77d6f75a8ed5|263 C

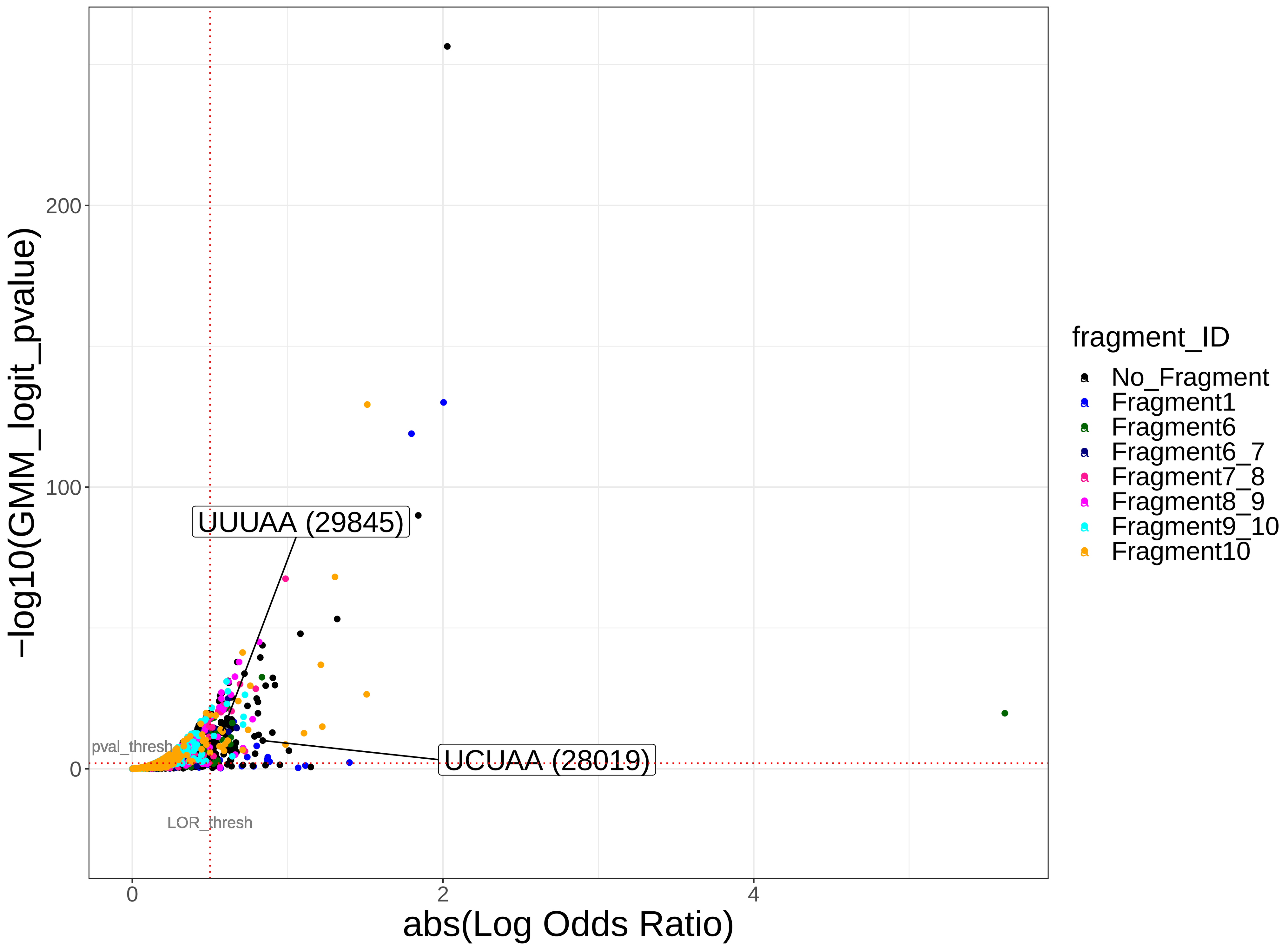


**NCAP_SARS2**
b1b33a2b−f576−4824−892e−def4cff7aa6f|86333 C

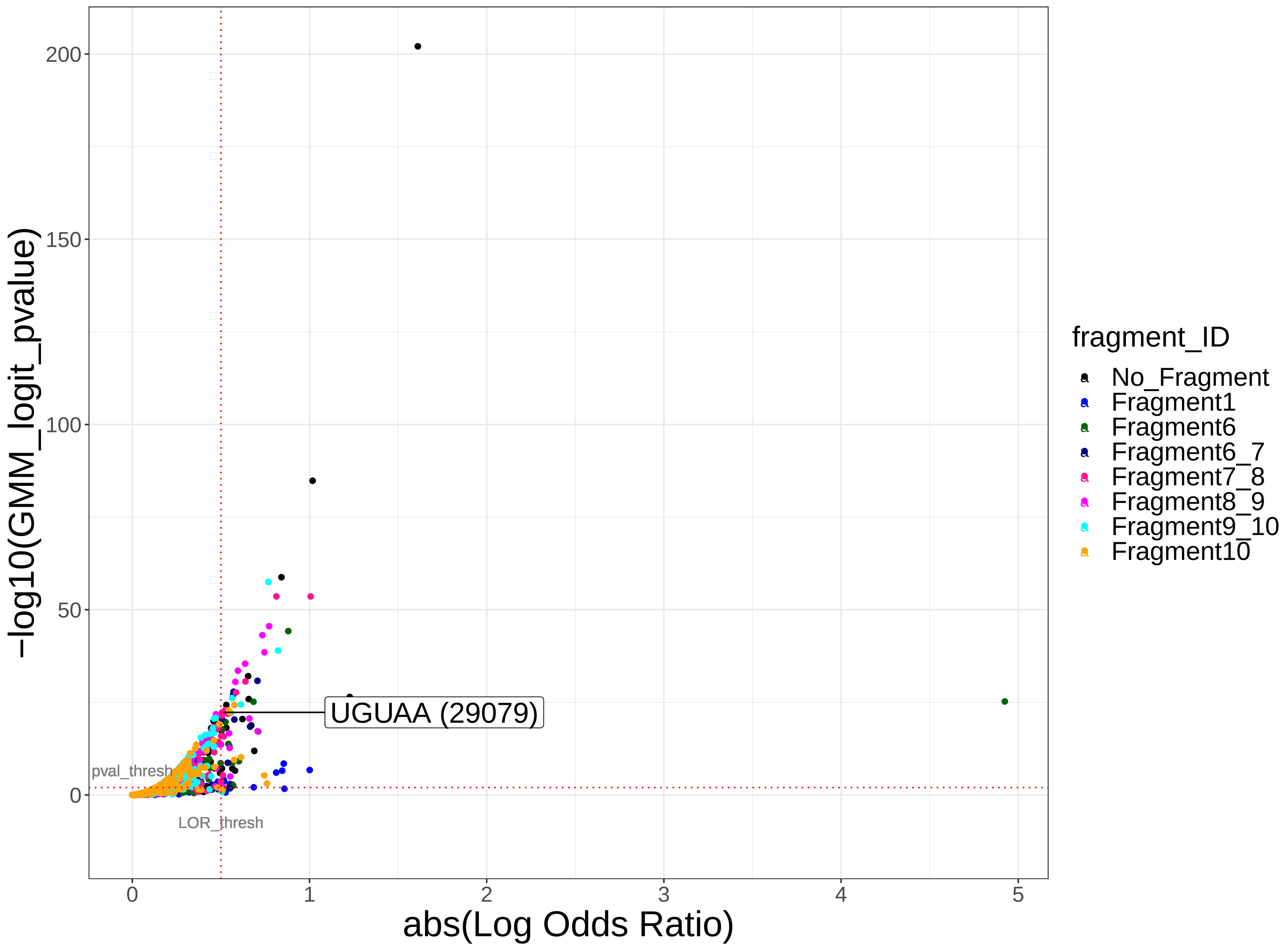


**NS8_SARS2**
bd49f404−9323−4537−9590−eafa3b7fb46f|3539 C

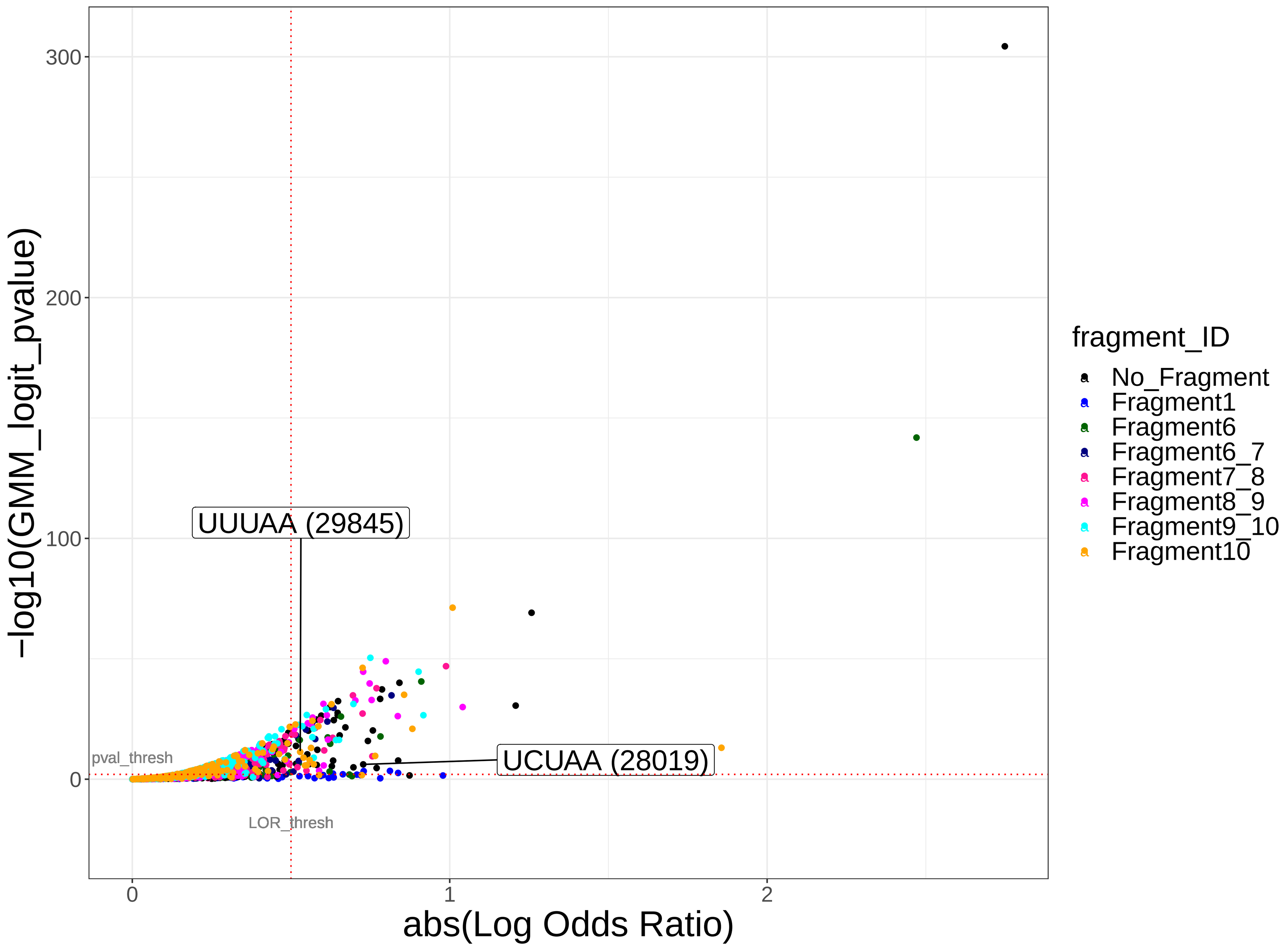


**ORF9D_SARS2**
de81ef19−655d−4ced−a9cc−cb8384001058|107 C

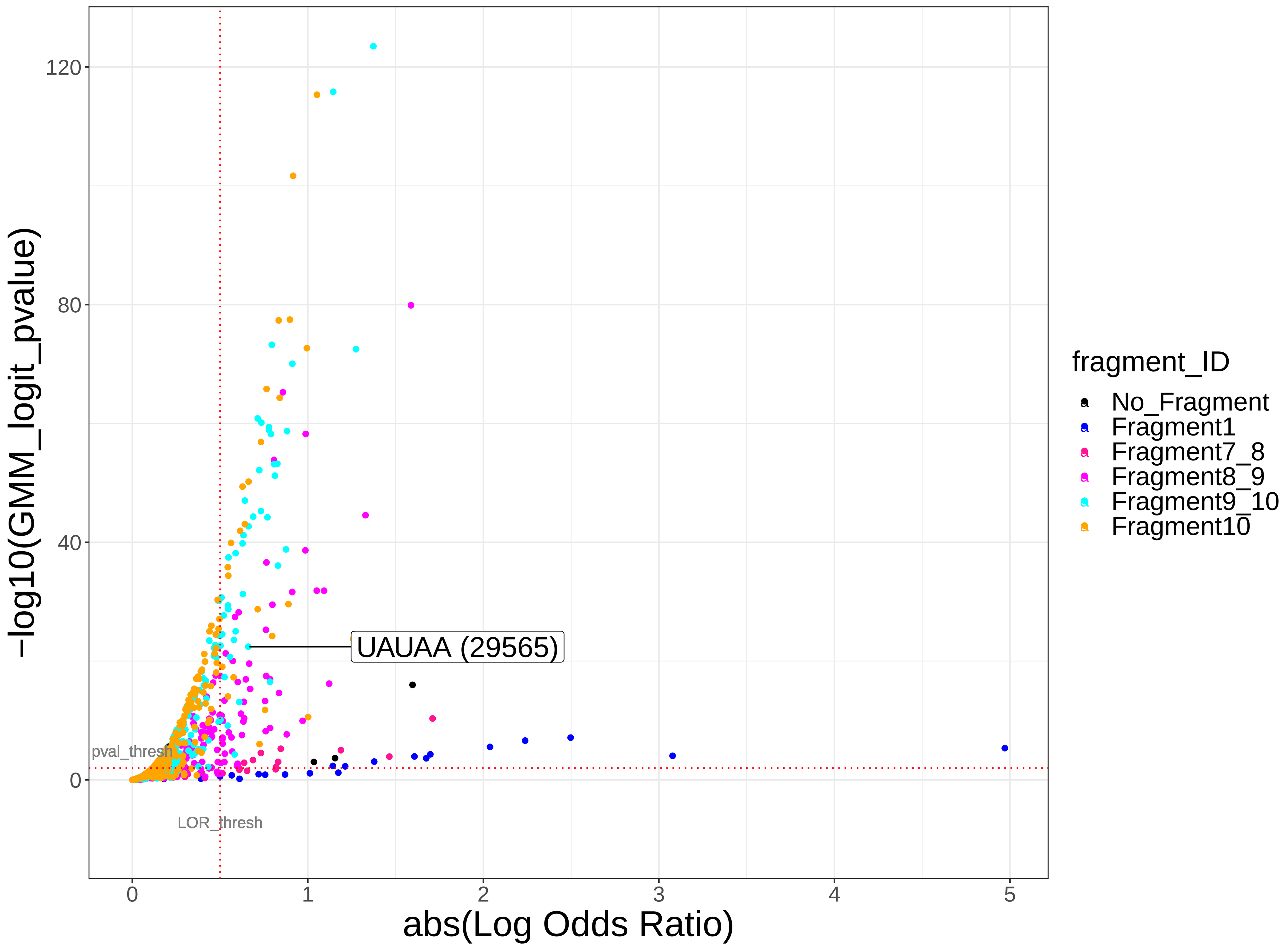


**AP3A_SARS2**
e94116d8−72eb−4252−819c−ad06fd9870d4|4145 C

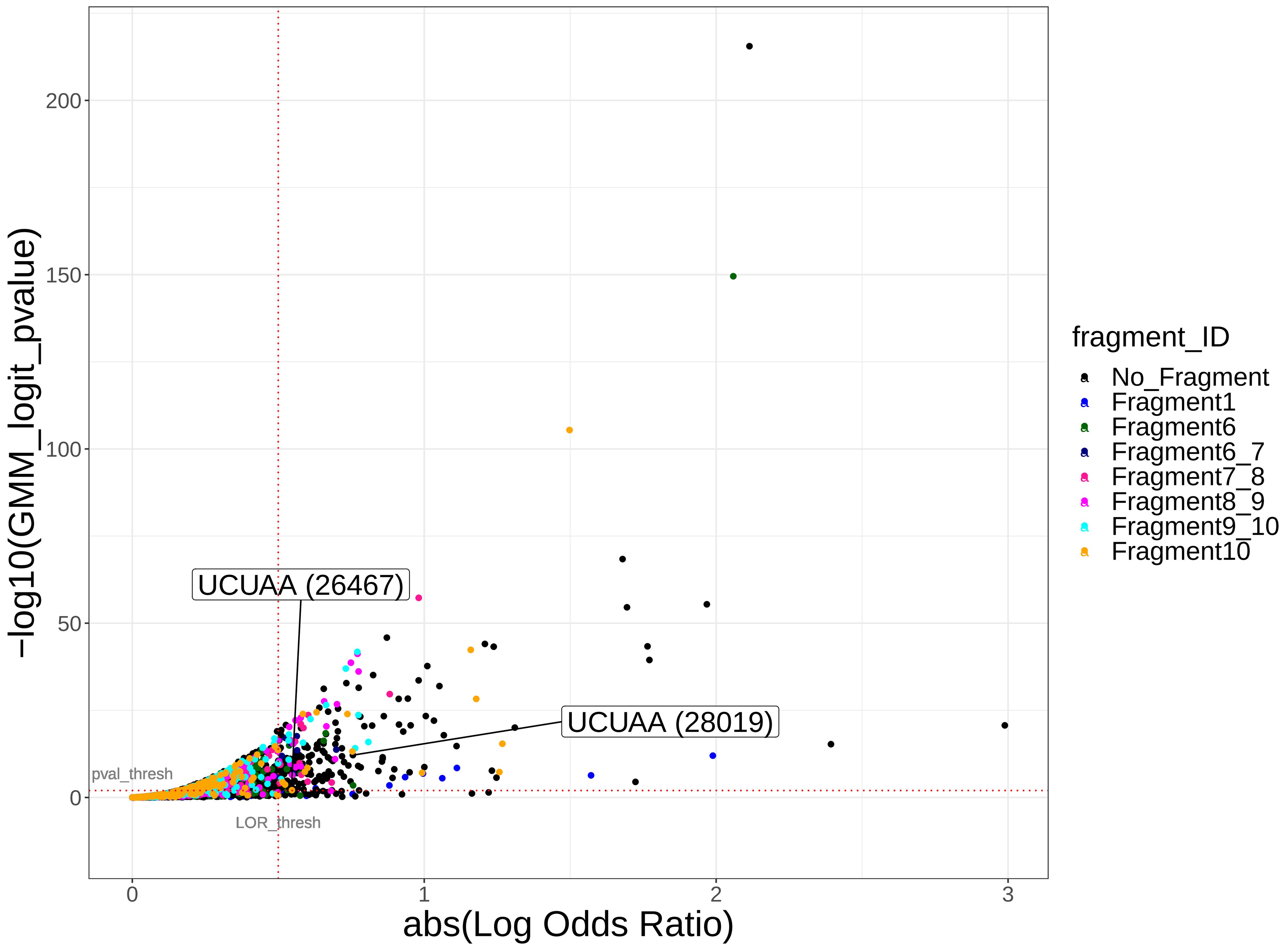

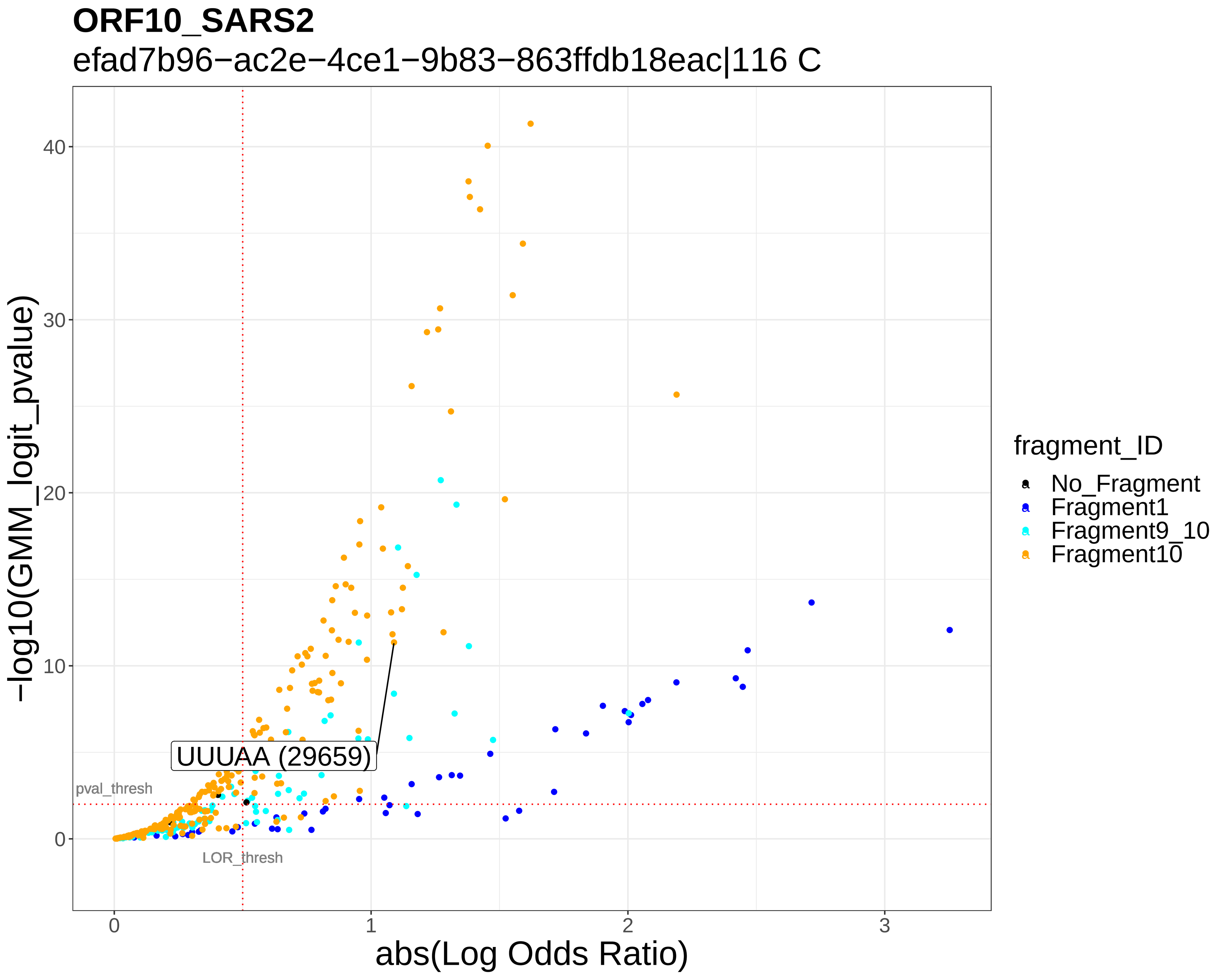
ORF10_SARS2
efad7b96−ac2e−4ce1−9b83−863ffdb18eac|116 C
−log10(GMM_logit_pvalue)
abs(Log Odds Ratio)
UUUAA (29659)
pval_thresh
LOR_thresh
fragment_ID
No_Fragment
Fragment1
Fragment9_10
Fragment10

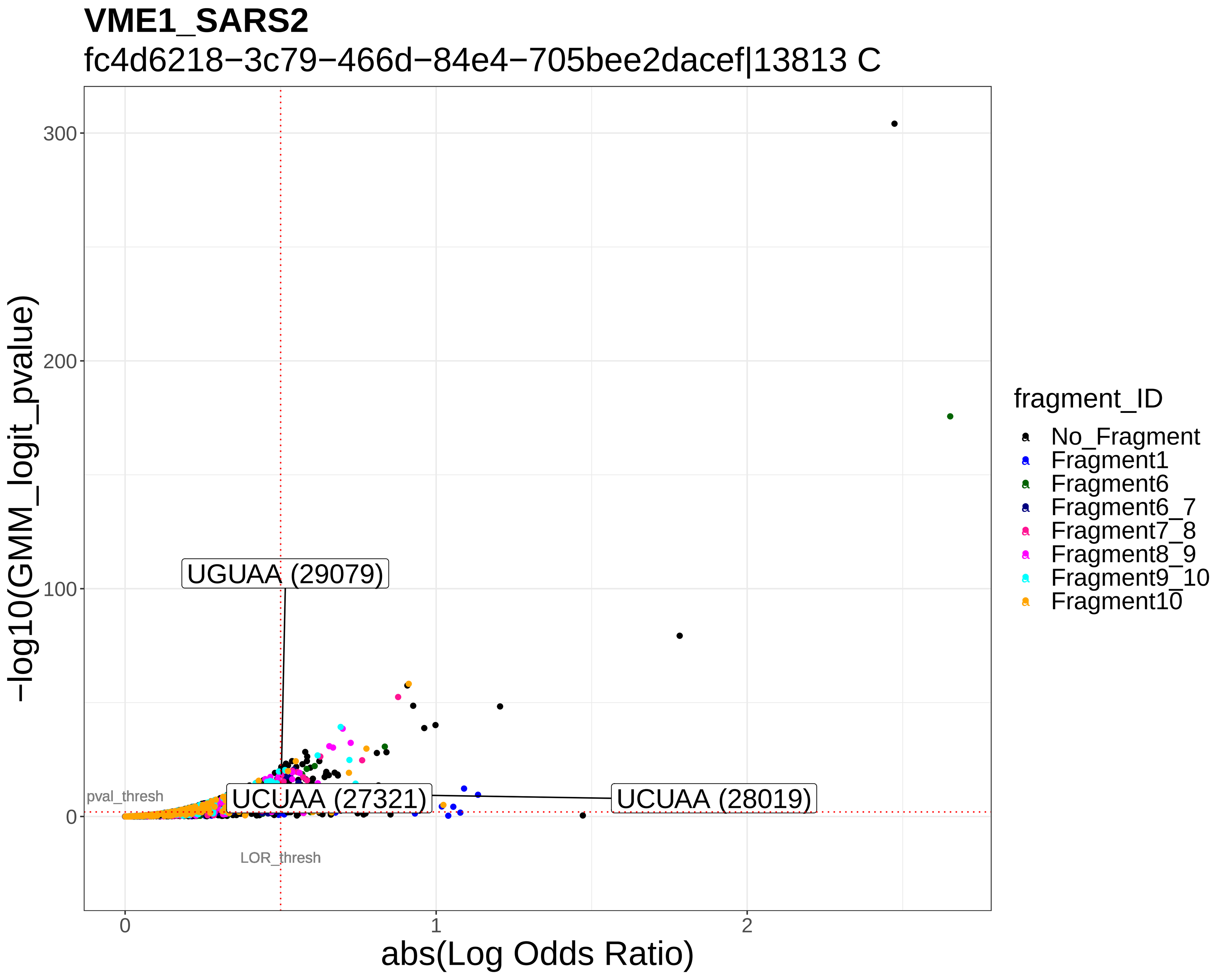
VME1_SARS2
fc4d6218−3c79−466d−84e4−705bee2dacef|13813 C
−log10(GMM_logit_pvalue)
abs(Log Odds Ratio)
UGUAA (29079)
UCUAA (27321)
UCUAA (28019)
pval_thresh
LOR_thresh
fragment_ID
No_Fragment
Fragment1
Fragment6
Fragment6_7
Fragment7_8
Fragment8_9
Fragment9_10
Fragment10

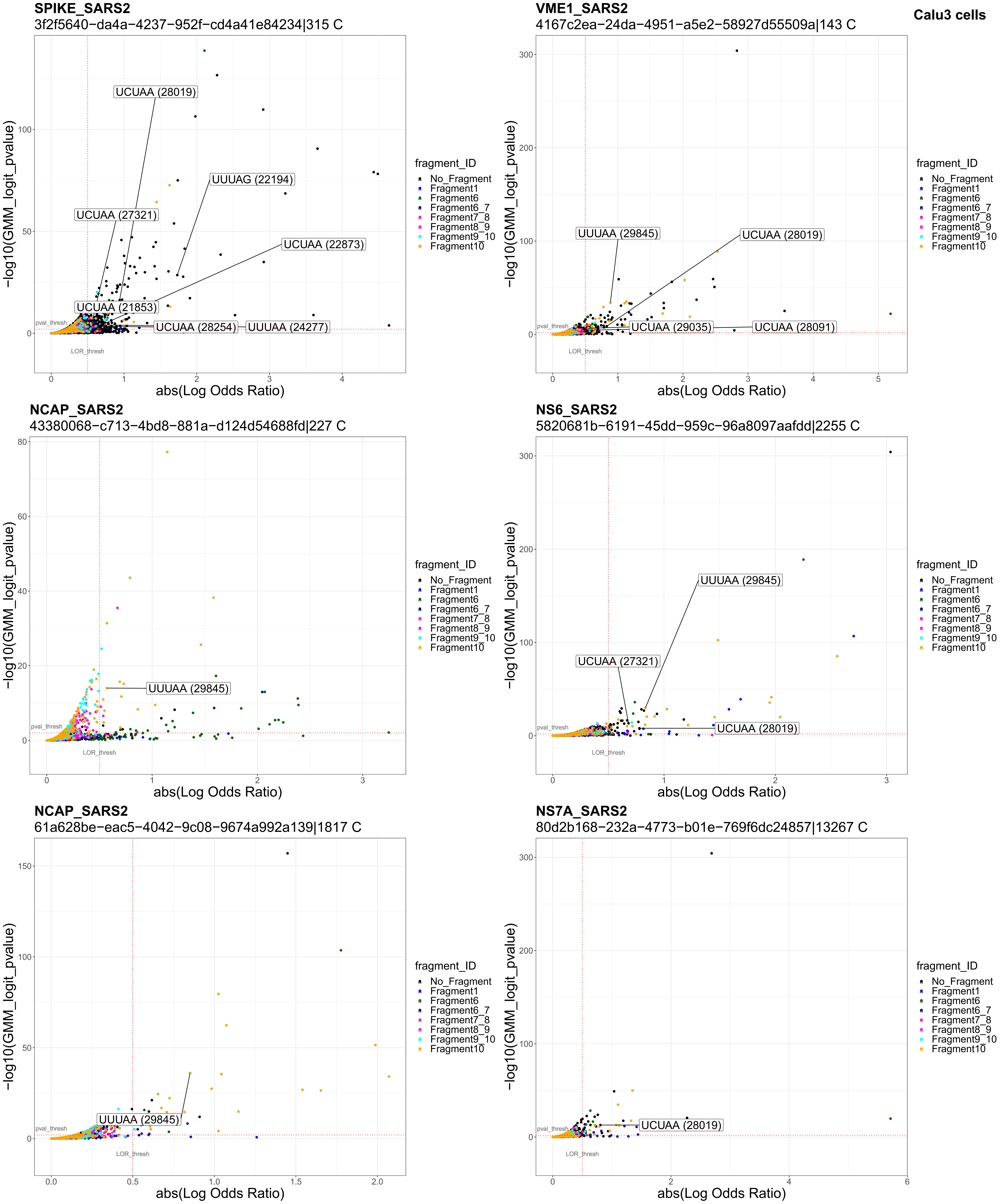
Calu3 cells
SPIKE_SARS2
3f2f5640−da4a−4237−952f−cd4a41e84234|315 C
UCUAA (28019)
UUUAG (22194)
UCUAA (27321)
UCUAA (22873)
UCUAA (21853)
UCUAA (28254)
UUUAA (24277)
VME1_SARS2
4167c2ea−24da−4951−a5e2−58927d55509a|143 C
UUUAA (29845)
UCUAA (28019)
UCUAA (29035)
UCUAA (28091)
NCAP_SARS2
43380068−c713−4bd8−881a−d124d54688fd|227 C
UUUAA (29845)
NS6_SARS2
5820681b−6191−45dd−959c−96a8097aafdd|2255 C
UUUAA (29845)
UCUAA (27321)
UCUAA (28019)
NCAP_SARS2
61a628be−eac5−4042−9c08−9674a992a139|1817 C
UUUAA (29845)
NS7A_SARS2
80d2b168−232a−4773−b01e−769f6dc24857|13267 C
UCUAA (28019)
−log10(GMM_logit_pvalue)
abs(Log Odds Ratio)
pval_thresh
LOR_thresh
fragment_ID
No_Fragment
Fragment1
Fragment6
Fragment6_7
Fragment7_8
Fragment8_9
Fragment9_10
Fragment10

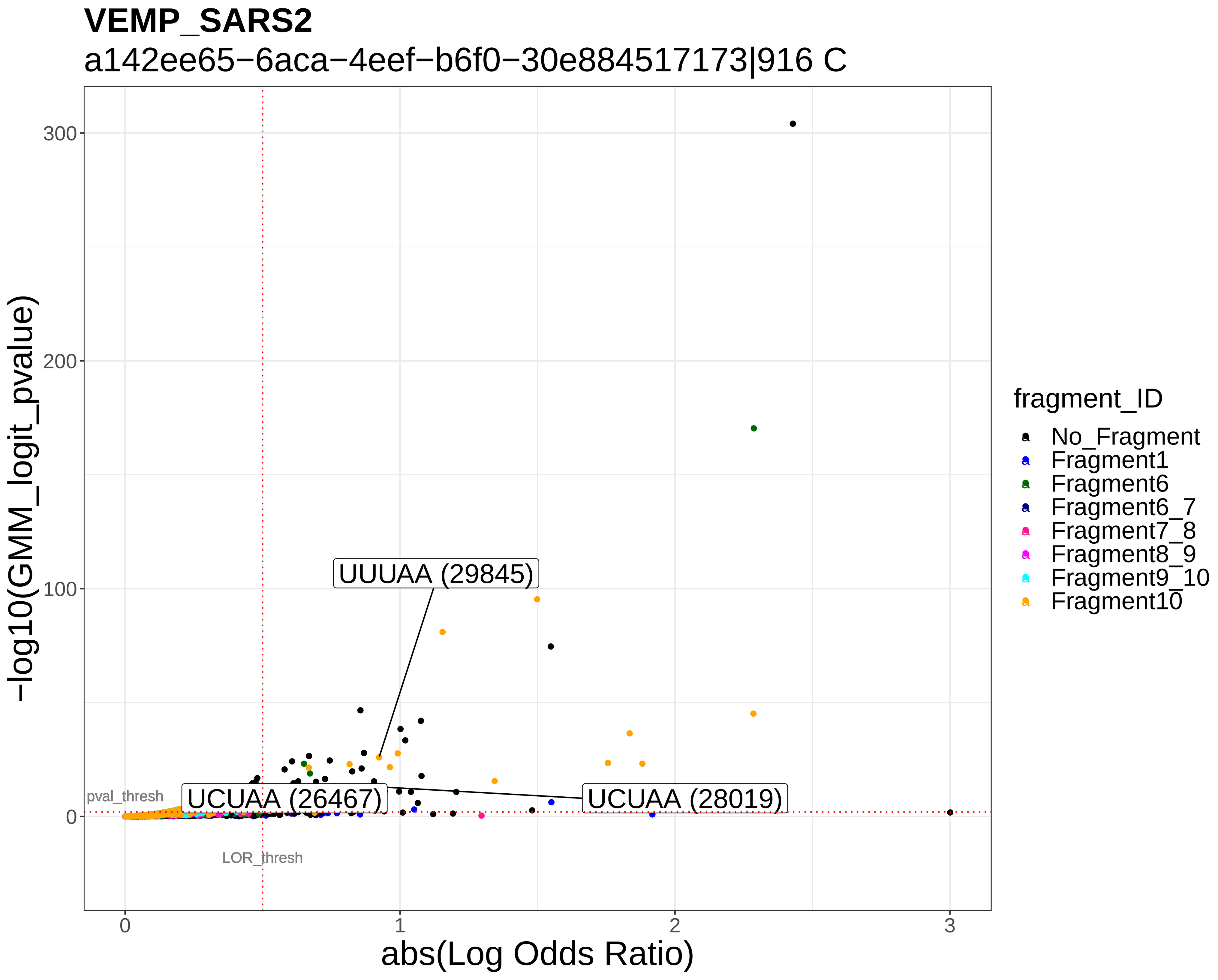
VEMP_SARS2
a142ee65−6aca−4eef−b6f0−30e884517173|916 C
−log10(GMM_logit_pvalue)
abs(Log Odds Ratio)
UUUAA (29845)
UCUAA (26467)
UCUAA (28019)
pval_thresh
LOR_thresh
fragment_ID
No_Fragment
Fragment1
Fragment6
Fragment6_7
Fragment7_8
Fragment8_9
Fragment9_10
Fragment10

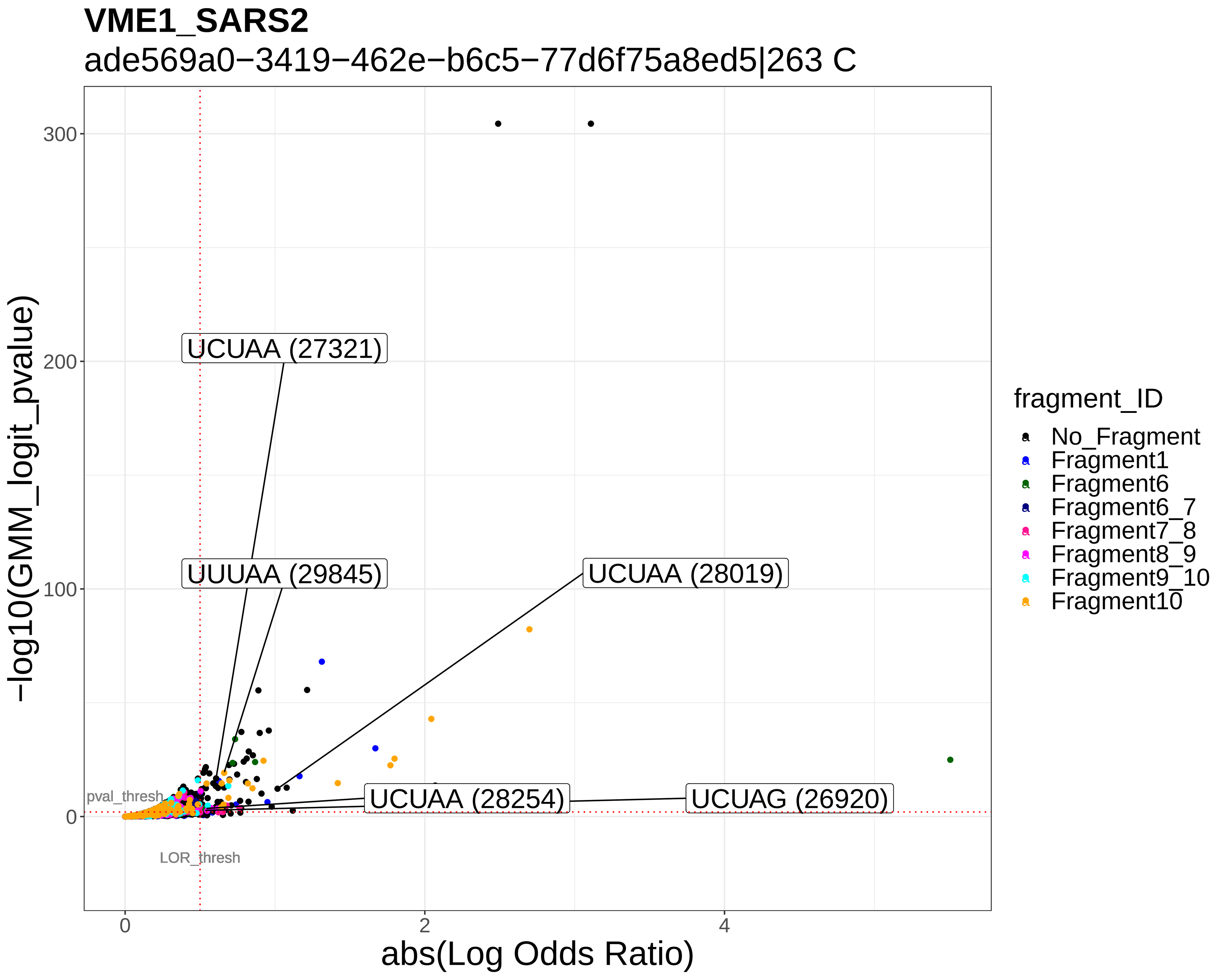
VME1_SARS2
ade569a0−3419−462e−b6c5−77d6f75a8ed5|263 C
−log10(GMM_logit_pvalue)
abs(Log Odds Ratio)
UCUAA (27321)
UUUAA (29845)
UCUAA (28019)
UCUAA (28254)
UCUAG (26920)
pval_thresh
LOR_thresh
fragment_ID
No_Fragment
Fragment1
Fragment6
Fragment6_7
Fragment7_8
Fragment8_9
Fragment9_10
Fragment10

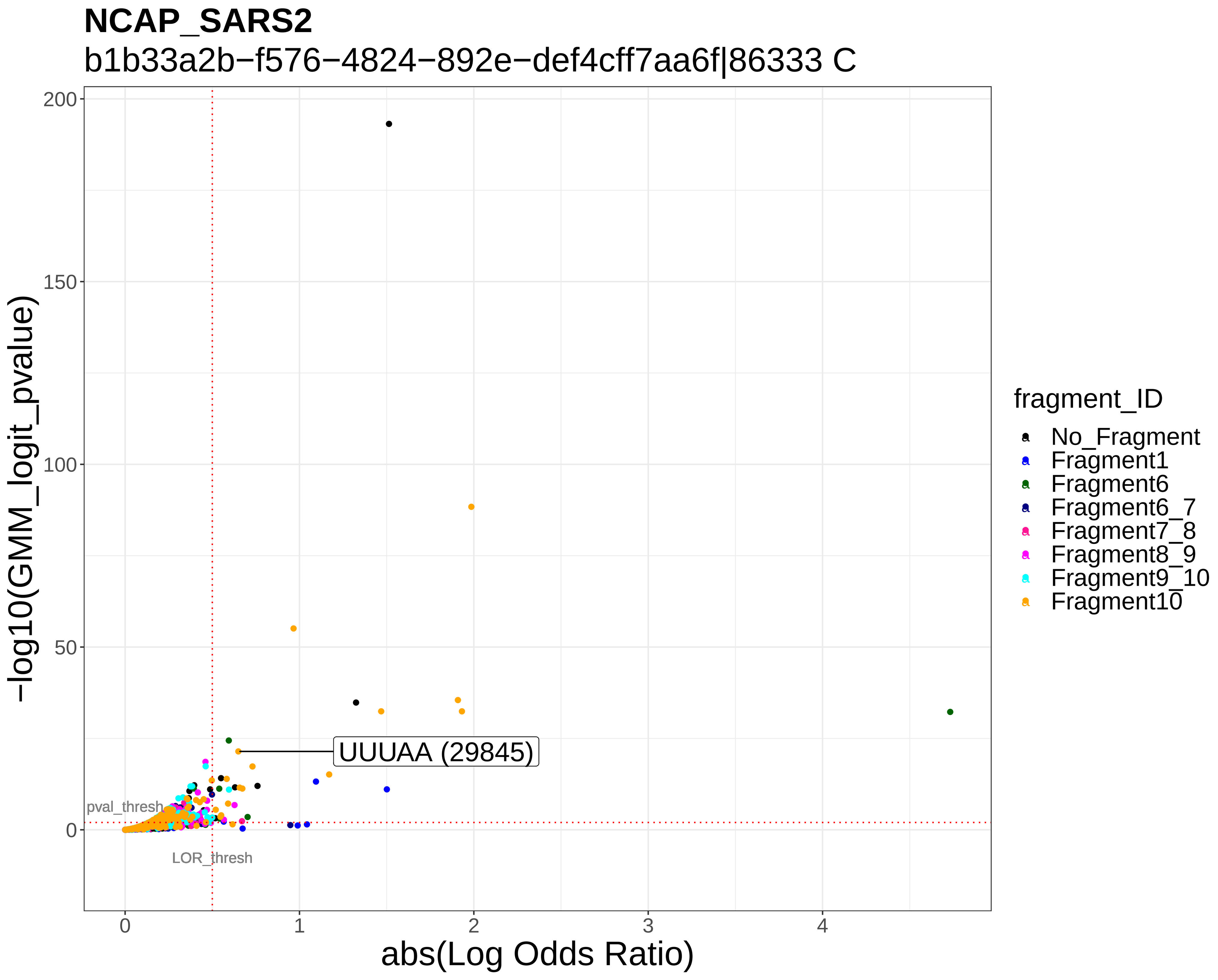
NCAP_SARS2
b1b33a2b−f576−4824−892e−def4cff7aa6f|86333 C
−log10(GMM_logit_pvalue)
abs(Log Odds Ratio)
UUUAA (29845)
pval_thresh
LOR_thresh
fragment_ID
No_Fragment
Fragment1
Fragment6
Fragment6_7
Fragment7_8
Fragment8_9
Fragment9_10
Fragment10

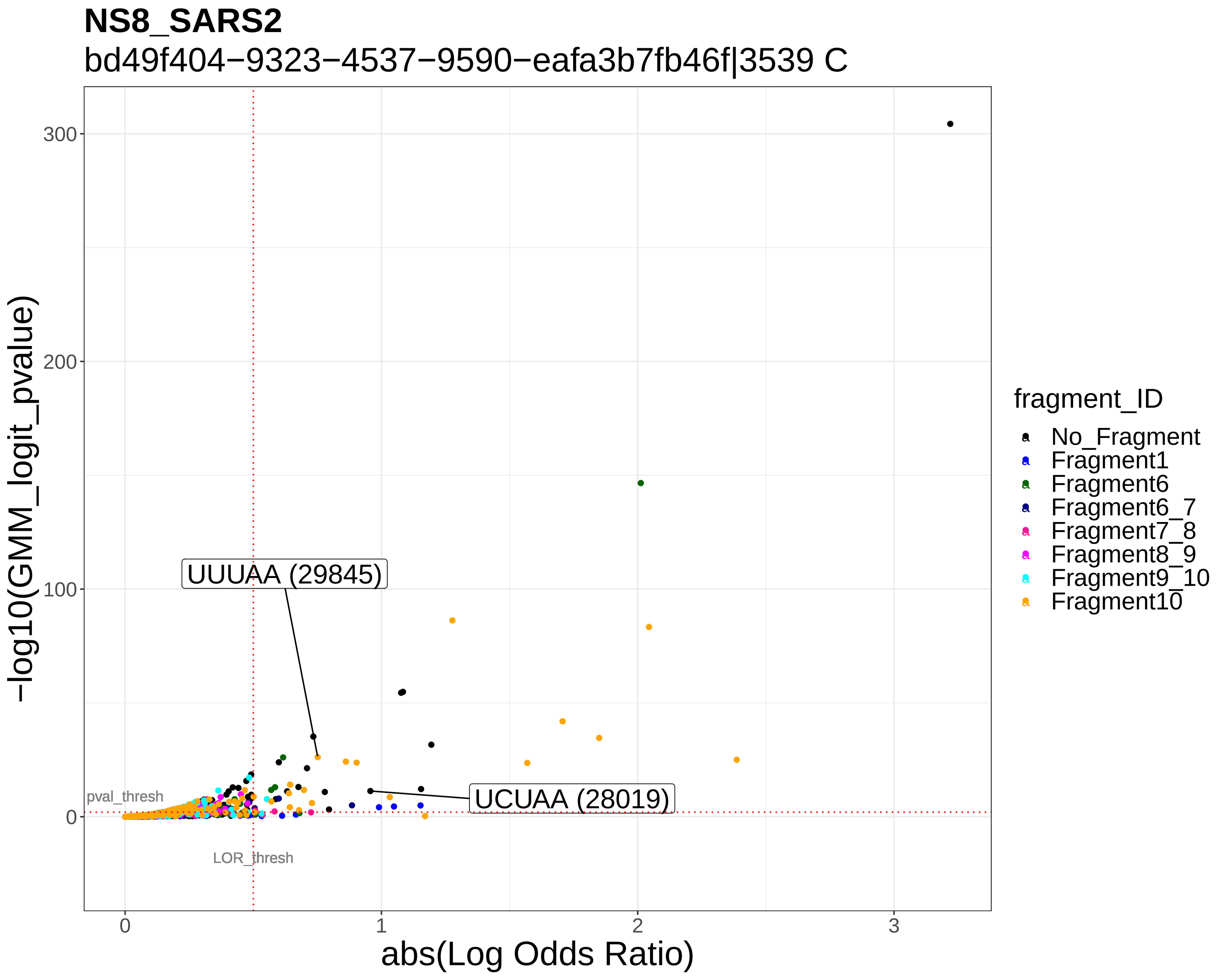
NS8_SARS2
bd49f404−9323−4537−9590−eafa3b7fb46f|3539 C
−log10(GMM_logit_pvalue)
abs(Log Odds Ratio)
UUUAA (29845)
UCUAA (28019)
pval_thresh
LOR_thresh
fragment_ID
No_Fragment
Fragment1
Fragment6
Fragment6_7
Fragment7_8
Fragment8_9
Fragment9_10
Fragment10

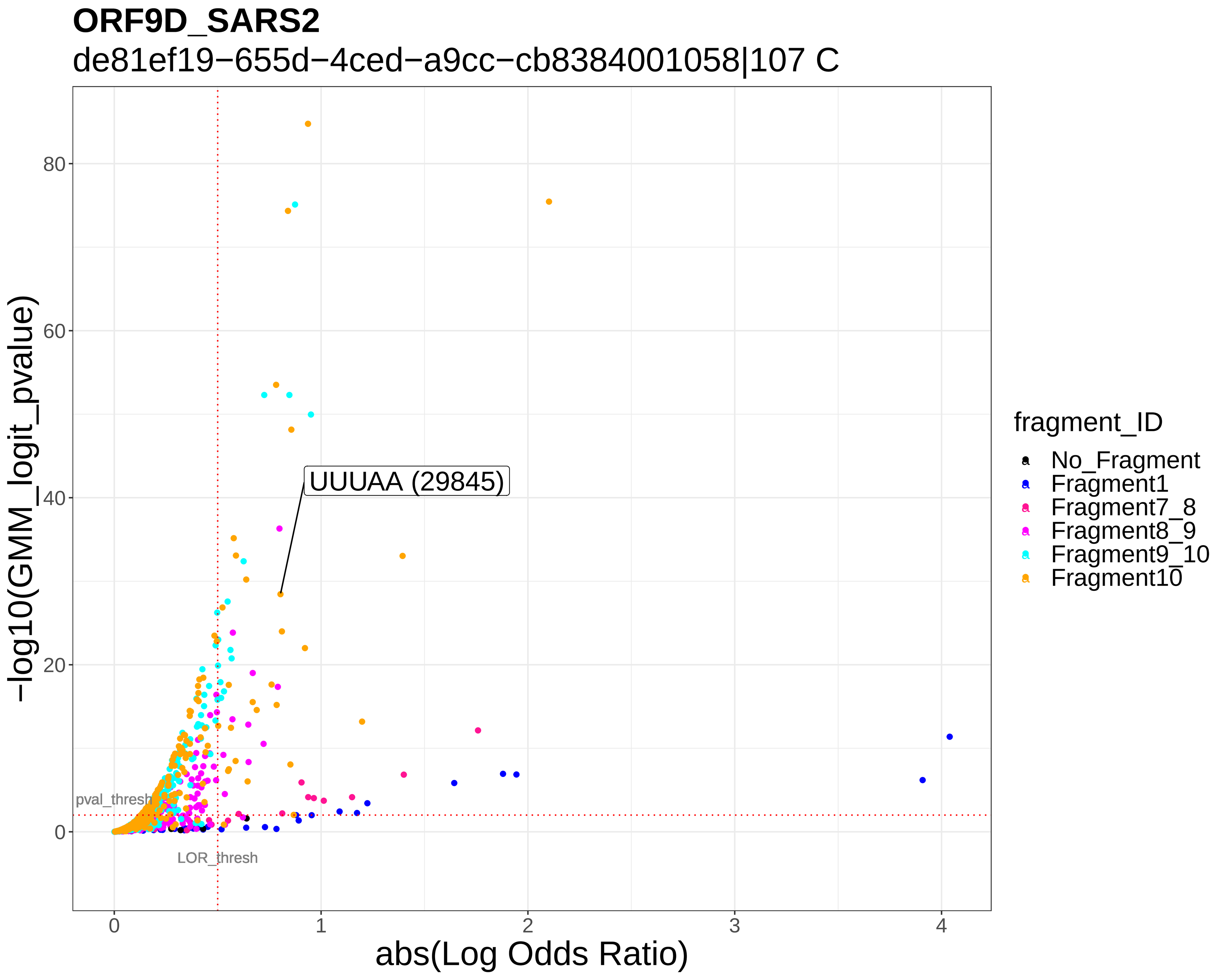
ORF9D_SARS2
de81ef19−655d−4ced−a9cc−cb8384001058|107 C
−log10(GMM_logit_pvalue)
abs(Log Odds Ratio)
UUUAA (29845)
pval_thresh
LOR_thresh
fragment_ID
No_Fragment
Fragment1
Fragment7_8
Fragment8_9
Fragment9_10
Fragment10

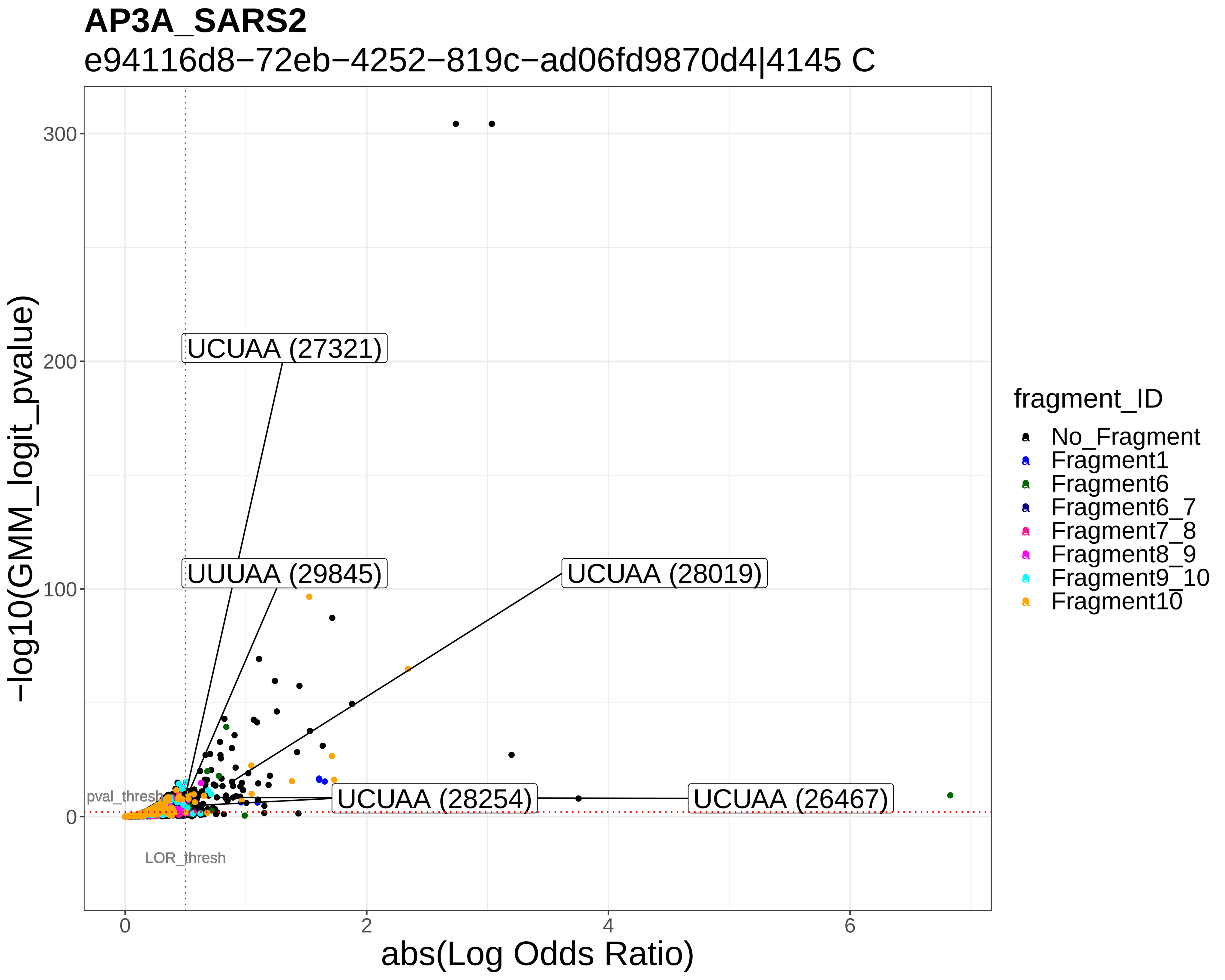
AP3A_SARS2
e94116d8−72eb−4252−819c−ad06fd9870d4|4145 C
−log10(GMM_logit_pvalue)
abs(Log Odds Ratio)
UCUAA (27321)
UUUAA (29845)
UCUAA (28019)
UCUAA (28254)
UCUAA (26467)
pval_thresh
LOR_thresh
fragment_ID
No_Fragment
Fragment1
Fragment6
Fragment6_7
Fragment7_8
Fragment8_9
Fragment9_10
Fragment10

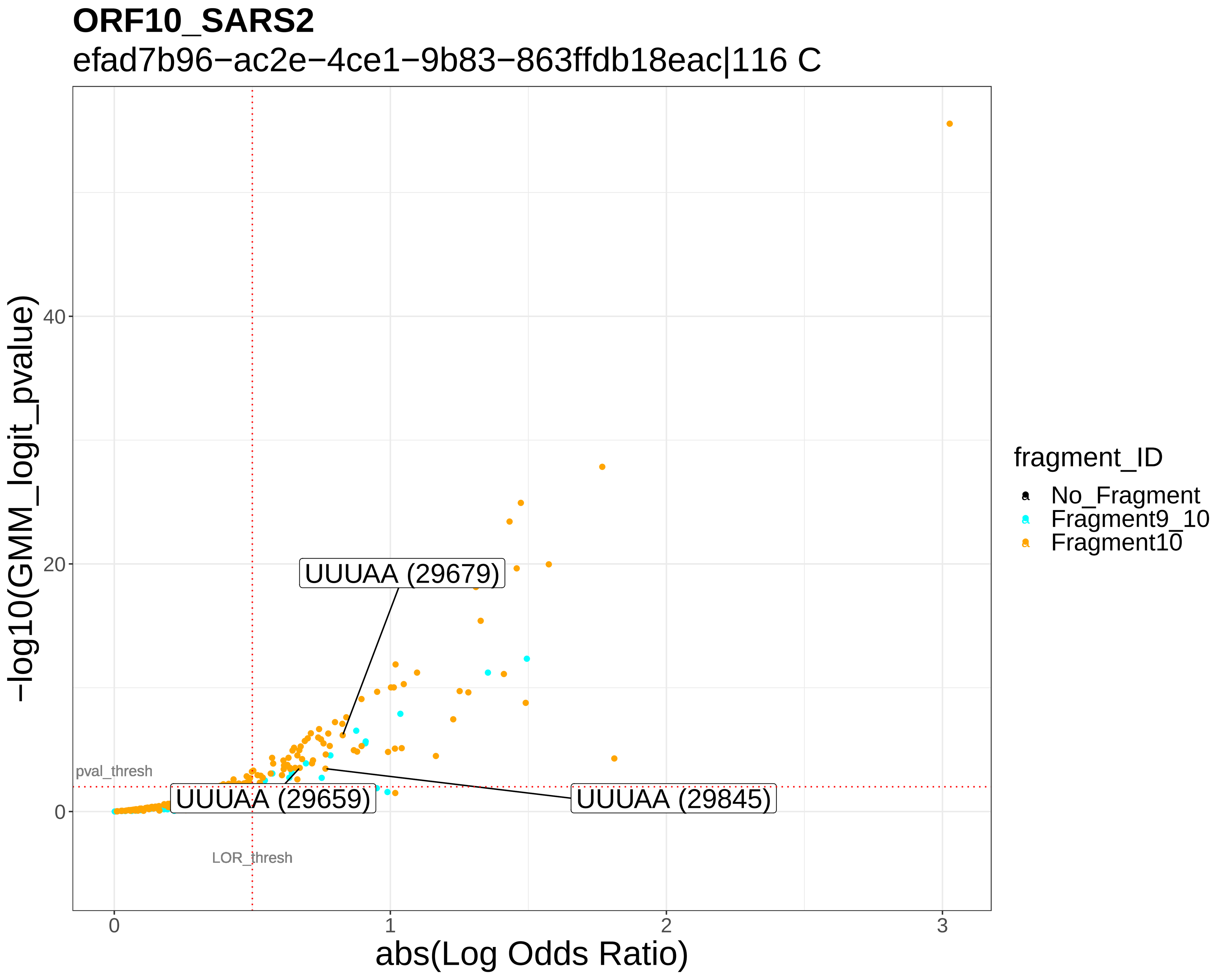

ORF10_SARS2
efad7b96−ac2e−4ce1−9b83−863ffdb18eac|116 C
−log10(GMM_logit_pvalue)
abs(Log Odds Ratio)
fragment_ID
No_Fragment
Fragment9_10
Fragment10
UUUAA (29679)
UUUAA (29659)
UUUAA (29845)
pval_thresh
LOR_thresh


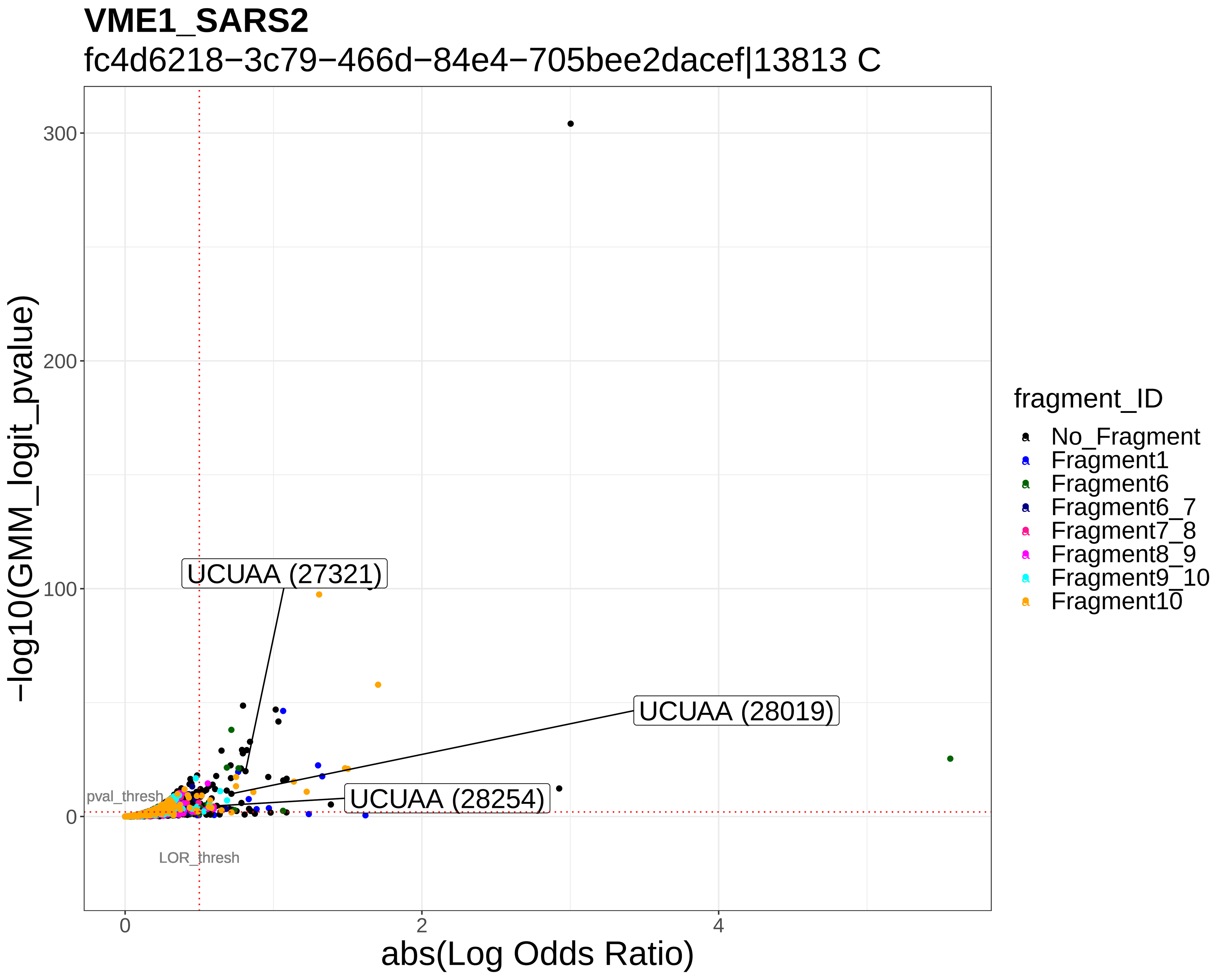

VME1_SARS2
fc4d6218−3c79−466d−84e4−705bee2dacef|13813 C
−log10(GMM_logit_pvalue)
abs(Log Odds Ratio)
fragment_ID
No_Fragment
Fragment1
Fragment6
Fragment6_7
Fragment7_8
Fragment8_9
Fragment9_10
Fragment10
UCUAA (27321)
UCUAA (28019)
UCUAA (28254)
pval_thresh
LOR_thresh

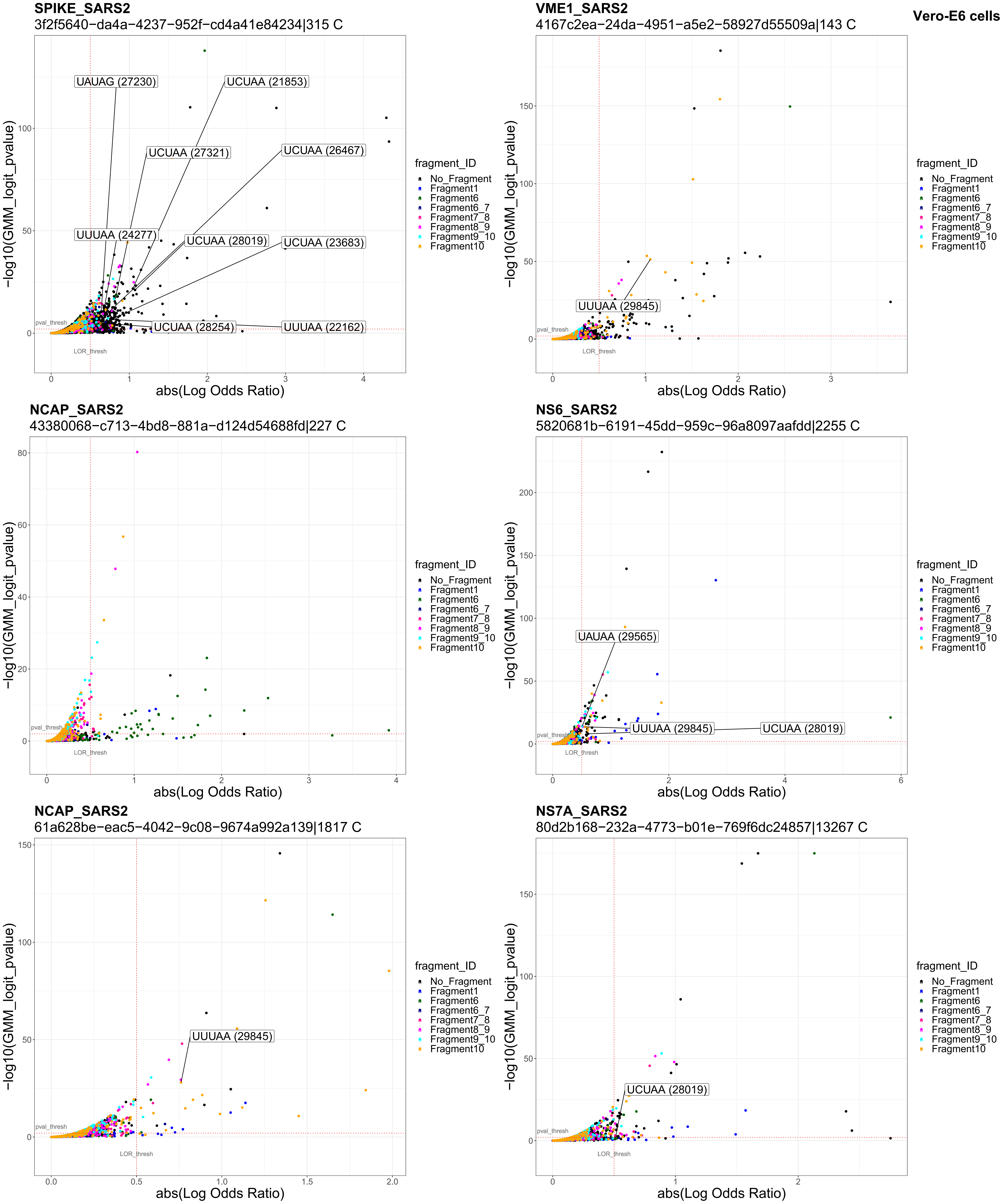
Vero-E6 cells
SPIKE_SARS2
3f2f5640−da4a−4237−952f−cd4a41e84234|315 C
UAUAG (27230)
UCUAA (21853)
UCUAA (27321)
UCUAA (26467)
UUUAA (24277)
UCUAA (28019)
UCUAA (23683)
UCUAA (28254)
UUUAA (22162)
pval_thresh
LOR_thresh
−log10(GMM_logit_pvalue)
abs(Log Odds Ratio)
fragment_ID
No_Fragment
Fragment1
Fragment6
Fragment6_7
Fragment7_8
Fragment8_9
Fragment9_10
Fragment10
VME1_SARS2
4167c2ea−24da−4951−a5e2−58927d55509a|143 C
UUUAA (29845)
NCAP_SARS2
43380068−c713−4bd8−881a−d124d54688fd|227 C
NS6_SARS2
5820681b−6191−45dd−959c−96a8097aafdd|2255 C
UAUAA (29565)
UUUAA (29845)
UCUAA (28019)
NCAP_SARS2
61a628be−eac5−4042−9c08−9674a992a139|1817 C
UUUAA (29845)
NS7A_SARS2
80d2b168−232a−4773−b01e−769f6dc24857|13267 C
UCUAA (28019)

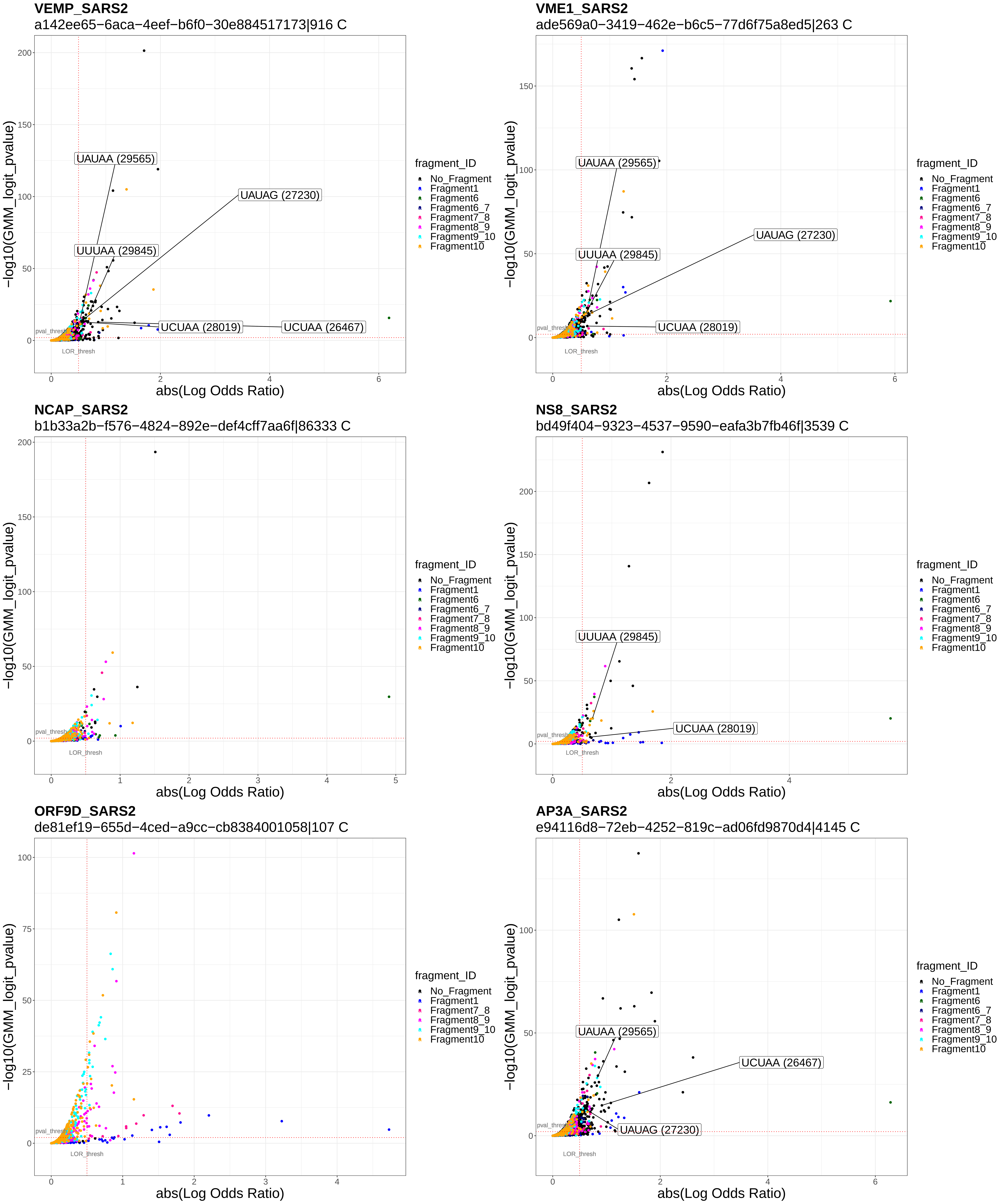
VEMP_SARS2
a142ee65−6aca−4eef−b6f0−30e884517173|916 C
UAUAA (29565)
UAUAG (27230)
UUUAA (29845)
UCUAA (28019)
UCUAA (26467)
pval_thresh
LOR_thresh
−log10(GMM_logit_pvalue)
abs(Log Odds Ratio)
VME1_SARS2
ade569a0−3419−462e−b6c5−77d6f75a8ed5|263 C
UAUAA (29565)
UUUAA (29845)
UAUAG (27230)
UCUAA (28019)
NCAP_SARS2
b1b33a2b−f576−4824−892e−def4cff7aa6f|86333 C
NS8_SARS2
bd49f404−9323−4537−9590−eafa3b7fb46f|3539 C
UUUAA (29845)
UCUAA (28019)
ORF9D_SARS2
de81ef19−655d−4ced−a9cc−cb8384001058|107 C
AP3A_SARS2
e94116d8−72eb−4252−819c−ad06fd9870d4|4145 C
UAUAA (29565)
UCUAA (26467)
UAUAG (27230)
fragment_ID
No_Fragment
Fragment1
Fragment6
Fragment6_7
Fragment7_8
Fragment8_9
Fragment9_10
Fragment10

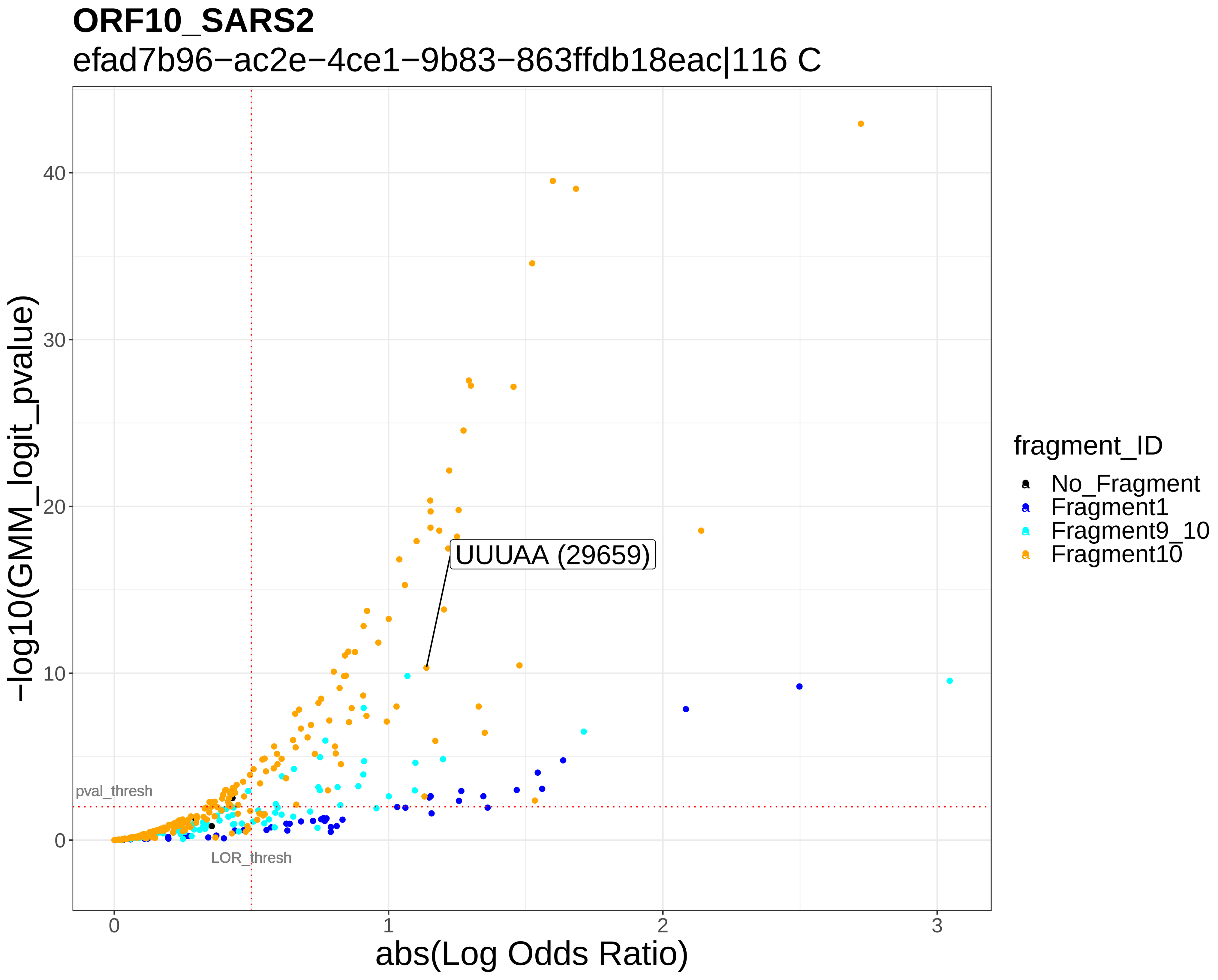

ORF10_SARS2
efad7b96−ac2e−4ce1−9b83−863ffdb18eac|116 C
−log10(GMM_logit_pvalue)
abs(Log Odds Ratio)
UUUAA (29659)
pval_thresh
LOR_thresh
fragment_ID
No_Fragment
Fragment1
Fragment9_10
Fragment10


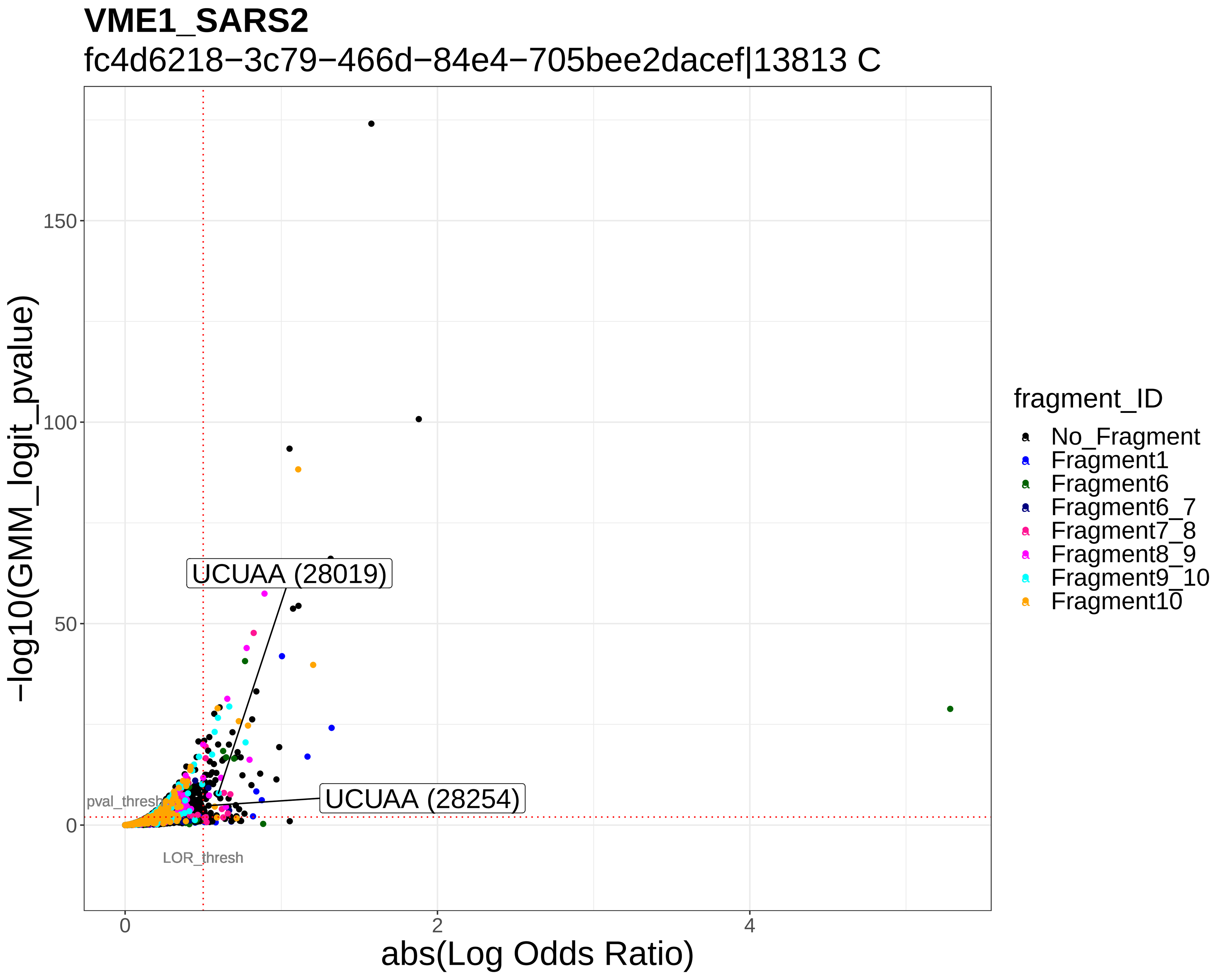

VME1_SARS2
fc4d6218−3c79−466d−84e4−705bee2dacef|13813 C
−log10(GMM_logit_pvalue)
abs(Log Odds Ratio)
UCUAA (28019)
UCUAA (28254)
pval_thresh
LOR_thresh
fragment_ID
No_Fragment
Fragment1
Fragment6
Fragment6_7
Fragment7_8
Fragment8_9
Fragment9_10
Fragment10
