## Supplementary figure legends and methods for "Discovering host protein interactions specific for SARS-CoV-2 RNA genome"

**Supplementary methods**

### *In-gel* digestion

Eluted biotinylated proteins were processed as previously described (https://doi.org/10.1038/nprot.2006.468). Briefly, proteins were initially separated on a precast 4-12% gradient gel (NP0322BOX, ThermoFisher Scientific). Each lane was divided in 6 slices that were cut from gels and destained in 50% v/v acetonitrile (ACN)/50 mM $NH_4HCO_3$. A reduction step was performed with 10 mM DTT, followed by alkylation with 55 mM iodoacetamide in the dark. After each step, samples were dehydrated with 100% ethanol and quickly dried in a centrifugal evaporator (SpeedVac). Subsequently, gel pieces were washed with 50 mM $NH_4HCO_3$ and overnight digested with 12.5 ng/ml trypsin (Promega, V5113) at 37 °C. The following day, tryptic digested peptides were extracted with Extraction Buffer (3% TFA, 30% ACN) and 100% ACN. Prior to MS, peptides were desalted and concentrated in a single step through reversed phase chromatography on micro-column C18 Stage Tips (https://doi.org/10.1038/nprot.2007.261 ) and eluted in 0.1% formic acid (FA).

### Nano-LC-MS/MS analysis

Peptide mixtures were analyzed by online nano-flow liquid chromatography tandem mass spectrometry using an EASY-nLC 1000 (Thermo Fisher Scientific, Odense, Denmark) connected to a Q-Exactive Plus instrument (Thermo Fisher Scientific) through a nano-electrospray ion source. The nano-LC system was operated in one column set-up with a 50-cm analytical column (75 µm inner diameter, 350 µm outer diameter) packed with C18 resin (EasySpray PEPMAP RSLC C18 2 µm 50 cm x 75 µm, Thermo Fisher Scientific) configuration. Solvent A was 0.1% FA in water and solvent B was 0.1% FA in 80% ACN. Samples were injected in an aqueous 0.1% TFA solution at a flow rate of 500 nL/min and separated with a gradient of 5%-40% solvent B over 50 min followed by a gradient of 40%–60% for 10 min and 60%-80% over 5 min at a flow rate of 250 nL/min in the EASY-nLC 1000 system. The Q-Exactive was operated in the data-dependent mode (DDA) to automatically switch between full scan MS and MSMS acquisition. Survey full scan MS spectra (from m/z 300-1150) were analyzed in the Orbitrap detector with resolution R = 35,000 at m/z 400. The ten most intense peptide ions with charge states R 2 were sequentially isolated to a target value of 3e6 and fragmented by Higher Energy Collision Dissociation (HCD) with a normalized collision energy setting of 25%. The maximum allowed ion accumulation times were 20 ms for full scans and 50 ms for MSMS and the target value for MSMS was set to 1e6. The dynamic exclusion time was set to 20 s.

### *In vitro* RNA binding assay

For the biolayer interferometry (BLI) assays, the two SL2 sequences with and without pseudouridine were purchased from Tebu-bio and Eurofins respectively, with a biotin molecule conjugated to the 5' end of the RNA. The experiments were performed at 25 °C on an Octet Red (ForteBio) instrument. RNAs and recombinant NSP1 protein (Biotechne, 10666-CV-050) were prepared in a buffer containing 25 mM HEPES pH 7.5, 150 mM KCl, 1 µM β-mercaptoethanol and 0.05% Tween-20. The biotinylated RNA sequences were diluted to a concentration of 1 µg/ml and loaded on streptavidin-coated biosensors. The protein was prepared at 5 µM and serially diluted 8 times. The binding constants ($K_d$) were estimated by fitting the response intensity as a function of the protein concentration. The experiment was performed in triplicates.

### UV-RIP assay

UV-RIP protocol was modified from https://doi.org/10.1016/j.cell.2011.06.026 and https://doi.org/10.1016/j.cell.2012.03.035. Briefly, HEK293T were seeded in 10 cm dishes and transfected with the indicated mammalian expression plasmids were harvested, washed once with PBS and UV-crosslinked on ice with 2 cycles of irradiations at 100000 µJ/cm2. Cells were lysed with Lysis Buffer (0.5% NP-40, 0.5% NaDeoxycholate, 1x Roche protease inhibitors mixture (04693116001 mercK), 25 U/mL RNAse inhibitor (M03070L, NEB) in PBS and rocked on a wheel for 30 min at 4°C. Afterwards, lysates were treated with 30 U of Turbo DNAseI (AM2239, ThermoFisher Scientific) for 30 min at 37 °C in a Thermomixer rotating at 1100 rpm and subsequently centrifuged for 5 min at 1350 g to remove cellular debris. The supernatants were used for the immuno-purification experiment, with an aliquot (10%) kept as input material. Half of the supernatant volume (45%) was incubated with 4ug of HA antibody (901502, BioLegend) in a final volume of 500 uL with additional Lysis Buffer and rocked overnight at 4°C. The day after, 50 uL protein G dynabeads (10007D, ThermoFisher Scientific) were first washed 3 times in Lysis Buffer and then added to each sample. Samples were rocked for additional 3 hours at 4 °C. Afterwards, dynabeads were washed 4 times with Washing Buffer I (PBS supplemented with 1% NP-40, 0.5% NaDeoxycholate, 300 mM NaCl, 1x Roche protease inhibitor mix and 25 U/mL RNAse inhibitor). The dynabeads were then resuspended in 100 μL RNAse-free water and treated again with the Turbo DNAseI for 30 min at 37 °C, in a Thermomixer at 1100 rpm. Input material was also treated with DNAseI for a second time. The dynabeads were then washed 4 times with Washing Buffer II (PBS supplemented with 1% NP-40, 0.5% NaDeoxycholate, 300 mM NaCl, 10 mM EDTA, 1xRoche protease inhibitor and 25 U/mL RNAse inhibitor). Finally, the RNA was eluted from the beads using the RNA-Lysis Buffer (Zymo Research), and extracted through the RNA-extraction kit (Zymo Research). The RNA was retro-transcribed into cDNA using the ImProm-IITM (A3800, Promega), according to the vendor's instruction. The cDNA was then diluted 1:5 with water and 5% of the diluted material was analyzed by RT-qPCR analysis using the Fast SYBR green master mix (4385614, ThermoFisher Scientific). The values obtained for each immunoprecipitated RNAs were normalized over the respective input material and plotted in a histogram, as relative fold enrichment.

**Nanocompore cell line-specific analysis**

The following analysis was performed on Nanocompore output for each cell line. K-mers were considered significant if they had a GMM_logit_p-Value ≤ 0.01, an absolute value of LOR ≥ 0.5, and one of the 5 nucleotides was a uridine. Significant k-mers were removed if they were within 15 nt of a TRS-L/B junctions (ORF junctions) or 25 nt of an IVT fragment boundary (IVT junctions) (https://doi.org/10.1016/j.cell.2020.04.011), because these regions are prone to aberrant signal artifacts that can cause false positives.

A single modification can affect multiple k-mers, thus we used the find_peaks function from scipy.signal (https://doi.org/10.1038/s41592-019-0686-2) to merge neighboring significant sites into unified peaks. The $-\log_{10}$ (p-values) of U-containing k-mers were used for peak calling with a threshold of 2 and a width of 5. Any of these k-mers that were within 15 nt of an TRS-L/TRS-B junction or 25 nt of an IVT junction were also removed as described above. Only k-mers which aligned to reference sgRNAs with the TRS-L/B junction next to ORF9d, ORF10 and canonical reference sgRNAs were included because early-terminating non-canonical sgRNAs were shown to be problematic to map correctly (https://doi.org/10.1093/nar/gkac144).

The resulting sites were compared to those identified in Fleming et al. (https://doi.org/10.1021/acscentsci.1c00788) by creating an array with genomic positions of the five nucleotides for every significant k-mer. The R function “%in%” was used to compute the intersection between each of these arrays and an array containing the genomic positions of Fleming et al. “high-confidence sites”. Any k-mer with a nucleotide that overlapped one of the “high-confidence” sites was considered a match.

Data was visualized using the UCSC Genome Browser tracks from BED files of the peakcalled and filtered k-mers (https://doi.org/10.1101/gr.229102) (see **Data availability**).

**Secondary structure of SARS-CoV-2 TRS-L**

The secondary structure of SARS-CoV-2 TRS-L represented in **Fig. 5A** was obtained generating a minimum free energy prediction for the first 80 nucleotides of the genomic viral sequence using the RNAfold Webserver. This prediction was used to build a stockholm format (.sto) file which was then annotated through a custom script (see **Code Availability**) with the Nanocompore p-values for the reads mapping to the NS6 sgRNA in CaCo-2 infected cells. This sgRNA was chosen among the three sgRNAs bearing the UGUAR modified site in the TRS-L because it is the only one corresponding to a single canonical transcript (https://doi.org/10.1093/nar/gkac144). K-mers with p-value $\leq 0.01$ and abs (LOR) $\geq 0.5$ were annotated. The annotated .sto file was then plotted using r2r (https://doi.org/10.1186/1471-2105-12-3).

**Supplementary figure legends:**

**Supplementary Figure 1.** Predicted RNA secondary structures of the selected BoxB-RNA fragments. The displayed RNAs represent the minimum free energy structures obtained using the ViennaRNA Web services.

**Supplementary Figure 2.** FACS analysis of cells co-transfected with the BoxB-RNA vector expressing also the GFP protein and the λN-HA-BASU vector expressing the RFP protein. The scatter plots showed the percentage of cells expressing the GFP, the RFP or both proteins. Untransfected cells (mock) or cells transfected with either λN-HA-BASU vector or BoxB-RNA vector were used as control to distinguish the GFP and RFP positive cells.

**Supplementary Figure 3. A)** Western Blot (WB) analysis of HEK293T whole cell extract (W.C.E.). Cells were transfected with plasmids expressing the λN-HA-BASU and one of the BoxB-RNA fragments. Biotin was administered into the medium 1h before cell harvesting. For WB analysis, 20 ug of each sample was loaded on a 4-12% pre-cast gel. The displayed data is the representative image of one of the three biological experiments performed. **B)** Streptavidin pulldown followed by WB analysis of the samples described in **A)**. For all samples, 3% of the eluted material (E) was loaded on a 4-12% pre-cast gel and stained with HRP-conjugated streptavidin (STREP-HRP). To control the efficiency of the streptavidin pulldown, 5% of input (I) and the relative flowthrough (F) material of the Scramble RNA was also loaded in the gel. The displayed data is the representative image of one of the three biological experiments performed.

**Supplementary Figure 4. A)** Number of total interactors identified for each RNA fragment (labeled with the letter “F” follow by a number) and definition of the interaction partners in common with the control “Scramble” (green) compared to the protein identified uniquely with SARS-CoV-2 RNA (pink). **B)** Volcano plots of the RaPID-MS analysis of each fragment against the Scramble control. Proteins statistically significant according to the student T-test ($P < 0.05$) are displayed in the figures. Volcano plots of the RaPID-MS analysis of each fragment against the Scramble control. Proteins statistically significant according to the student T-test ($P < 0.05$) are displayed in the figures.

**Supplementary Figure 5. A)** Venn diagram showing the overlap of the generated RaPID-MS dataset and the previously published dataset on the interactions between SARS-CoV-2 RNA and the host proteins in human cells. **B)** List of the common interactors of RaPID-MS dataset with at least one of the published datasets. **C)** Gene Ontology (GO) analysis performed on the

73 proteins specifically associated with the 10 SARS-CoV-2 RNA fragments and identified by RaPID-MS.

**Supplementary Figure 6. A)** Scatter chart of the analysed RBPs ranked according to their relative $\log_2$ *cat*RAPID score, displaying each RNA fragment at time. **B)** Boxplot representation of the distribution of the predicted interactions according to the log2 *cat*RAPID score.

**Supplementary Figure 7. A)**. Nanocompore analysis identified 63 significant k-mers over the SARS-CoV-2 gRNAs. The figure shows a UCSC Genome Browser (*http://genome.ucsc.edu*) annotation of the NRCeq reference assembly, the full-length viral genome, the modified k-mers across the whole reference genome, the modified k-mers common to the list of gRNA and sgRNAs sites. UniProt Protein Products are present as a reference. **B)** Nanocompore plot for sites found in the gRNAs analysis. Thresholds are displayed as red dashed lines. Every displayed point represents a k-mer identified as significant compared to the IVT sample. The position is referred to the 0-based first nucleotide of the k-mer. The color code represents k-mers overlapping with RaPID fragments. **C)** Venn diagram of the significant U-containg sites identified in SARS-CoV-2 gRNA and sgRNAs.

**Supplementary Figure 8.** Nanocompore plots for each reference canonical sgRNA in every cell line. On the y-axis is displayed the GMM p-value, while on the x-axis the log odds ratio (LOR). Thresholds are displayed through red dashed lines. Every point represents a k-mer identified as significant compared to the IVT sample. Of these significant k-mers, those containing the UNUAR motif have been labeled. The position is referred to the 0-based first nucleotide of the k-mer. The color code represents k-mers overlapping with RaPID fragments.

**Supplementary Figure 9.** Distributions of dwell time and median intensity per position for the eight significant sites, shared between at least two cell lines and containing the UGUAR motif. Each position displays distributions for two conditions, the reference (in this case the IVT) and the test condition.

**Supplementary Figure 10. A)** Investigation of the secondary structure analyzed by circular dichroism. The higher intensity of the maximum at 270 nm indicates an increased structural content. **B)** Normalized binding curves for NSP1 with SL2 containing the consensus sequence “UGUAR” or “UGψAR”, determined by biolayer interferometry. The calculated dissociation constants ($K_d$) are 300+/-54 nM for the non-modified sequence and 170+/-28 nM for the pseudouridylated one.

**Supplementary Table 1:** DNA sequences of the SARS-CoV-2 fragments used in this study.

**Supplementary Table 2:** Library of 2064 human RNA-binding proteins used in the analyses.

**Supplementary Table 3:** List of the proteins identified in association with each fragment by RaPID-MS analysis. The protein interactors for each fragment are reported in an independent spreadsheet named with the respective SARS-CoV-2 fragment. The spreadsheet named “Summary” displayed a summary of only the significant interactors identified, according to the T-Test analysis. In addition, it was indicated if proteins were annotated as RNA binding proteins (RBPs) and if they were interacting with only one of the 10 RNA fragments. The spreadsheet named “Overlap published datasets” displayed all the identified proteins and the significant proteins identified in the RaPID-MS dataset and in the published datasets.

**Supplementary Table 4:** *cat*RAPID performances on RAPID-RBP dataset. For each protein-RNA pair, the *cat*RAPID predictive score, the median LFQ value, the abundance and length of the protein and the normalized LFQ values are shown. Both the overall interactome and the single fragments interactomes are available. The list of proteins with a $\log_2$ *cat*RAPID score higher than 5.22 (85th percentile) removing those enriched with Scramble RNA is provided in the datasheet named “85th percentile no scramble”. The datasheet named “statistics” contains the Chi squared test applied on the distribution of the *cat*RAPID score of each RNA-protein interaction for each SARS-CoV-2 RNA fragment.

**Supplementary Table 5**: Modification sites in SARS-CoV-2 gRNAs in infected CaCo-2, CaLu-3 and Vero E6 cells.

- Sheet *all_sites*: Nanocompore results including significant and non-significant k-mers.
- Sheet *significant_sites*: All k-mers passing LOR and GMM p-value thresholds (k-mers overlapping IVT or SNPs are excluded).

- Sheet *Us_sign_sites*: All uridine-containing k-mers passing LOR and GMM p-value thresholds (k-mers overlapping IVT or SNPs are excluded).
- Sheet sign_U_sites_common_to_sgRNAs: All uridine-containing k-mers passing LOR and GMM p-value thresholds (k-mers overlapping IVT or SNPs are excluded) common to at least one of the significant uridine-containing sites present in at least one of the canonical transcripts in the pulled analysis of sgRNAs sites (see **Supplementary Table 10**).
- Sheet *SNPs*: all SNPs for the different viral strains used to infect CaCo-2, CaLu-3 and Vero E6 cells.

**Supplementary Table 6, 7, 8, 10**: Modification sites in SARS-CoV-2 sgRNAs in infected CaCo-2, CaLu-3 and Vero E6 cells.

- Sheet *samples* - List of samples for the cell line of interest and analysis parameters.
- Sheet *all_significant_sites* - All k-mers passing LOR and GMM p-value thresholds (k-mers overlapping IVT or ORF junctions are not excluded).
- Sheet *all_canonical_Us_sign_sites* - K-mers passing LOR and GMM p-value thresholds with at least one uridine nucleotide and belonging to a canonical sgRNA (k-mers overlapping IVT or ORF junctions are excluded).
- Sheet *5p_significant_sites* - K-mers passing LOR and GMM p-value thresholds located before genomic position 100 (k-mers overlapping IVT or ORF junctions are not excluded).
- Sheet *5p_canonical_Us_sign_sites* - K-mers passing LOR and GMM p-value thresholds with at least one uridine nucleotide, belonging to a canonical sgRNA and located before genomic position 100 (k-mers overlapping IVT or ORF junctions are not excluded).
- Sheet *Fleming_redundant_all_sites* - All k-mers overlapping one of the Fleming et al. sites according to our analysis (k-mers overlapping IVT or ORF junctions are not excluded).
- Sheet *Fleming_redundant_sign_sites* - K-mers passing LOR and GMM p-value thresholds overlapping one of the Fleming et al. sites according to our analysis (k-mers overlapping IVT or ORF junctions are not excluded).
- Sheet *Fleming_non_red_CandNC_sites* - Non-redundant k-mers passing LOR and GMM p-value thresholds overlapping one of the Fleming et al. sites according to our analysis (k-mers overlapping IVT or ORF junctions are not excluded).

- Sheet *Fleming_non_red_CandN_all_sites* - Non-redundant k-mers overlapping one of the Fleming et al. sites according to our analysis (k-mers overlapping IVT or ORF junctions are not excluded).
- Sheet *peakcalled_canonical_Us* - K-mers passing LOR and GMM p-value thresholds with at least one uridine nucleotide, peakcalled as in Leger et al.(36) and belonging to a canonical sgRNA (k-mers overlapping IVT or ORF junctions are excluded).
- Sheet *Fleming_in_peakcalled_canonical* - K-mers passing LOR and GMM p-value thresholds with at least one uridine nucleotide, peakcalled as in Leger et al.(36), belonging to a canonical sgRNA and overlapping one of the Fleming et al. sites (k-mers overlapping IVT or ORF junctions are excluded).

**Supplementary Table 9**: Modification sites shared between the different cell lines.

- Sheet *shared_2_U* - modified sites shared between at least two cell lines.
- Sheet *shared_3_U* - modified sites shared between all the three cell lines.
